# Neural circuit models for evidence accumulation through choice-selective sequences

**DOI:** 10.1101/2023.09.01.555612

**Authors:** Lindsey S. Brown, Jounhong Ryan Cho, Scott S. Bolkan, Edward H. Nieh, Manuel Schottdorf, David W. Tank, Carlos D. Brody, Ilana B. Witten, Mark S. Goldman

## Abstract

Decision making is traditionally thought to be mediated by neurons that accumulate evidence through persistent activity. However, recent decision-making experiments in rodents have observed neurons across the brain that fire sequentially, rather than persistently, with the subset of neurons in the sequence depending on the animal’s choice. We developed two new candidate circuit models in which neurons are active sequentially and transfer evidence faithfully to the next active population. One model encodes evidence in the relative firing of two competing chains of neurons, and the other in the network location of a stereotyped pattern (“bump”) of neural activity. Neural recordings from four brain regions during an evidence accumulation task revealed that different regions displayed evidence tuning consistent with different candidate models. This work provides a mechanistic explanation for how graded information may be precisely accumulated within and transferred between neural populations, and suggests that different brain regions may accumulate evidence through different circuit mechanisms.

## INTRODUCTION

The accumulation of evidence for different alternatives is thought to be a fundamental cognitive operation. As such, neural recordings have been performed in many tasks designed to probe how evidence is accumulated to make a decision, including tactile^1,2^, visual^3–11^, olfactory^12^, auditory^13,14^, motor^15^, and value discrimination tasks^16–20^. Experiments using these tasks have been conducted with the goal of understanding how the neural circuitry within and across different brain regions contributes to the accumulation of evidence, subsequent maintenance of this information in working memory, and ultimate commitment to a decision based on this evidence^21^.

A major class of models proposed to describe this decision making process is the drift diffusion model^22–24^, in which evidence is accumulated with a noisy drift process, and decisions are made based on when the resulting accumulation reaches some threshold. Many circuit-based models have been shown to be equivalent to the drift-diffusion model^25^. Such circuit-based models include mutual inhibition models^26^, feedforward inhibition models^27^, and pooled inhibition models^28^. These basic circuit models have been extended to explain a number of different decision-making experiments with recordings in different regions where neurons show persistent activity^29–34^.

The mechanistic essence of the circuit-based models is the accumulation of evidence along a low-dimensional attractor representing the decision variable. Thus, these models predict that the activity of single neurons ramps up or down with evidence throughout a decision-making task^35,36^. Such ramping activity during decision-making tasks has been observed in a number of different brain regions in different paradigms, including in the lateral intraparietal area (LIP), medial temporal area (MT), and ventral intraparietal area (VIP) of monkeys in a random dot motion discrimination task^37–40^, in the dorsal premotor cortex of monkeys during discrimination tasks^41,42^, and in the posterior parietal cortex (PPC) and frontal orienting fields (FOF) of rats in an auditory evidence accumulation task^14,43^. These classes of neural circuit models that accumulate evidence through persistent, ramping activity successfully capture much observed experimental data.

However, a mounting body of evidence from large scale recordings during a range of decision-making tasks suggests that, in many decision-making contexts, neurons across cortical and subcortical regions are not persistently active. Instead, neurons fire transiently and sequentially in a choice-specific manner. This occurs in decision-making tasks both with navigation^44–49^ and without navigation^50–58^, including in evidence accumulation tasks^44,49^. These data challenge existing models of evidence accumulation by contradicting a core tenet of previous models: the presence of persistent, rather than sequential, neural activity.

For a model to accumulate evidence through sequential neural firing, it must perform two key computations. First, as in traditional models based on persistent activity, it must accumulate evidence received at each position. Second, unlike traditional models, it must transfer information between neurons at sequential positions.

Here, we develop two classes of neural circuit models that perform these computations. In each model class, the transfer of accumulated evidence between neurons encoding sequential positions is accomplished via a position-modulated gating signal. The key difference in the models is in their encoding of evidence, which is either in the relative *amplitude* of activity in two competing chains of neurons or in the network *location* of a stereotypically shaped, unimodal pattern (“bump”) of neuronal firing. We tested these differentiating evidence tuning predictions in a dataset of 14,247 imaged neurons, collected across 4 cortical and subcortical regions in mice performing an evidence accumulation task in virtual reality^59^. This analysis revealed that while seemingly similar choice-selective sequences are present in this task across these regions, neurons in different regions differ in their evidence tuning properties. In the neocortex, neuronal tuning curves mainly exhibit a monotonically increasing or decreasing encoding of evidence, characteristic of the competing chains model class, whereas in the hippocampus neurons tend to have narrow, non-monotonic tuning curves, consistent with the bump model. Thus, our results reveal different encodings of accumulated evidence across brain regions and suggest that different regions could use distinct circuit mechanisms to form their evidence representations.

## RESULTS

### Traditional Integrator Models Fail to Explain Choice-Selective Sequences in the Accumulating Towers Task

The models presented below are motivated by a large dataset (n = 14,247 neurons; N=26 mice) of newly acquired (previously unpublished) calcium imaging data (see Methods) from anterior cingulate cortex (ACC) and dorsomedial striatum (DMS), as well as recently published imaging data from hippocampus (HPC)^49^ and retrosplenial cortex (RSC)^44^. These data were all obtained during the same navigation-based, accumulation of evidence task in virtual reality (“accumulating towers task”, Fig. 1A). In this task^59^ (see Methods), mice navigate a T-maze with Poisson counts of visual cues in the form of towers appearing to each side. After a delay region with no cues, the mouse reaches the arms of the maze and is rewarded for turning to the side with more cues (Fig. 1A). To make the correct decision, the mouse must integrate the number of cues and hold this information in working memory during the delay region. Mice are able to perform this task with good accuracy that increases with the magnitude of the difference between the number of cues to each side^59^.

**Figure 1.**
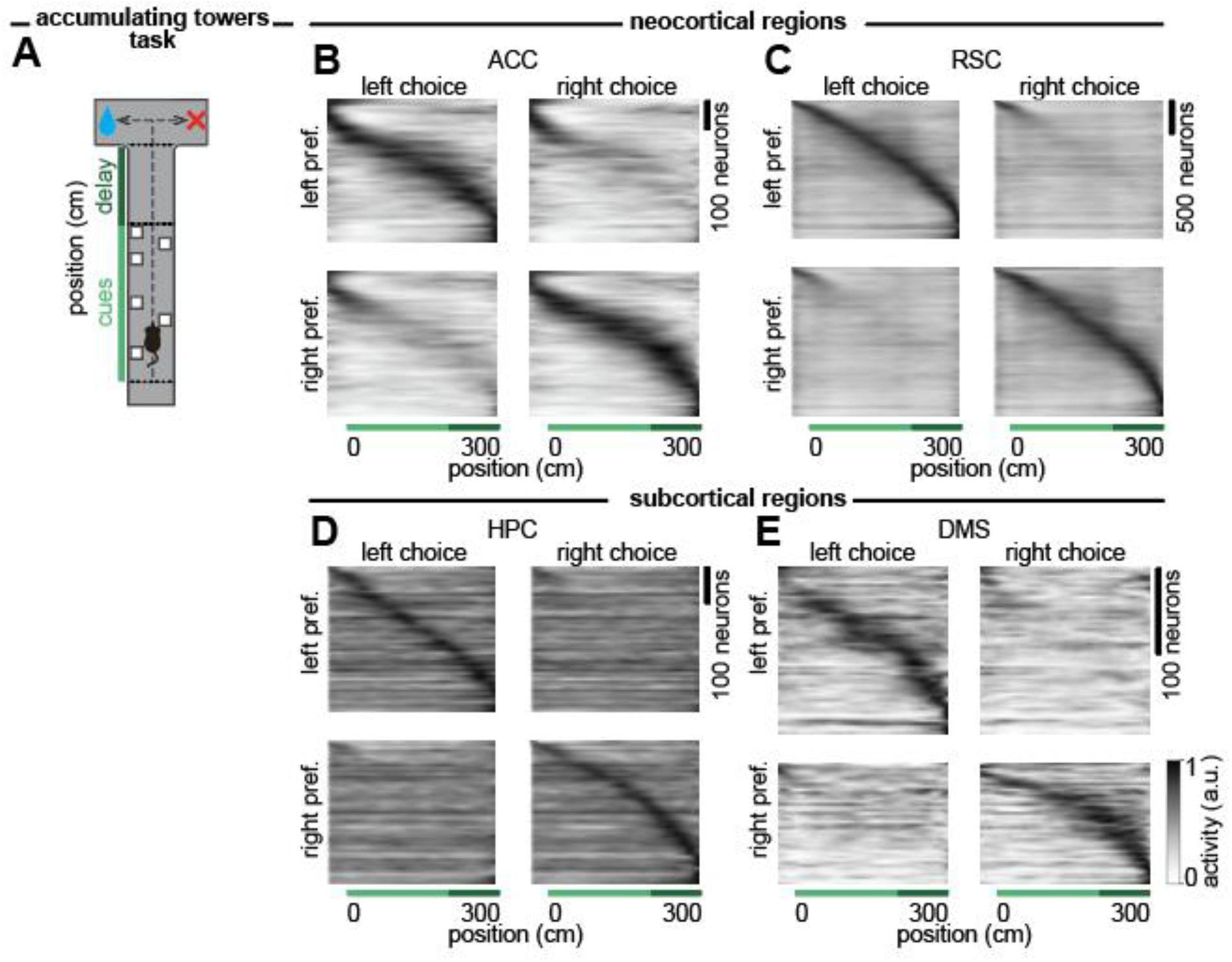
Choice-selective sequences are observed across brain regions in a navigation-based, accumulation-of-evidence task. **(A)** Schematic of a behavioral task in which mice navigate a T-maze in virtual reality. Visual cues (towers, white rectangles) are presented to either side of the maze during the cue period (light green, 200 cm long). At the end of the delay period (dark green, 100 cm long), mice are rewarded for turning to the side with more towers. **(B)** Each row shows the peak-normalized estimated firing rate at each position in the maze of a neuron with significant evidence tuning (see Methods) recorded during this task from anterior cingulate cortex (ACC, n = 964 neurons, see Methods), averaged across trials when the animal turned left (left choice, left column) or right (right choice, right column). Neurons were assigned as left or right choice preferring based on their evidence-selectivity (see Methods) and ordered based on the position of peak activity. **(C)** As in (B) but for retrosplenial cortex (RSC, n = 4190 neurons, data from Koay et al. (2021)^66^). **(D)** As in (B) but for hippocampus (HPC, n = 791 neurons, data from Nieh et al. (2021)^49^). **(E)** As in (B) but for dorsomedial striatum (DMS, n = 322 neurons, see Methods). Sequences separated by even and odd trials are shown in Extended Data Fig. 1.

Across brain regions, neurons fired sequentially rather than persistently (Fig. 1B-E; Extended Data Fig. 1). Such sequences were present throughout the task, beginning in the cue region and continuing throughout the delay region. Moreover, many of the neurons in these sequences were choice-selective, such that neurons preferentially fired depending upon whether the animal will ultimately choose to turn left or right (Fig. 1B-E).

Previous models of evidence accumulation do not produce the observed choice-selective sequences. This includes traditional circuit models that neurally instantiate the drift diffusion model (Fig. 2A-B), as well as bump-based integrators (Fig. 2C-D) commonly used to accumulate head velocity signals into a representation of head direction. Further, canonical sequence models such as synfire chains^60–62^ and more recent models of replay sequences^63^ do not accumulate evidence, while other sequence models accumulate external signals but are not choice-selective^64,65^. This motivates our construction of new, sequence-based evidence-accumulation models.

**Figure 2.**
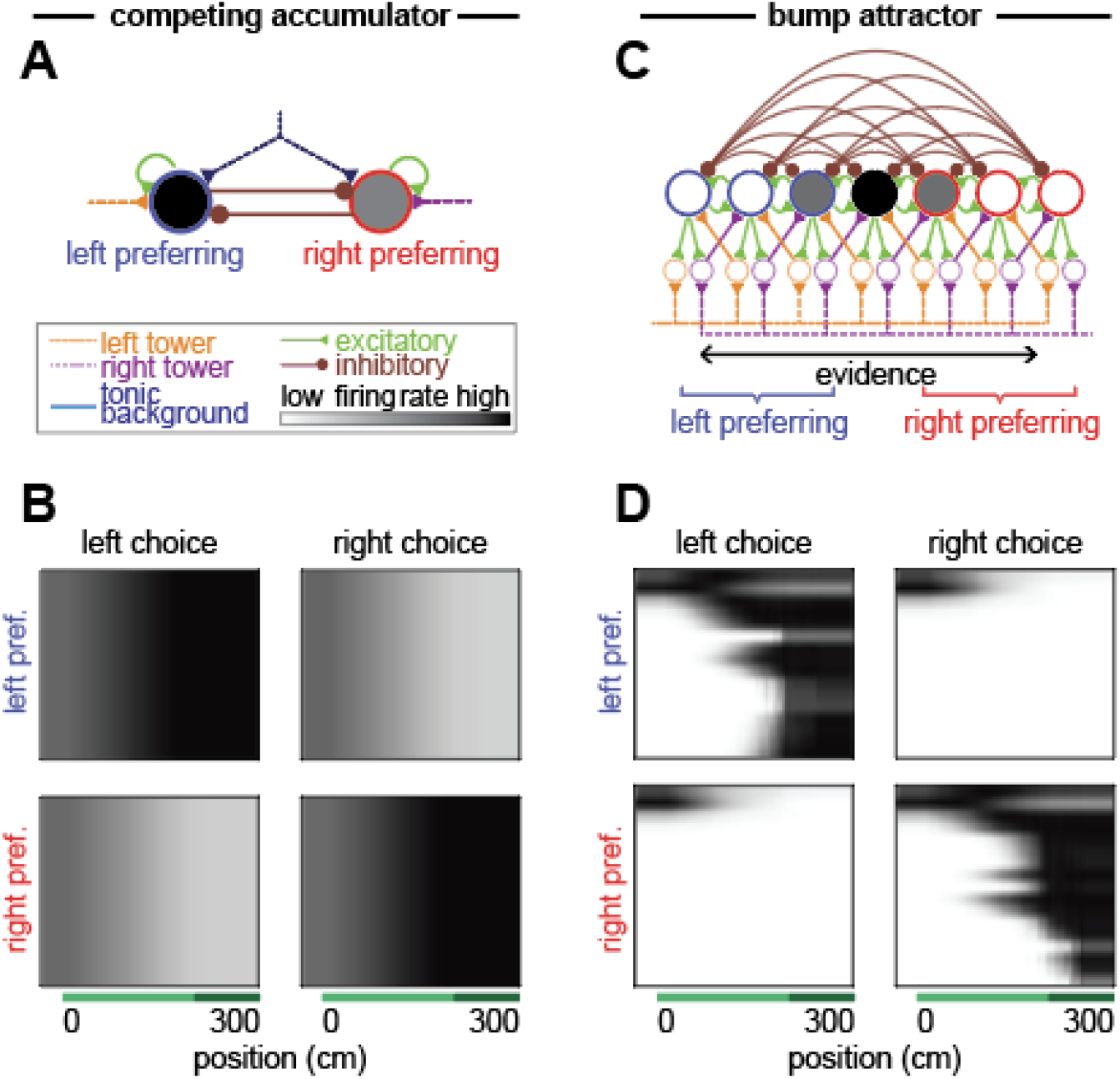
Choice-selective sequences are not predicted by traditional integrator models. **(A)** Schematic of a traditional mutually inhibitory integrator model for decision making. **(B)** Average peak-normalized, simulated activity of neurons in the model during task performance, divided into the left and right population and sorted by the peak of mean firing activity. **(C-D)** As in (A-B), but for a traditional bump attractor model. For model equations, see Methods.

### Competing Chains Models

We first present a class of models consisting of two chains of neurons. In these models, the current position of the animal is represented by the location of elevated neural activity along the chains and the accumulated evidence is represented by the relative amplitude of firing of the two neurons corresponding to this location. Here, we consider a “mutually inhibiting” chains model with mutual inhibition between neurons at the same position in the chains (Fig. 3A), but other competing chains models are possible (see below and Supplementary Text). The temporal dynamics of the firing rate *r*_*i,L*_ of the *i*^th^ neuron in the left (L) side chain are given by the equation

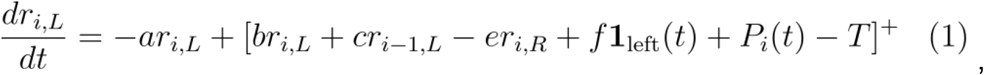

and an analogous equation describes the firing rates *r*_*i,R*_ of neurons in the right (R) side chain. Here, *a* is the exponential decay rate of activity in the absence of input, *b* is the weight of self-excitation (Fig. 3A, green self-loops), *c* is the strength of feedforward synaptic connections from the previous neuron (green feedforward connections), and *e* is the strength of the inhibitory connection from the neuron at the same position in the opposite chain (brown connections). The external inputs to be integrated by the network consist of brief pulses of the form ***1***_*left*_*(t)*, which is defined to be 1 if there is a left cue within 0.5 cm of the current position and 0 otherwise, and project to the chain with synaptic strength *f. P*_*i*_(*t*) is the position-gating signal, and *T* is the firing threshold. The position-gating signal *P*_*i*_(*t*) controls the progression of activity along the chain by raising the neuron’s input above the firing threshold *T*, and is zero for all neurons except the pair of neurons (one from each chain) corresponding to the position of the animal in the environment. The firing rates of neurons not at the active position decay exponentially with time constant *1/a*. Firing rates and inputs during a subset of an example trial are shown in Fig. 3B. See Methods for complete details and parameterization of the model.

**Figure 3.**
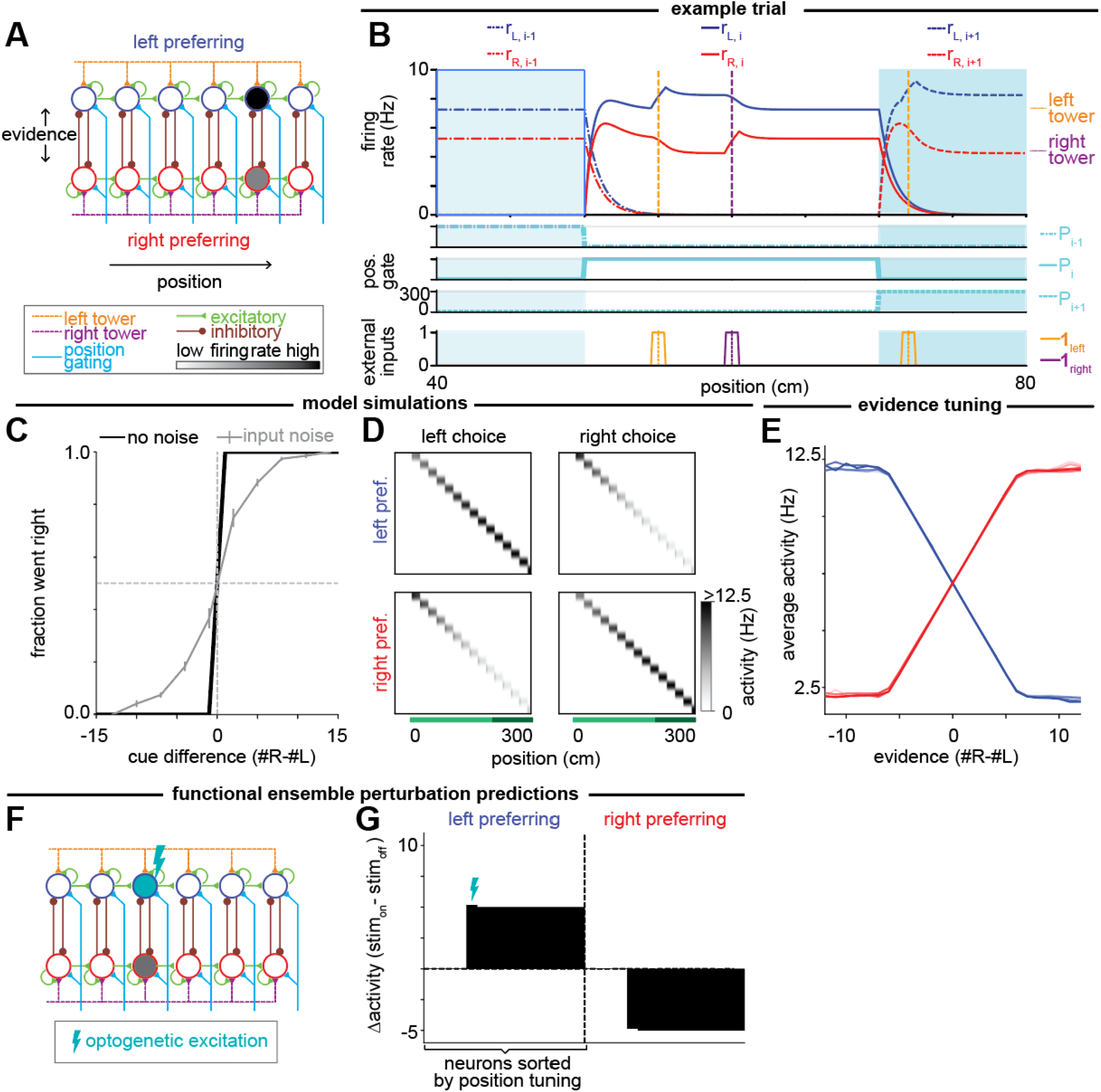
Mutually inhibiting competing chains model of evidence accumulation through sequences. **(A)** Schematic of neural circuit model for the mutually inhibiting competing chains model, showing excitatory (green) and inhibitory (brown) connections between neurons (circles) as well as external inputs to the circuit from the left (orange) and right (purple) towers and a position gating signal (cyan). **(B)** Top: Firing rates of neurons in the left (blue) and right (red) chains during an example trial, shown for positions 40 to 80 cm. Vertical dashed lines indicate the location at which left (orange) and right (purple) towers appeared. Background shading highlights the regions where position gates *P*_*i - 1*_, *P*_*i*_, and *P*_*i+1*_ are on. Middle: Position gating signal at each position. Bottom: Input current to the left (orange) and right (purple) chains resulting from the external inputs. **(C)** Psychometric data for the model simulated on 1000 trials describing how often the amplitude of the final neuron in the left chain was greater than that of the final neuron in the right chain for cases where the model was simulated with (gray) and without (black) noise in the input. Error bars show standard error of the mean (s.e.m.). **(D)** Each row shows the non-normalized amplitude of a model neuron at each position in the maze, averaged across simulated trials without input noise when the greater final amplitude was in the left (left choice, left column) or right (right choice, right column) chain. Neurons were divided based on their choice-selectivity (equivalent to the left and right chains in this model; see Methods) and ordered based on the position of peak activity (equivalent to position in the chain). **(E)** Tuning curves of individual neurons to evidence, defined by the average activity for different evidence levels when the neurons are activated by the position-gating signal, for left-preferring (blue) and right-preferring (red) neurons. **(F)** Schematic of a single-neuron perturbation experiment where optogenetic excitation is applied to a single neuron in the left chain. **(G)** Simulated changes of the firing rates of all neurons in the absence of cues when a single neuron (denoted with the lightning bolt as in (F)) is optogenetically stimulated.

At a given position, this model has the same structure as the traditional mutual inhibition model^26^, and this motif is repeated across different positions to form two chains. This motif accumulates and maintains evidence through positive feedback within and between the two neural populations receiving the active position gating signal. Competition is mediated by mutually inhibitory positive feedback between the chains, such that increases of activity in one chain results in corresponding decreases of activity in the other chain. We note that the accumulation process saturates at an upper bound when neurons in the non-dominant chain reach zero firing. This is because, when one chain is at zero firing rate, the disinhibitory component of positive feedback between the chains is lost, so that further increases in the magnitude of evidence cannot be maintained. This saturation is not a core feature of the model, however, and the non-saturating range of evidence levels can be modified by the amount of external input to each chain or the synaptic strength of the visual input connections (Extended Data Fig. 2J). As in the case of traditional, non-sequential accumulator models, perfect accumulation up to this upper bound requires fine-tuning of the parameters to prevent leak or uncontrollable growth in the integrator^35^ (see Supplementary Text). Previous studies of such networks have also suggested that such accumulator networks may be made more robust to these fine-tuning conditions through incorporation of bistable components^67,68^ or corrective feedback mechanisms^69,70^.

Only the neurons receiving the position gating signal actively represent and accumulate evidence. When the animal moves to a new position, the position gating signal activates the next pair of neurons in the chain, and evidence is transferred to the newly activated pair of neurons by the feedforward synaptic connections^64^ (see Supplementary Text for conditions on the feedforward connectivity for faithful transfer of evidence between positions). Repeating this process as the animal navigates down the track results in a sequence of neurons firing, with each successive pair of activated neurons representing the accumulated evidence in the difference of their firing rates.

We simulated the model on the accumulating towers task (see Methods), demonstrating that this model performs the task perfectly in the absence of noise for appropriately tuned parameters (Fig. 3C, see Supplementary Text). Previous analysis of the accumulating towers task and other evidence accumulation tasks suggest that noise in an animal’s performance is dominated by sensory noise, so that only a fraction of input cues are integrated^13,59^. When we correspondingly introduce input noise into our model (see Methods), the performance decreases in a manner similar to that observed in animals, with a more gradual increase in performance with increasing magnitude of the difference between cues (Fig. 3C). The neurons in the model exhibit choice-selective (or, more precisely, “accumulated-evidence-selective”) sequential activity, with neurons in each chain showing greater activity for the side of the evidence that it receives (Fig. 3D). Thus, accumulated evidence is encoded monotonically (up to saturation) in the relative firing rates of the two chains in this class of models (Fig. 3E).

Variant architectures of competing chains models are possible, including an “uncoupled” competing chains model where competition between opposing inputs occurs without inhibition between the chains (Extended Data Fig. 2A, see Supplementary Text). At a given position, this model has a “push-pull” feedforward arrangement of evidence inputs, with cues favoring one side exciting one chain and inhibiting the opposing chain, as in traditional decision-making models based on uncoupled competing accumulators^27^. This motif is then repeated to form two chains. As with the model based on mutually inhibiting chains of neurons, this uncoupled competing chains model is able to perform the task accurately (Extended Data Fig. 2B) and produce choice-selective sequences (Extended Data Fig. 2C) in which individual neurons have monotonic tuning curves to evidence (Extended Data Fig. 2D). As in the mutually inhibitory chains model, this model can either saturate or not depending on the strength of the synaptic input connections and assumptions about the dynamic range of the neurons (Extended Data Fig. 2K).

Although these variant competing chains models cannot be reliably distinguished by their tuning since both models can be modified to have either linear or saturated tuning curves, the models instantiate different hypotheses that could be distinguished by functionally-targeted optogenetic perturbations^71^ (Fig. 3F, Extended Data Fig. 2E). We simulated the delivery of an excitatory input to a neuron (or similarly tuned neuronal population) while the animal is at the neuron’s active position (Fig. 3F, Extended Data Fig. 2E) for a trial with no external cues so that the accumulated evidence prior to perturbation is zero. Due to the gating of neural activity by the forward-traveling position signal, both models predict that only neurons tuned to later positions will be affected by the optogenetic perturbation. For the mutually inhibiting chains model, subsequent neurons in the same chain will have increased activity due to this feedforward excitation while subsequent neurons in the opposing chain will have decreased activity due to the mutual inhibition between the chains (Fig. 3G). By contrast, for the models based on uncoupled chains in which the anti-correlated activity of the two chains is solely due to opposing external cue inputs (Extended Data Fig. 2), an excitatory perturbation in one chain increases firing only within the same chain but has no impact on the other chain (Extended Data Fig. 2F).

Another difference between the mutually inhibiting and uncoupled competing chains models is that the model based on mutual inhibition is more robust to the tuning of the position-gating. Specifically, the mutual inhibition model requires only that the position-gating signal *P*_*i*_ be greater than or equal to the threshold *T* at the active position (Extended Data Fig. 2G,H), whereas the uncoupled chains model requires that *P*_*i*_ be exactly equal to *T* to avoid continual growth of neural activity from temporally integrating the position-gating signal^72^ (see Supplementary Text).

Taken together, these competing chains models represent a class of models in which position-gating gives rise to choice-selective sequences that encode graded evidence signals monotonically in the neuronal firing rates at each position.

### Position-Gated Bump Attractors

We next consider a class of models in which evidence is encoded in the location of a stereotypical, unimodal pattern (“bump”) of neural activity in the population. Such bump attractor models have been used to describe the neural circuitry that computes head direction in the rodent^73–78^ and the fly^79–85^, where the location of the bump corresponds to the animal’s heading. Similarly, two-dimensional bump attractor models have been proposed to describe path integration in the hippocampal place cell system^86–88^. In these models, motion is temporally integrated to determine head direction or to perform path integration, and analogously our models temporally integrate evidence from the visual cues. A traditional bump model that only accumulates evidence does not produce sequences like those observed in the neural data because the bump would recur at the same location whenever the accumulated evidence is at the same level and would remain at a fixed location throughout the delay period when the evidence level does not change (Fig. 2D). Thus, we modify these traditional bump attractor models to have separate dimensions for encoding position and evidence (Fig. 4 and Extended Data Fig. 3).

**Figure 4.**
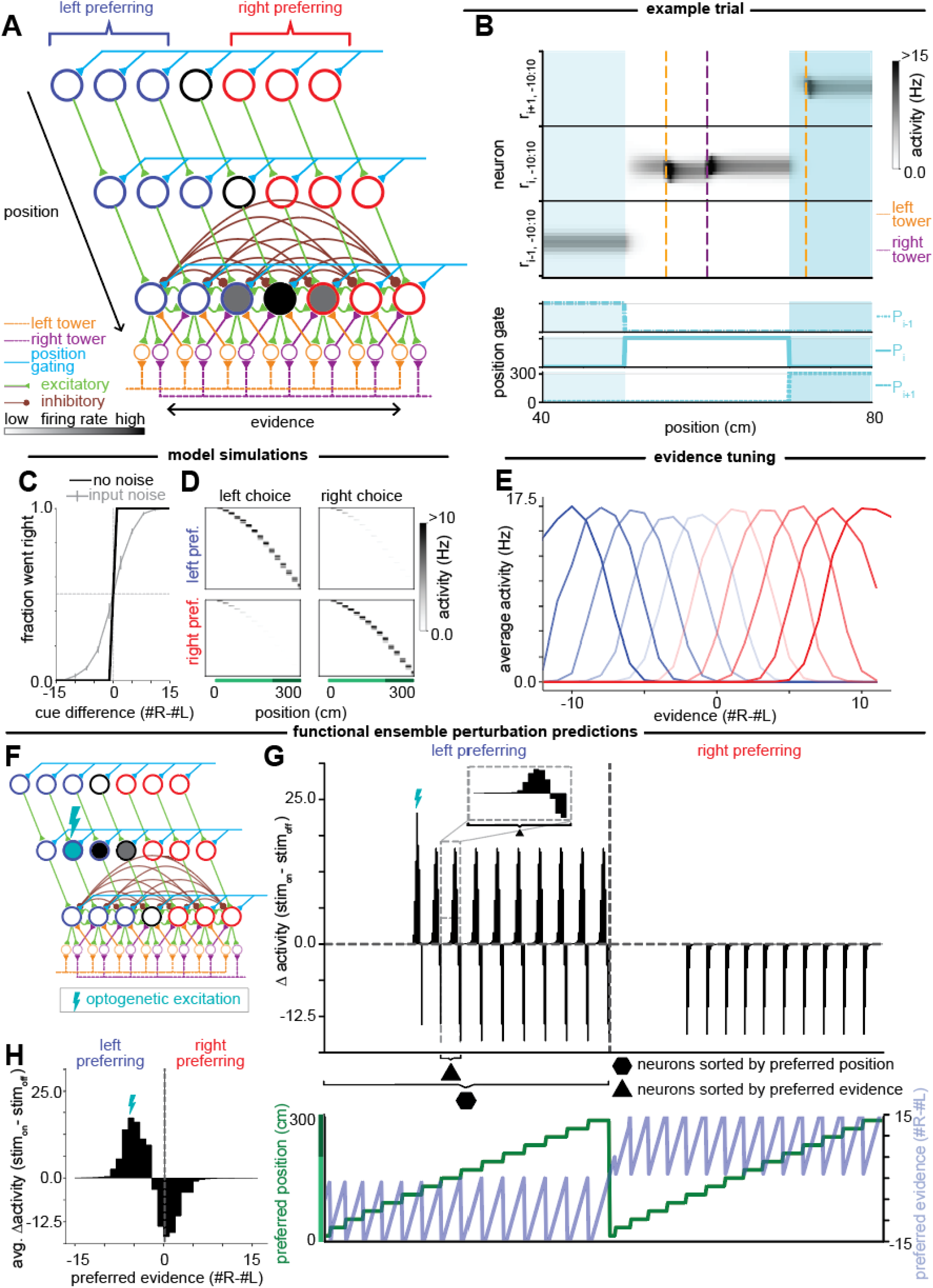
Position-gated bump attractor model of evidence accumulation through sequences. **(A)** Schematic of neural circuit model for the position-gated bump attractor model. Each row represents neurons (circles) that respond at a given position. Within a row, each neuron represents a different evidence level, ranging from left-most to right-most. Blue: left-preferring neurons; red: right-preferring neurons. Activation of neurons encoding a given position is controlled by a position gating signal (cyan). Transfer of information between positions is controlled by excitatory feedforward connections between successive positions (green). Bottom row of neurons illustrates the cue-related inputs received by, and connectivity between, neurons within any given row: local excitatory (green) and broader inhibitory (brown) connections between neurons as well as external inputs to the circuit from the left (orange lines) and right (purple lines) towers via the corresponding shifter neurons (orange and purple circles). **(B)** Top: Each row shows the firing rate of a subset of neurons, shown for positions 40 to 80 cm during an example trial of the maze. Neurons are first sorted by position tuning (black horizontal lines denote divisions between positions) and then by evidence tuning within a position. Dashed vertical lines indicate the position at which the left (orange) and right (purple) cues appeared. Background shading highlights the regions where position gates *P*_*i - 1*_, *P*_*i*_, and *P*_*i+1*_ are on. Bottom: Position gating signal at each position. **(C)** Psychometric data for 1000 simulated trials, indicating how often the index of the maximally firing active neuron in the final layer of neurons was to the left of the center neuron for cases with (gray) and without (black) noise in the input. Error bars show s.e.m. **(D)** Each row shows the non-normalized firing rate of a model neuron at each position in the maze, averaged across simulated trials without input noise when the peak of the bump was in the left-preferring (left choice, left column) or right-preferring (right choice, right column) population. Neurons were divided based on their choice-selectivity (see Methods) and ordered based on the position of peak activity. **(E)** Tuning curves of a subset of individual neurons to evidence, calculated when the neurons are activated by the position-gating signal, for left-preferring (blue) and right-preferring (red) neurons. **(F)** Schematic of a single-neuron perturbation experiment in which optogenetic excitation is applied to a single neuron in the left-preferring population. **(G)** Top: Simulated changes of the firing rates of all neurons in the absence of cues when a single neuron (denoted with a lightning bolt as in (F)) is optogenetically stimulated. Neurons are first sorted by left- or right-preferring, then by position tuning, and then by evidence tuning within a position. Bottom: Corresponding preferred position (green) and preferred evidence (light blue) tuning of each neuron shown in the top panel. **(H)** Average change in firing rate for neurons tuned to different evidence levels.

The model consists of layers of recurrently connected neurons, with each layer corresponding to a given position and neurons within the layers representing the accumulated evidence at this position. The activity *r*_*i,j*_ of a neuron at position *i* and evidence level *j* evolves according to

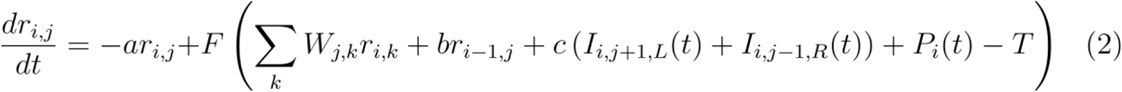

where *a* is the exponential decay rate of activity in the absence of input, *P*_*i*_(*t*) is the position-gating signal that selects the active layer of neurons, and *T* sets the “soft” threshold of the (1 + tanh(·)) neuronal output nonlinearity *F*. Within a layer, *W*_*j,k*_ is the strength of the connection from the neuron representing evidence level *k* onto the neuron encoding evidence level *j. W*_*j,k*_ is excitatory for connections of a neuron onto itself and its immediate neighbors (Fig. 4A, green recurrent connections) and inhibitory onto all other neurons in that layer (Fig. 4A, brown recurrent connections). The coefficient *b* is the synaptic strength of feedforward excitatory connections between layers of neurons encoding neighboring positions, and we assume for simplicity that these connections are only between neurons encoding the same evidence level (Fig. 4A, green feedforward connections). The terms *I*_*i,j+1,L*_ and *I*_*i,j-1,R*_ are the firing rates of the “shifter neurons” (analogous to the “rotation cells” in Skaggs et al. (1995)^73^) (Fig. 4A, orange and purple circles) that shift the bump left or right in response to external cue inputs (Fig. 4A, left and right tower inputs; Fig. 4B, shift in location of the bump following an input). The left shift (orange) neuron is activated when jointly receiving input from evidence accumulating neuron *j+1* and a left-side external cue. In this manner, if the peak of the bump of neuronal activity was previously at *j+1*, this peak activity then (through a synaptic connection of strength *c* from the shifter neuron) drives the neuron at evidence level *j*. Mathematically, the firing rates of the left, and corresponding right, shifter neurons are given by

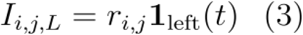

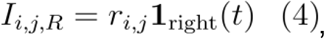

where ***1***_*left/right*_(*t*) equals 1 if there is a left or right cue, respectively, within 0.5 cm of the current position. For the complete model specification, see Methods.

Within the active position, accumulated evidence is represented and maintained in the location of a bump of activity along the evidence axis. As in traditional bump attractor models^89–91^, the shape of the bump is maintained in the absence of input by local symmetric excitation and long range inhibition. Asymmetric input connections cause the bump to shift left or right in response to a left or right cue, respectively^74,78^, as described above (see Supplementary Text). This shift of the localized bump of activity occurs despite the external cue signal being global (Fig. 4A, tower inputs). Mechanistically, the ability to shift a localized bump of activity with a global external signal is achieved via the multiplicative gating of the external input by the activity of the evidence-encoding neurons (Eqs. (3) and (4)) so that the global external input signal only activates the shifter neurons close to the location of the current bump. Alternatively, this gating could be accomplished with additive inputs plus thresholds^85,89^.

As the animal progresses forward in the maze, the active position changes so that information must be transferred to the next layer of neurons, requiring both a change in the position gate and feedforward connections between layers (Fig. 4B). Here, position-gating is controlled by the position signal *P*_*i*_(*t*) and the “soft” threshold *T*, so that the output of *F* is only significantly greater than zero when the position signal is active (i.e., *P*_*i*_(*t*) ≥ *T*). Away from the active position (i.e., where *P*_*i*_(*t*) = 0), activity decays exponentially with time constant *1/a* (Fig. 4B). Feedforward excitation between the layers then transfers information to neurons at the next position. We have modeled these feedforward connections as only projecting between neurons representing the same evidence level, but this projection could be broader as long as the feedforward input is symmetric and centered around the current evidence level (see Supplementary Text). As in the chains-based models, we have presented the position signal as an external input. However, position could simultaneously be generated within the network through an integration of velocity inputs, by including a similar arrangement of excitatory and inhibitory connectivity along the position axis. In this case, the network forms a planar bump attractor that jointly encodes both evidence and position (Extended Data Fig. 3, see Supplementary Text).

We simulated the position-gated bump attractor model for the same set of trials as the competing chains models. As in the competing chains models, for appropriately tuned parameters, the position-gated bump attractor also performs the task perfectly in the absence of noise, with performance worsening when input noise is introduced (Fig. 4C). The simulated neural activity also produces choice-selective sequences (Fig. 4D). As seen by the downward curvature of the sequences in Figure 4D, the sequences are marked by the feature that more neurons respond at late positions than respond at early positions in the sequence. This is because the random walk of the bump location as evidence is accumulated leads to a greater range of evidence levels being observed at later positions in the trial; thus, on average across trials, there are more different neurons active at later positions, with each neuron corresponding to a different possible evidence level. However, we note that this is not a fundamental prediction of the model because, experimentally, the exact shape of the choice-selective sequence depends on the number of neurons residing within each uniformly spaced evidence-position location in the model (Fig. 4A). We also note that our assumption of perfect symmetry simplifies the analysis, but is not required to produce bump attractor models^92,93^ and that the robustness of bump attractor models to diffusion and drift of the bump in the presence of noise has been extensively studied^92,94–96^ and may be reduced through error-correcting codes^97^, extending representations to multiple bumps^98^, or relaxing the continuous attractor assumption to have only a discrete set of memory states^99^.

The most straightforward difference between the position-gated bump attractor and the competing chains models is in the shape of their evidence tuning curves. Due to the structure of the network connectivity, the neurons in the position-gated bump attractor are non-monotonically tuned to specific evidence levels at their active position (Fig. 4E), in contrast to the monotonic tuning curves of the competing chains models (Fig. 3E).

To further differentiate the position-gated bump attractor model from the competing chains models, we consider predictions for the effects of optogenetic excitation of a single neuron during a trial with no cues presented (Fig. 4F). Here, we consider the regime where the stimulus is sufficiently large to cause the bump to move to the stimulated location. Due to the similar feedforward connectivity patterns between positions in both model classes, stimulation of a neuron active at one position only affects the activity of neurons tuned to later positions, as in the competing chains models. However, unlike the competing chains models, we see that rather than impacting every neuron at a later position equally, the effects are sparse and differ in magnitude across neurons (Fig. 4G). These differences are because the stimulation has caused the bump to move to a new evidence level, which is then sustained at subsequent positions. Thus, neurons with a similar evidence tuning to the neuron that was stimulated increase in activity, while those tuned to where the bump of activity would be located in the absence of stimulation (at 0 evidence) decrease their activity since the bump has moved to the new location (Fig. 4H). We note that for smaller excitatory stimuli that do not shift the bump center to the site of stimulation, and dependent upon the exact details of the excitatory and inhibitory connections, smaller shifts in the bump either toward or away from the stimulated location are possible^96,100–102^, but the perturbations still maintain the properties that changes are sparse and that changes within the same choice-preferring population as the stimulated neuron may be bidirectional.

### Neural Activity in Different Brain Regions Correlates with Different Classes of Models

Above, we presented two novel circuit models for accumulating evidence through sequences (competing chains models: Fig. 3, Extended Data Fig. 2; position-gated bump attractor model: Fig. 4). While both models produce neurons that are sequentially tuned to position, the models predict different evidence coding schemes. The competing chains models make two core predictions: first, evidence should be monotonically encoded in neuronal activity and, second the monotonic encoding within the population should be graded across a range of evidence levels. The bump attractor model likewise makes two core predictions: first, evidence should be encoded with relatively narrow, unimodal tuning and, second, the peaks of these tuning curves should be broadly distributed across the entire evidence range. We looked for signatures of these different evidence encoding schemes in recordings from ACC, RSC, DMS and HPC during the accumulating towers task (Fig. 1A).

Towards this end, we first identified cells that were evidence-tuned, showing any consistent response pattern at different accumulated evidence values. For these evidence-tuned cells, we then identified whether neurons were monotonic-like or unimodal-like in their evidence tuning and also, for unimodal tuning, we characterized whether the peaks of the tuning curves were broadly distributed across the range of evidence levels. Within the class of monotonically evidence-tuned neurons, we next assessed whether the evidence tuning curves had broadly graded tuning to evidence, versus more binary tuning that would be consistent with a “choice-like” representation, and also how this distribution varied with position in the maze. We further examined whether graded evidence could be decoded from the population as a whole and whether the encoding of evidence across the population was consistent with the prolonged, temporal response kernels required for graded accumulation of evidence.

#### HPC has Narrow Unimodal Tuning that Spans Position and Evidence Space While ACC and RSC Primarily have Monotonic Evidence Tuning

Identifying neurons that were tuned to evidence requires jointly, rather than independently, fitting position and evidence to account for the correlated, nonuniform sampling of these two variables (Extended Data Fig. 4A-B). For example, neurons tuned to early positions in the maze are only active when accumulated evidence is small, given that little evidence has been presented at early positions. Thus, if such neurons were directly fit to evidence tuning alone, rather than jointly fit to evidence and position, they would appear to be tuned to small absolute evidence levels even if they had no tuning to evidence at all. Likewise, because the evidence levels sampled at later parts of the maze were most concentrated around moderate-to-large absolute values (Extended Data Fig. 4A), neurons untuned to evidence but tuned to later positions in the maze would artifactually appear to have a bimodal tuning to evidence.

To capture many different tuning curve shapes across the population, we first modeled the joint tuning for position and evidence as a product of a Gaussian function of position and a Gaussian function of evidence (Extended Data Fig. 4B, Fig. 5C,D,G,H,K,L, Extended Data Fig. 5C,D; see Methods). The Gaussian in position intuitively captures the non-monotonic, sequential position tuning. The Gaussian in evidence, although non-monotonic in its mathematical form, allowed us to identify not only non-monotonic but also monotonic evidence tuning, because in the latter case the best fit Gaussian has a peak near the extreme of the evidence range (i.e., normalized peaks near +/- 1; Extended Data Fig. 4D). Broad monotonic ramps like those of the competing chains models also have a large standard deviation relative to the observed evidence range.

**Figure 5.**
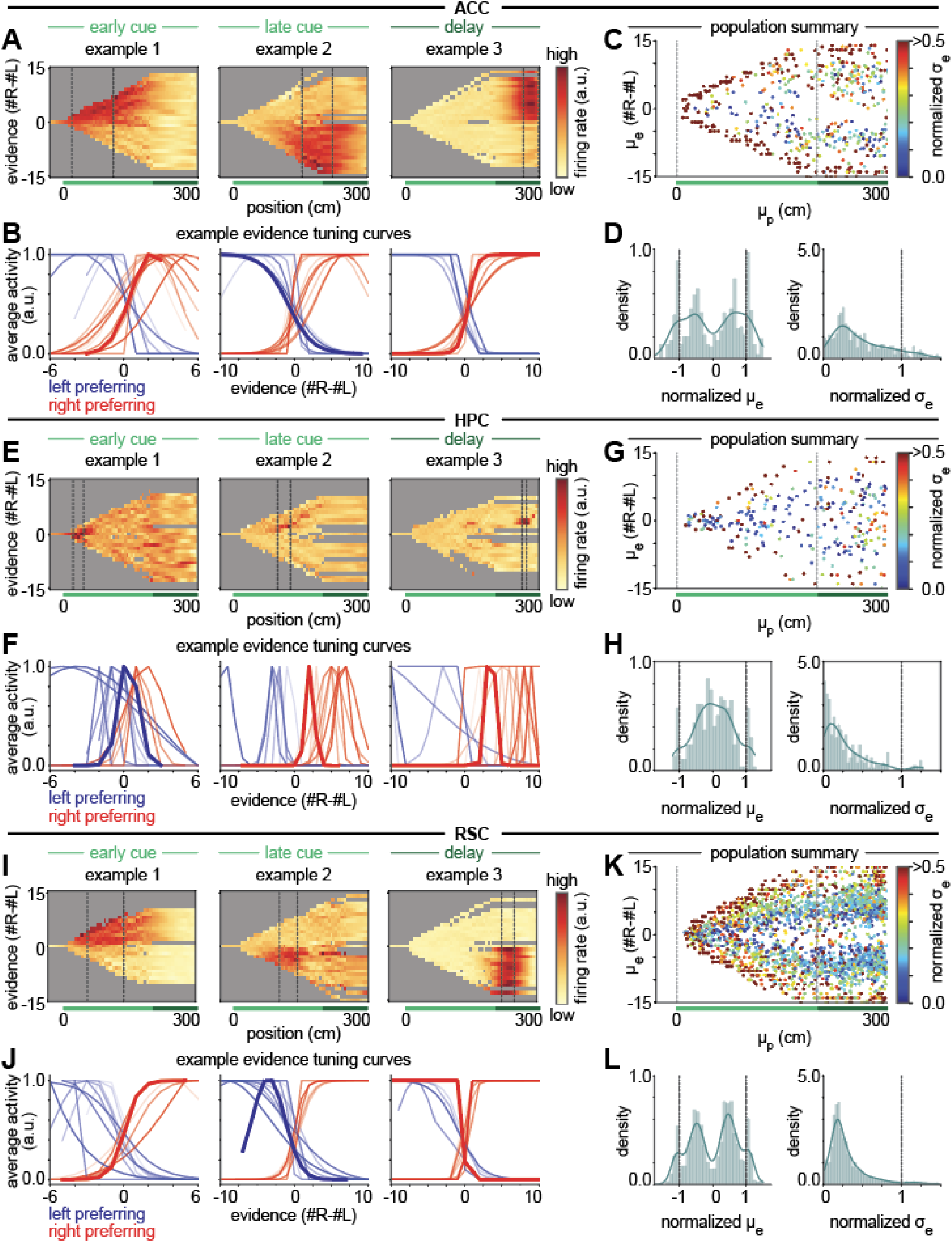
HPC has narrow unimodal tuning that spans position and evidence space while ACC and RSC primarily have monotonic evidence tuning. **(A)** Heatmaps showing the average firing in position-by-evidence bins for example individual neurons in ACC with peak position tuning in the early cue (left), late cue (middle), or delay (right) region of the maze. Gray bins denote regions for which there were fewer than 2 samples during the session. **(B)** Example ACC evidence tuning curves fit to the region of the neuron’s peak activity (defined as observations within 0.5 position standard deviations of the neuron’s fit peak position tuning; see Methods) for a collection of neurons with mean position tuning in the early cue (left), late cue (middle), or delay (right) region of the maze. Red coloring indicates neurons classified as right-preferring, and blue indicates left-preferring. Bold lines correspond to the examples in (A), for which the neuron’s region of peak activity is the region between the dashed vertical lines. For comparison, the raw evidence tuning curves are provided in Extended Data Fig. 6. **(C)** Scatter plot showing the location of the fit mean position (*μ*_*p*_) and fit mean evidence (*μ*_*e*_) with color indicating the normalized fit evidence standard deviation (*σ*_*e*_) (see Methods) of the 80% of neurons recorded in ACC with the best fit between the neural data and the model predictions. Narrowly tuned neurons are only plotted if they pass non-outlier criteria (see Methods). Dashed lines indicate the boundaries of the cue period. **(D)** Density plots of the normalized fit mean evidence (left) and the normalized fit evidence standard deviation (right) for the neurons shown in (C). Solid blue lines show the kernel density estimate. **(E-H)** Same as (A-D) but for neurons recorded in HPC. **(I-L)** Same as (A-D) but for neurons recorded in RSC. Distributions for all evidence-modulated neurons (regardless of goodness-of-fit and outlier tests) are shown in Extended Data Fig. 12.

Jointly fitting position and evidence tuning across the 14,247 neurons revealed that 56% of all neurons in ACC, 48% in RSC, 25% in HPC, and 41% in DMS had significant evidence tuning (showing any pattern of response to evidence with statistical comparison to pseudosessions; see Methods). In analyzing neurons with significant evidence tuning in each region, we saw distinguishing patterns emerge in the nature of the evidence tuning (Fig. 5, Extended Data Fig. 5).

In ACC, many individual neurons exhibited primarily monotonic and relatively broad evidence tuning (Fig. 5A-B, Extended Data Figs. 6A,7). To accurately visualize evidence tuning, in addition to plotting raw position and evidence fields (Fig. 5A), we also fit both a Gaussian and a logistic function to the peak position tuning (Extended Data Fig. 4C) for a number of example neurons, and plotted the best fit function (Fig. 5B,F,J). Similar to these example neurons, among the population of significantly evidence-tuned neurons, peak evidence tuning was towards the extreme of the observed evidence range, consistent with monotonic tuning (Fig. 5C-D). The presence of monotonic tuning is most consistent with predictions of the competing chains models (Fig. 3E, Extended Data Fig. 2D,J,K, Extended Data Fig. 4E,F), although the logistic fits revealed many neurons had steeper sigmoidal tuning than produced by the competing chains models (Fig. 3E, Extended Data Fig. 2D), especially at later positions in the maze (Fig. 5B, Extended Data Fig. 6A). The steepness of such sigmoidal tuning, and its variation across different positions of the maze, is treated further in the next section.

In contrast, HPC neurons showed primarily non-monotonic and narrow evidence tuning (Fig. 5E,F, Extended Data Figs. 6B,8), which tiled the range of observed evidence (Fig. 5F-H). These narrow tuning curves occurred throughout the cue and delay region (dark blue neurons in Fig. 5G). The presence of non-monotonic, narrow tuning curves that tile evidence space is consistent with predictions of the bump model (Fig. 4E, Extended Data Fig. 4G,H). In addition to narrow tuning curves, the bump model also predicts that the population should track the accumulated evidence on individual trials. Such narrow tuning curves make decoding challenging because, to perfectly decode evidence, recordings would need to include cells tuned to each combination of position, evidence, and other variables previously identified in this task^49^, and the HPC recordings are sparse. However, previous analysis of these data suggests that evidence is represented in the population on a nonlinear neural manifold in HPC^49^, and we similarly find that evidence can be decoded above chance using another nonlinear embedding technique^103^ (Extended Data Fig. 11).

In RSC, the majority of neurons showed monotonic patterns (Fig. 5I-J, Extended Data Fig. 9) like those observed in ACC, while a smaller population showed more narrow tuning curves like, but typically somewhat wider than, those observed in HPC (example 2 in Fig. 5I, examples 3, 9, 16, 22, 28, and 34 in Extended Data Fig. 9). These observations are seen in the population summary of evidence and position tuning (Fig. 5K-L), which shows that many neurons have wide evidence tuning with peaks at the extreme (deep red) while many others have narrow intermediate evidence tuning (dark blue). RSC also has a particularly large population of neurons that, when fit to a Gaussian, have intermediate widths and evidence peaks not fully at the extremes (Fig. 5K, green); such neurons typically exhibit a steep sigmoidal tuning curve such as that shown in example 3.

DMS was dominated by monotonic evidence tuning, but with some narrowly tuned neurons (Extended Data Fig. 5, Extended Data Fig. 10). However, DMS contained relatively few evidence-modulated neurons with peak position tuning in the cue region. Instead, most evidence-modulated neurons had their peak position tuning in the delay region and had “choice-like” steep sigmoidal evidence tuning (Extended Data Fig. 5C,E).

#### Shift from Evidence Tuning in the Cue Period to Choice Tuning in the Delay Period in ACC and RSC

While the monotonicity of the evidence tuning of individual cells observed in ACC and RSC is consistent with the competing chains models, a candidate integrator must represent graded evidence beyond monotonic, step-like evidence tuning driven by choice. Since the units of our models should not be thought of as individual cells, but rather populations of neurons with similar position tuning, the graded representation predicted within our model units could emerge from either individual cells having graded representations or individual cells stepping on at different evidence thresholds such that population averages are graded^104^. Therefore, we examined whether the regions dominated by monotonic tuning for evidence contained representations of evidence in the population beyond binary choice tuning, using both single-cell and population-level analyses.

We first asked whether there were signatures of evidence accumulation within the population by deriving cue response kernels from a linear cue encoding model fit to the difference in right- and left-preferring population activity (Fig. 6A-C, see Methods). As required for integration, we found cue-locked, sustained, and relatively constant steplike responses to the cues, with left and right cues having a similar pattern but with opposite signs (Fig. 6D,E). When accounting for the amount of evidence accumulated so far, the magnitude of the cue responses decreased, consistent with a saturation at more extreme evidence levels and a more choice-like representation as the maze progresses (Extended Data Fig. 13A,B). Interestingly, the cue response kernels decreased less as a function of position in the maze than as a function of accumulated evidence (Extended Data Figure 13C,D; note nonzero kernels even for later positions), suggesting that the transition observed as a function of previously accumulated evidence may be due to animals committing to a decision with larger levels of evidence^105,106^, rather than solely being due to being at a later position in the maze. Most importantly, this analysis suggests that accumulated evidence is represented beyond choice at the population level (at least for intermediate values of evidence).

**Figure 6.**
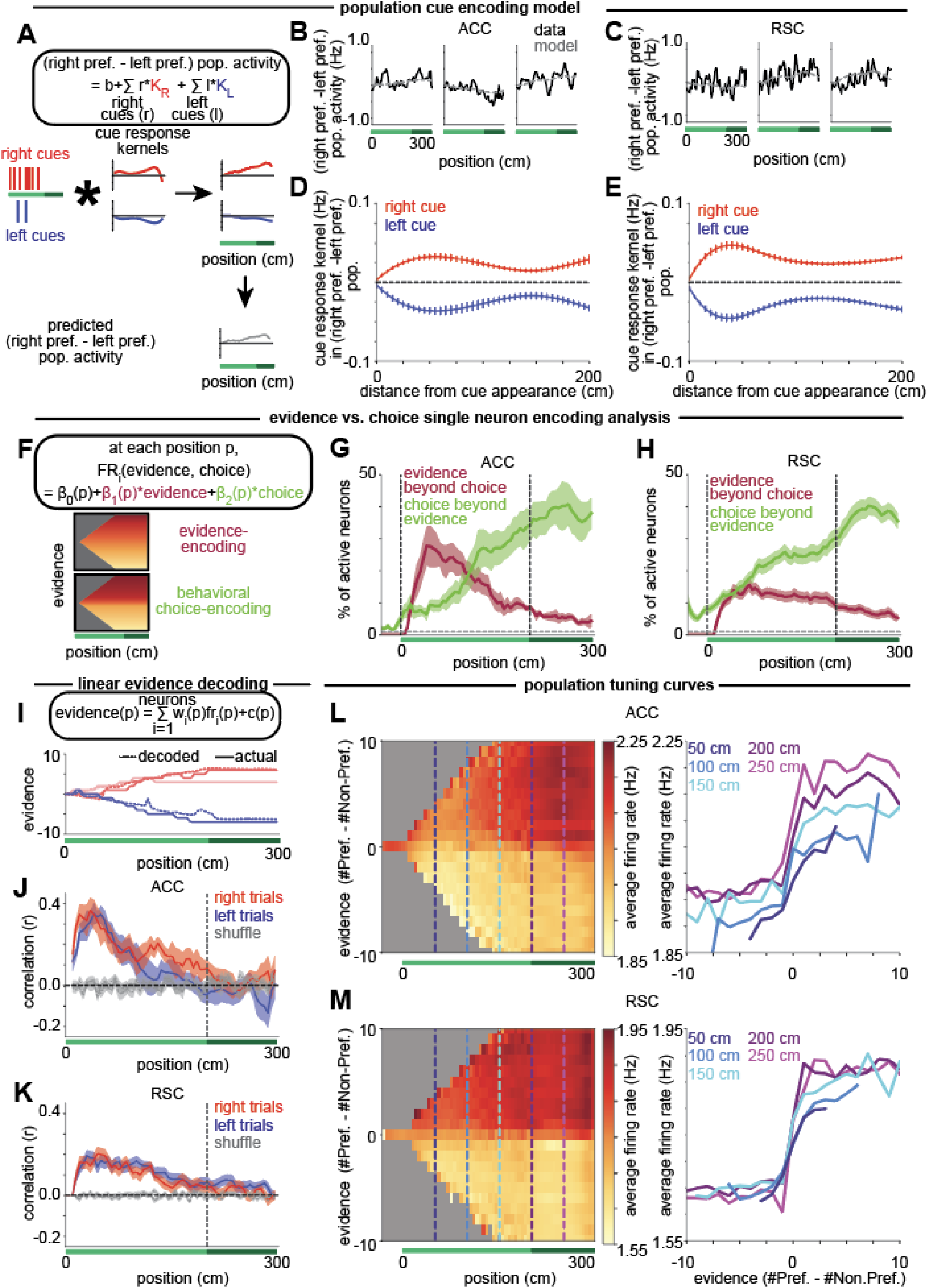
In ACC and RSC, Evidence Tuning in the Cue Period Shifts to Choice Tuning in the Delay Period. **(A)** Schematic of the population cue encoding model, where each left and right cue is convolved with a left or right cue response kernel, *K*_*L*_ or *K*_*R*_ respectively, to predict the population activity. **(B)** Example single-trial population activity (black) compared to the model predicted activity (dashed gray) in ACC. (Average r^2^ = 0.26 across sessions.) **(C)** Same as B but for RSC. (Average r^2^ = 0.20 across sessions.) **(D)** Cue kernels fit from the cue encoding model to right cues (red) and left cues (blue). Error bars indicate s.e.m. across sessions. **(E)** Same as D but for RSC. **(F)** Evidence vs. choice single-neuron encoding analysis. Top: Regression equation. Bottom: Schematics of the firing across positions and evidence levels for a purely evidence encoding cell and a purely choice encoding cell on correct trials. **(G)** For each position, the average fraction of active neurons in each session in ACC (defined as neurons with fit position mean within one position standard deviation of the animal’s location) with significant coefficients for evidence (crimson) or choice (lime) in the choice vs. evidence encoding model, based on the F-statistic. Error bars indicate s.e.m. across sessions. The horizontal dashed gray line indicates chance level (1%). **(H)** Same as (C) but for RSC. **(I)** Top: Linear evidence decoding model at each position. Bottom: Example trials with actual (solid) and decoded (dashed) evidence in ACC. **(J)** At each position, the cross-validated correlation between actual evidence and predicted evidence from a linear population decoder in ACC on correct right evidence (*e* ≥ 0) trials (red) and left evidence (*e* ≤ 0) trials (blue), compared to shuffle (gray). Error bars indicate s.e.m. across sessions. **(K)** Same as (J) but for RSC. **(L)** Left: Heatmap showing average firing rates across all evidence-tuned neurons in ACC in bins of position by preferred-evidence. Gray indicates bins that were not sampled more than twice on at least 10% of the sessions. Right: Cross-sections of the heatmap at points in the cue period (50 cm, dark blue; 100 cm, medium blue; 150 cm, cyan) and delay period (200 cm, purple; 250 cm, magenta), showing average firing rate across evidence-tuned neurons as a function of preferred evidence. **(M)** Same as (A) but for RSC.

We next sought to quantify the mixture of graded evidence and binary choice activity in individual cells. To do so, we fit single-neuron evidence vs. choice encoding models at each position bin, using evidence and behavioral choice as regressors for both correct and incorrect trials (Fig. 6F, see Methods). At each position, we calculated the fraction of active neurons (see Methods) whose firing rate was significantly modulated by the linear evidence or the binary choice variable, while accounting for the contribution to neural activity of the other variable. In both ACC and RSC, we observed neurons with significant encoding of evidence beyond that explained by choice during the cue period, with decreasing proportions of cells encoding evidence during the delay period (crimson traces, Fig. 6G,H). In contrast, the fraction of significant choice encoding neurons (beyond that explained by the evidence) increases throughout the maze, with large fractions of active neurons encoding choice in the delay period (lime traces, Fig. 6G,H). The above analysis primarily differentiates behavioral choice-tuned neurons from evidence-tuned neurons by their change in tuning to evidence between correct and incorrect trials. However, a subtlety is that a neuron that has binary tuning to the “current best choice” (i.e., the current sign of the evidence), rather than graded tuning to evidence, may be identified as evidence tuned despite not being graded in its response. Therefore, to test whether there was graded response in the population of cells identified as evidence-tuned, we plotted the trial-averaged firing of the active cells in this subpopulation and confirmed that this subpopulation exhibited graded patterns of activity with evidence during the cue period, consistent with our model predictions for accumulation (Extended Data Fig. 13E,F).

To complement this single-neuron encoding analysis, we also asked whether evidence could be linearly decoded from the population (Fig. 6I). Similar to the findings from the single-neuron analysis, evidence can be linearly decoded from the population of active cells above chance early in the maze, falling to chance levels in the delay region (Fig. 6J,K). This is true even when controlling for the sign-of-evidence and considering only nonzero evidence levels to eliminate the possibility of a binary encoding of just the current sign-of-evidence (Extended Data Fig. 13G,H). Together, these results imply that graded evidence representations in the cue region transition to a more binary choice representation in the delay period.

We sought to summarize these results with a population level visualization. The population-averaged position-evidence responses in both ACC and RSC show graded increases in activity with preferred-evidence (i.e., evidence calculated in reference to each neuron’s preferred direction of evidence) during the early portion of the cue period (Fig. 6L,M, dark blue traces; population responses based on averaging all evidence-modulated neurons, see Methods). Consistent with the other analyses (Fig. 6G,H,J,K), with progression along the maze, this evidence tuning sharpens, transitioning to a step function by the start of the delay period (magenta and purple traces) that resembles tuning to choice rather than graded evidence tuning. In ACC (and more subtly in RSC), population responses also increase later in the maze, due to the fact that there are more evidence-tuned neurons in the delay region that contribute to these population averages (Extended Data Fig. 12A,B, left panels).

#### Augmenting the Circuit Models to Recapitulate the Transition from Graded Evidence to Binary Choice Tuning

Neither of the competing chains models produce this transition from relatively more evidence-encoding to relatively more choice-encoding, since the models were designed to perfectly accumulate evidence. This transition could be realized in the models in multiple manners. First, a population of choice readout cells at each position could compare the accumulation of the left and right chains at that position and respond in a binary fashion when either the left or right chain has greater activity, where the number of readout cells at each position increases over the course of the maze (Fig. 7 A-C). Second, this pattern could be realized if the parameters of the mutually inhibiting chains model are slightly mistuned such that the integrator becomes increasingly unstable later in the maze (Fig. 7 D-F). Such slight instability has been suggested in previous integrator models, where they have been functionally interpreted to provide an “urgency” signal that accelerates decision commitment^107,108^. These mechanisms are not mutually exclusive. Thus, the prevalence of choice cells in the data complements the integration processes focused upon in our models by suggesting that there are additional processes at play driving the transition from evidence accumulation to choice representation.

**Figure 7.**
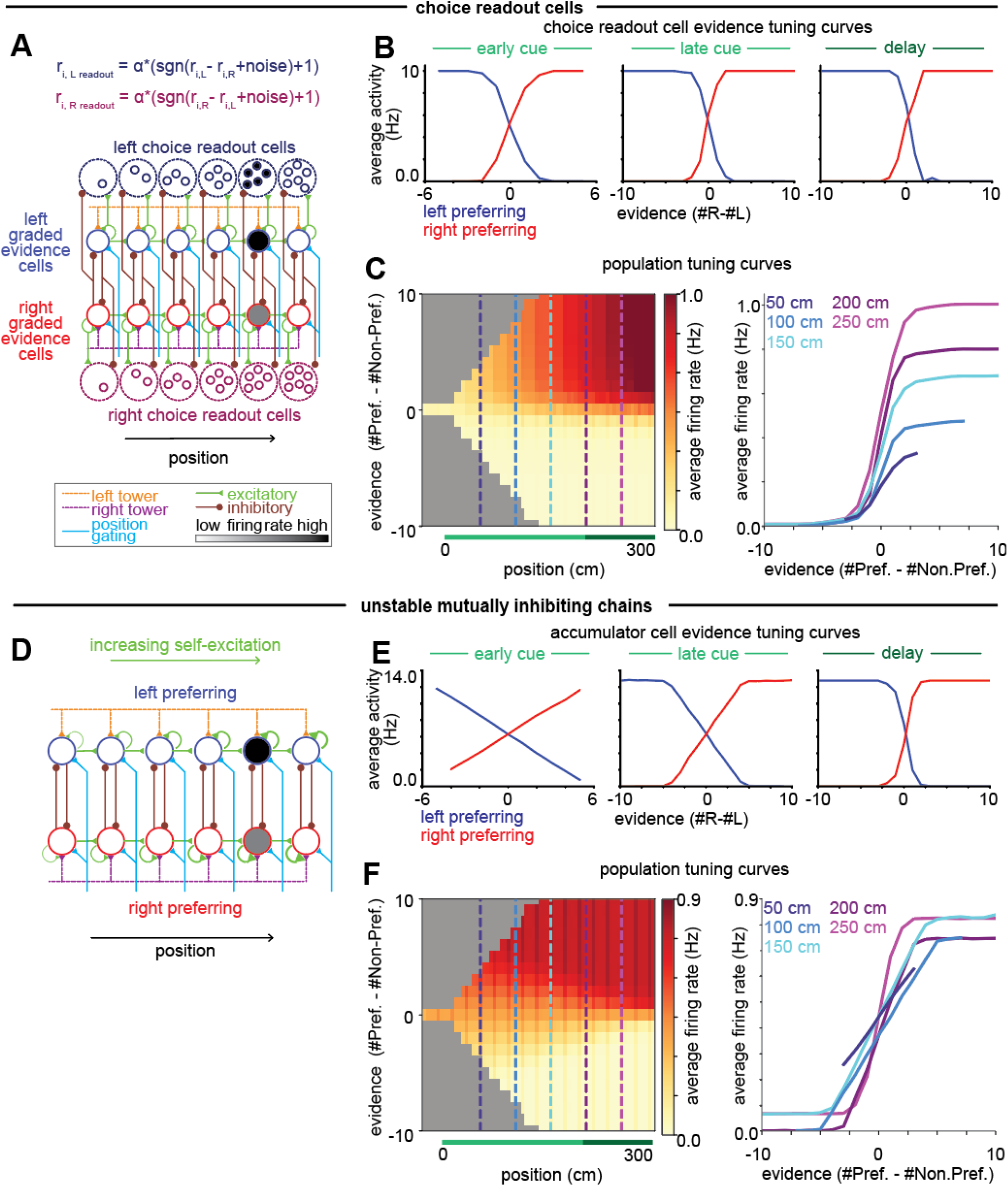
Simulated shifts towards choice tuning in the population. **(A)** Model schematic of a population of choice cells that readout from the accumulator and indicate if there is more left evidence (navy blue cells) or right evidence (dark magenta cells). The number of neurons in the choice readout population tuned to each position grows along the chain. **(B)** Tuning curves of the choice cells at their preferred position. **(C)** As in Figure 6, population average tuning curves of the entire population of evidence-tuned cells including both cells in the accumulator and the choice-readout cells. Cross sections show a shift towards choice tuning with increasing position in the maze. **(D)** Schematic of the mutually inhibiting chains model with unstable tuning, in which the strength of self-excitation increases with position along the chain. **(E-F)** Same as B-C but for cells in the unstable mutually inhibiting chains model.

## DISCUSSION

This work proposes mechanistic models for the widespread observation of choice-selective sequences during evidence accumulation (Fig. 1). Since traditional models of evidence accumulation based on persistent activity fail to explain choice-selective sequences (Fig. 2), we present two new classes of circuit models: “competing chains” models (Fig. 3) and “position-gated bump attractor” models (Fig. 4). These two model classes represent two different evidence accumulation schemes, in which evidence is encoded either in the amplitude of the neuronal firing rates or in the set of neurons that fire with stereotyped amplitude, and predict different effects of targeted perturbations. We tested the evidence tuning predictions in imaging data across multiple cortical and subcortical brain regions, demonstrating that although different regions show seemingly similar choice-selective sequences, their underlying evidence representations are distinct (Fig. 5, Fig. 6). Specifically, neocortical regions (ACC and RSC) had primarily monotonic evidence tuning curves in the early cue region, characteristic of the competing chains models, while the HPC showed non-monotonic evidence tuning with broadly distributed tuning curve peaks throughout the maze, consistent with the position-gated bump attractor.

### Transfer of Accumulated Evidence Across Positions is a Fundamental Cognitive Operation

Although our models were tested on data from a specific task, at the essence of accumulating evidence through sequences is the ability to both integrate evidence and transfer information between different populations of neurons. These two operations apply quite generally to many decision-making contexts. For example, position in our models should be interpreted broadly as a sequentially progressing state of the animal (e.g., it could represent time), so these models apply to navigation or non-navigation tasks. Moreover, perceptual decision making tasks may require transferring accumulated evidence across neuronal populations when the population that needs this information to carry out an action changes (e.g., changing the visual gaze position while maintaining evidence for two saccadic choice targets^109^). While models for evidence integration have been explored extensively and form the core of our models’ computations within a position, traditional models do not transfer graded information between positions, a novel feature in our models.

To achieve the transfer of information between positions, our models use a network architecture in which neurons project synaptically to neurons encoding the subsequent position, but these connections are only effective in transferring information when a position-gating signal pushes the receiving neurons above their firing threshold. While not explicitly modeled here, such spatio-temporal gating could result from inputs from the basal ganglia^110–113^, place cells in the hippocampus, or cells within each region that show position but not evidence tuning^44,49^ (Extended Data Fig. 1). The position gating signal in our model may itself result from an integration process, as in previous models of hippocampal path integration^86–88^, with the position integration performed in a separate population (Extended Data Fig. 2I) or within the same population (Extended Data Fig. 3) as the evidence integration.

### Despite Similar Choice-Selective Sequences, Evidence Coding Differs Across Regions

Despite similar choice-selective sequences, we found that evidence coding schemes across regions were surprisingly distinct. The neocortical regions primarily had monotonic encoding of evidence, as in the competing chains models, whereas the hippocampus had primarily non-monotonic encoding as in the position-gated bump attractor model. This result emphasizes the importance of examining how variables are represented, not just if they are represented, to compare computational mechanisms and functions across regions^114^.

The distinct evidence tuning properties we observed in hippocampus, ACC, RSC, and DMS support current ideas of their function in decision-making. The non-monotonic tuning curves in the hippocampus allow for a direct readout of the evidence based on the location of the active bump. This form of coding is more computationally intensive in the number of neurons required but could provide more refined information about the current state of the animal, supporting a joint cognitive map of position and evidence^48,115,116^, as suggested by previous analysis^49^. In contrast, the monotonic encoding of evidence in ACC, RSC, and DMS does not allow a direct readout of evidence from the identity of the active neurons. Instead, it may be ideally suited for making a unitary decision based on the comparison of activity along the two chains, and is similar to the graded coding that has been reported for value or confidence in ACC^117–123^. Interestingly, RSC, while dominated by monotonic tuning curves, did exhibit a population of narrowly tuned neurons. This may reflect that the hippocampus acts as a major input area to RSC, while RSC receives from and projects to other areas of cortex^124–127^, providing a possible suggestion as to how information about accumulated evidence is passed and transformed between brain regions. One possibility is that there is a single evidence-accumulation mechanism carried out in one set of brain regions, and other regions read out and transform this evidence into different tuning curve shapes. An alternative possibility is that both evidence accumulation mechanisms are used, but in different brain regions and for different purposes. Finally, the fact that DMS primarily responds in the delay region, where it mainly shows binary choice tuning (Extended Data Fig. 5), aligns with work suggesting the role of the striatum in action selection^128–130^, consistent with recordings in a different working memory task^55^.

In regions with primarily monotonic evidence tuning (ACC, RSC), we observed that this tuning was graded early in the maze and sharpened to more choice-like tuning later in the cue period and throughout the delay (Fig. 6). While the graded representations of evidence during the early cue period in ACC and RSC are consistent with the competing chains models of evidence accumulation, evidence accumulation models alone do not explain the transition towards more choice-like tuning later in the trial. This transition could reflect a change in dynamics as a result of decision commitment105,106 or increasing urgency^107,108^, and could be produced through many different mechanisms, including through a separate choice readout population or by making the integrator increasingly unstable (Fig. 7)^33^. The observation of a transition across the population from graded evidence coding to binary choice coding is reminiscent of the conclusions from a subpopulation of neurons in another accumulation of evidence task in rats^105^ and from the evolution across a trial of the power to predict choice from neural responses in monkeys^40^.

We applied our models to interpret neural activity in biological circuits, but the motifs and predictions of these models could also be tested for in trained recurrent neural network models. This could shed light on how less explicitly structured networks perform evidence accumulation^131^. A similar approach has been taken in the study of the path integration circuitry of the entorhinal cortex, where core foundational motifs of connectivity of the style presented here were revealed within the complex circuitry of the trained network^132^. Such studies also may reveal conditions favoring the emergence of evidence accumulation through sequences as opposed to traditional persistent activity, as has been analyzed for other cognitive operations^133^. Thus, tuning curve analysis in both biological and artificial networks may help to determine how these circuits perform the task.

### Functionally Targeted Optogenetic Perturbations Could Provide Causal Tests of Predicted Mechanisms of Evidence Accumulation

We were able to compare evidence tuning in new and existing datasets to those predicted by the models. However, unlike targeted perturbations, correlational analyses do not provide causal evidence for the underlying computations in a region. For example, they cannot distinguish neurons involved in accumulating evidence from those that simply readout evidence accumulated elsewhere. Further, within our set of models, they do not differentiate between the varying competing chains models, whereas optogenetic perturbations can (Fig. 3F, Extended Data Fig. 2F). Recent work, using electron microscopy, has suggested that the synaptic connections in the posterior parietal cortex provide a mutually inhibitory motif for evidence accumulation^134^, and these connections could be further probed through direct perturbations. Previous experiments have used optogenetic perturbations to identify brain regions that affect evidence accumulation at different times in the accumulating towers task^135,136^, but perturbation experiments that selectively target neurons with specific coding properties^71,137,138^ remain to be performed during evidence accumulation. Perturbations in the oculomotor neural integrator, where functionally distinct populations are conveniently anatomically separated, have successfully tested model predictions about the dynamics of the integrator^139–141^ and allowed the parameters of a model of this system to be fit directly from neural responses in perturbed and unperturbed trials^140^.

Targeted perturbations could provide insight into circuit mechanisms, and more broadly whether the evidence tuning observed in each region results from local evidence accumulation versus a downstream reflection of evidence accumulated in another region^14,142^. For example, evidence may be encoded in a traditional, non-sequential manner at some location and then combined at a downstream readout with a sequential representation of position to give the observed neural activity. If so, then an experimental prediction would be that activating choice-selective, evidence-encoding neurons during the cue region would not increase the firing rates of neurons tuned to later positions nor create a behavioral choice bias. Experiments such as these, combined with the mechanistic models and associated analyses presented in this work, should provide a foundation for elucidating how the neural signals guiding decision-making are accumulated within, and faithfully transferred between, neuronal populations.

## METHODS

### The Accumulating Towers Task

The accumulating towers task is a behavioral paradigm, performed in virtual reality, for studying accumulation-of-evidence based perceptual decision making. The complete details of the task and experimental setup can be found in Pinto et al. (2018)^59^. Here, we briefly summarize the key features of the task. Head-fixed mice run on an 8-inch styrofoam ball within a virtual environment, generated by ViRMEn^143^, projected onto a surrounding screen. The virtual environment has a T-maze structure. In the first 30 cm (the precue region), there are no visual cues. In the next 200 cm (cue region), visual cues appear as towers on both sides, with the total count of towers on each side determined by a Poisson distribution of different means. Towers are positioned randomly with an enforced minimal distance between them. A tower appears when the mouse is within 10 cm of its position and disappears 200 ms later. Following the cue region, there is a 100 cm delay region during which the mouse must remember which side was presented with more towers. At the end of the delay region, the mouse receives a sweetened liquid reward for turning to the side with more cues. For water restriction and shaping protocols, see Pinto et al. (2018)^59^. Recording sessions were approximately 1.0-1.5 h in duration, consisting of approximately 150-250 trials per session.

In addition to the accumulation of evidence trials described above, sessions also included some warm-up maze trials (cues on one side only, no distractors, tall visual guides at the end of the maze) at the beginning of the session and interspersed in the session if the animal’s performance dropped below some threshold, as described in Pinto et al. (2018)^59^. Since these trials do not require evidence accumulation, we did not include them in our analyses.

### Neural Recordings

Animal procedures were conducted in accordance with standards from the National Institutes of Health and under the approval of the Princeton University Institutional Animal Care and Use Committee.

The head-fixed setup of the accumulating towers task allows for neural imaging throughout the task. Our neural datasets consist of calcium fluorescence data previously recorded from retrosplenial cortex (RSC, 8 animals, 41 sessions (29 sessions from layers 2/3 and 12 sessions from layer 5), 8579 total neurons)^44^ and hippocampus (HPC, 7 animals, 7 sessions, 3144 total neurons)^49^, as well as previously unpublished recordings from anterior cingulate cortex (ACC, 3 animals, 7 sessions, 1720 total neurons) and dorsomedial striatum (DMS, 8 animals, 11 sessions, 804 total neurons). See below for details of ACC and DMS recordings.

#### Animals and Surgery for ACC Recordings

Three female mice of CaMKIIa-tTA (JAX 007004) crossed with tetO-GCaMP6s^144^ (JAX 024742) were used for two-photon imaging of ACC. These crossed lines show robust and stable expression of GCaMP6s in a subset of CaMKIIa-positive, glutamatergic neurons in the cortex. All mice were between 12-16 weeks when they underwent sterile stereotaxic surgery under isoflurane anesthesia (4% for induction, 1-1.5% for maintenance). The skull was exposed by making a vertical cut with a scalpel. Two craniotomy holes were made, one over ACC (anteroposterior axis: 0.8 mm, mediolateral axis: -0.5 mm, relative to bregma, diameter of ∼1.2 mm) and another over DMS (anteroposterior axis: 0.75 mm, mediolateral axis: -1.1 mm, relative to bregma, diameter of ∼0.3 mm). Adeno-associated virus (AAV; AAVretro2 serotype) encoding mCherry under CaMKIIa promoter (diluted to 2.5 × 10^12^ genome copies/mL, packaged at Princeton Vector Core) was injected to the DMS (anteroposterior axis: 0.75 mm, mediolateral axis: -1.1 mm, dorsoventral axis: -3.1 and -2.8 mm, relative to bregma) to provide information about a projection target and to be used for better motion correction. A total of 300 nL of AAV was infused at each site along the dorsoventral axis, at a rate of 30 nL/min. After injection, the needle was held in the same place for an additional 10 minutes. The needle was slowly withdrawn over 10 minutes to prevent backflow. After injections, the dura below the craniotomy was carefully removed within the ACC craniotomy site, and the cortical tissue above ACC was carefully aspirated up to 300 *μ*m below the surface. Then we implanted a 1.0-mm diameter GRIN lens (4.3 mm length, Inscopix, 1050-002184) into the ACC, down to 1.5 mm in the dorsoventral axis (relative to bregma), using a 3D-printed custom lens holder. After implant, a small amount of diluted metabond (Parkell) was applied to affix the lens to the skull using a 1-mL syringe and 25-gauge needle. After 10 minutes, the lens holder grip was loosened while the lens was observed through the stereoscope to make certain that there was no movement of the lens. After applying a more diluted metabond around the lens, a titanium headplate was positioned over the skull using a custom tool and aligned parallel to the stereotaxic frame using an angle meter. The headplate was affixed to the skull using a metabond. Finally, a titanium ring to hold the immersion medium was placed on the headplate, with lens in the middle, and then affixed to the headplate using dental cement blackened with carbon. After the dental cement was fully cured, mice were unmounted from the stereotaxic frame and protective head caps were placed. Recovery of the mice was monitored for ∼2 hours. After one or two weeks of recovery, mice started water-restriction, and after 5 days, behavioral training for the accumulating towers task began, as described in Pinto et al. (2018)^59^. Histology showed that GRIN lenses sampled mostly deep layers (layers 5 and 6) in two animals, while in one mouse a GRIN lens was medially implanted to cover both superficial (layers 2 and 3) and deep layers. For the data analysis, we did not make distinctions between neurons in different layers but instead pooled neurons from all layers together.

#### Animals and Surgery for DMS Recordings

D1R-Cre (n = 3, female, EY262Gsat, MMRRC-UCD), A2a-Cre (n = 3, male, KG139Gsat, MMRRC-UCD), and D2R-Cre (n = 2, male, ER44Gsat, MMRRC-UCD) mice were used for DMS two-photon imaging. Surgical procedures were identical to those described above for the ACC except for the following details. Approximately one week prior to GRIN lens implantation, a Cre-dependent AAV expressing GCaMP6f under either the CAG promoter (AAV5-CAG-DIO-GCaMP6f, Addgene, diluted to 3.5 × 10^12^ genome copies/mL) or synapsin promoter (AAV1-Syn-DIO-GCaMP6f, diluted to 3.8 × 10^12^ genome copies/mL) was delivered (500ul per injection site; 200 ul/min) to the DMS (anteroposterior axis: 0.7 mm, mediolateral axis: 1.15 and 1.85 mm, dorsoventral axis: -3 mm, relative to bregma) and the surgical incision was closed using monofilament non-absorbable sutures (Johnson and Johnson). This allowed for expression of GCaMP6f in D1R- or D2R-expressing medium spiny neurons according to the transgenic mouse line used, and facilitated the settling of brain tissue prior to GRIN lens implantation. Following recovery (5-10 days), sutures were removed and the viral injection craniotomy was expanded and dura removed to accommodate a 1.0-mm diameter GRIN lens with a prism (4.3 mm length, Inscopix, 1050-004601). The lens was implanted with prism window facing posterior and just anterior to viral injection sites (anteroposterior axis: 0.75 mm, mediolateral axis: 1.35 mm, dorsoventral axis: -3.25 mm, relative to bregma). Fixation of the lens to the skull, as well as the custom-printed headplate, water immersion ring, and protective covering were identical to that described above for ACC. Across all DMS targeted GCaMP6f imaging mice we obtained three distinct fields of view from one mouse, two distinct fields of view from a second mouse, and the remaining mice contributed single fields of view (8 animals, 11 sessions, 804 total neurons).

#### Optical Imaging and Data Acquisition for ACC and DMS Recordings

Imaging was performed using a custom-built two-photon microscope compatible with virtual reality (VR). The microscope was equipped with two high-power femtosecond lasers (Alcor, Spark Laser), with fixed wavelengths for 920 and 1064 nm. The scanning units used a 5-mm galvanometer and a 8kHz resonant scanning mirror (Novanta). The collected photons were passed to a dichroic mirror (XYZ) that separates near-infrared light from the visible light and then split into two channels by another dichroic mirror (FF562-Di03, Semrock). The light for respective green and red channels was filtered using a bandpass filter (FF01-520/60 and FF02-607/70, Semrock), and then detected using GaAsP photomultiplier tubes (1077PA-40, Hamamatsu). The signal from the photomultiplier tubes was amplified using a high-speed current amplifier (59-179, Edmund). Output of the amplifier was sent to a data acquisition system (National Instruments). Black rubber tubing was attached around the objective (Zeiss, W N-Achroplan 20x/0.5 M27, 20x magnification, 0.5 NA) to shield the light from the virtual reality screen. The objective was aligned to the implanted GRIN lens within the implanted titanium headring. Double-distilled water was filled inside the headring and used as the immersion medium. The amount of laser power at the objective was 30∼40 mW for ACC and 40∼50 mW for DMS. Control of the microscope and imaging acquisition was performed with the ScanImage software (MBF Bioscience), on a separate computer from the VR-running one. Images were acquired at 30 Hz with a size of 512 × 512 pixels. Synchronization between behavioral logs and acquired images was achieved by sending behavioral information from the VR controlling computer to the scanning computer via an I2C serial bus each time the ViRMEn environment was refreshed. Specifically, numeric information about the current session, trial, and frame were written in the header of TIFF imaging files.

#### Data Processing for ACC and DMS Recordings

Both ACC and DMS imaging data were processed with Suite2p. For ACC recordings in which mCherry signals are present, rigid motion correction was performed with that channel and then applied to the functional green channel with GCaMP6s, with the following parameter settings: nimg_init = 1000, smooth_sigma = 1.15, maxregshift = 0.1. For DMS recordings, the same motion correction was performed, but with the green channel. Regions-of-interest (ROIs) corresponding to individual neurons were detected, and calcium signals were extracted and neuropil-corrected (neucoeff = 0.7). Subsequently, an experimenter-mediated curation was performed on these ROIs to discriminate putative neurons from noise. Neuropil-corrected fluorescence from these curated neurons was exported to MATLAB, and the ΔF/F was calculated, taking baseline fluorescence as the mode of the fluorescence within 3 minute long windows (the same baseline as applied in the RSC and HPC datasets).

#### Spike Inference

In order to compare neural recordings across datasets with different calcium indicators, we performed spike inference on our data based on a recently developed and validated algorithm that has only two free parameters, the average estimated firing rate, and the calcium decay rate^145,146^. For each ΔF/F dataset, we assumed a baseline firing rate of 2 Hz (but our results were not sensitively dependent on this specific choice) and set the calcium decay rate according to the values fit by Jewell et al. (2020)^145^ for each calcium indicator (0.9885 for GCaMP6s (ACC), 0.9704588 for GCaMP6f (all other regions)). From these inferred spikes, we inferred a continuous firing rate by smoothing with a gaussian window function (window length = 1 s, standard deviation = 0.25 s).

#### Position Binning of Neural and Evidence Data

To compare neural data along maze positions and evidence levels, we resampled the neural (continuous firing rate as calculated above) and behavioral (cumulative evidence up to that point) data in position bins. Specifically, firing rates were averaged based on the position of the animal within 5 cm bins from the start of the trial (−30 cm) to the end of the delay region (300 cm) for each trial. Cumulative evidence for each bin was defined by the number of right minus the number of left cues observed before the start of the position bin.

#### Trial Selection Criteria

We discarded a small fraction of trials from analysis using the following criteria. As mentioned above, any trials from the warm-up maze were excluded, as these trials do not require evidence accumulation. Additionally, trials with excessive travel were discarded (3.2% of trials in ACC sessions, 8.0% of trials in DMS sessions), which were identified as trials where mice showed a large view angle deviation (more than 90 degrees, either left or right) before the end of the delay region.

### Determining Choice Selectivity of Neurons

To identify choice-selective neurons (Fig. 1B-E), we first jointly fit the position and evidence tuning of each neuron in our dataset as described in Data Analysis below. We then considered only neurons that had significant evidence tuning. To produce the sequence plots, neurons were first sorted as right-preferring if the mean of the neuron’s evidence tuning was greater than 0 and left-preferring otherwise and then sorted by their position mean. Using this sorting, we then took the average activity of each neuron at different positions for correct left and correct right trials. The resulting traces for each neuron were then peak normalized.

### Traditional Integrator Models Simulations

To show the activity of traditional integrator models (Fig. 2), we simulated equations corresponding to previously developed models of integration: a competing accumulator model and a bump attractor model.

#### Competing Accumulator Model

We modeled two competing populations of neurons, one left-preferring population and one right-preferring population. The activity *r*_*L*_ of the neuron representing the left population is given by

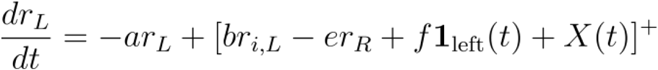

and a corresponding equation governs the right population, where

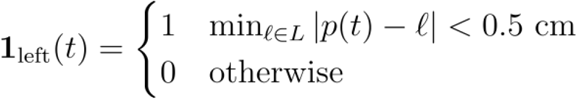

and

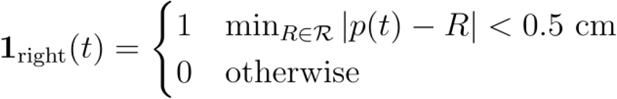

where *p*(*t*) is the position of the animal at time *t, L* is the set of positions of left cues, and ℛ is the set of positions of right cues.

In our simulations, we used parameters *a* = 50 s^−1^, *b* = 10 s^−1^, *e* = 40 s^−1^, *f* = 100 s^−1^, and, common external input to both populations, *X*(*t*) = 500 Hz/s. The model was simulated as described in Simulation of Differential Equations.

#### Bump Attractor Model

For the bump attractor model, we assumed each neuron has its peak response at a different evidence level. The activity of the neuron with peak response at evidence *j* is given by

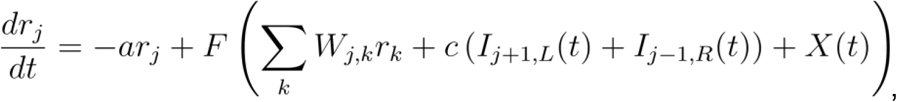

where

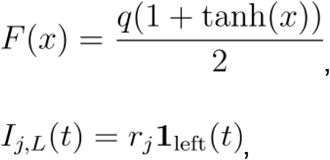

and

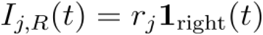

where

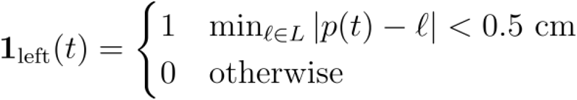

and

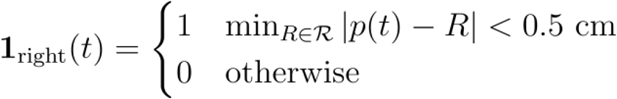

where *p*(*t*) is the position of the animal at time *t, L* is the set of positions of left cues, and ℛ is the set of positions of right cues.

We assigned the neuron at each evidence level an angle, evenly spaced between 0 and 2*π*, and set the synaptic connection weights as

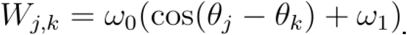

We note that, since evidence is not periodic, a sufficiently large number of evidence levels were used such that substantial interactions do not occur between the nominally periodically connected endpoints.

For the shown simulation, we used 35 neurons, corresponding to evidence levels of -17 to 17. We used the parameters *a* = 55 s^−1^, *X*(*t*)=0.04 Hz/s, *q* = 1250Hz/s, *ω*_*0*_ = 0.12 s^−1^, *ω*_*1*_ = -1.0, and *c* = 0.2 s^−1^. Neurons *r*_*-1*_, *r*_*0*_, and *r*_*1*_ were initialized to 15 Hz, 17.5 Hz, and 15 Hz activity, respectively, and all other neurons were initialized to 0 Hz activity. The model was simulated as described in Simulation of Differential Equations.

### Parameterization of the Mutually Inhibiting Chains Model

As in Eq. (1), the activity of the neuron at position *i* in the left chain can be described as

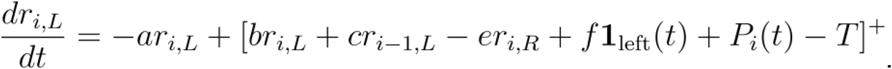

A neuron’s active position is defined by a square pulse over a range of positions, where *p(t)* is the position of the animal at time *t*,

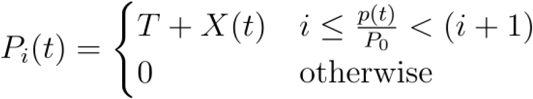

The external input is of the form

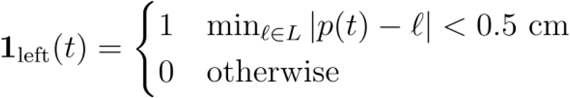

And

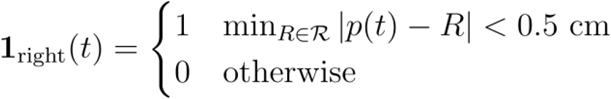

where *L* is the set of positions of left cues and ℛ is the set of positions of right cues.

In our simulations, we used parameters *a* = 50 s^−1^, *b* = 10 s^−1^, *c* = 50 s^−1^, *e* = 40 s^−1^, *f* = 100 s^−1^, *T* = 15000 Hz/s, and *X*(*t*) = 500 Hz/s. (See Supplementary Text for necessary conditions on the parameters.) We take *P*_*0*_ *=* 20 cm, which divides the total length of the maze (330 cm) into 17 different sections of active positions. Therefore, in these models we simulate 17 neurons in each chain for a total of 34 neurons. This choice of parameters was selected to recapitulate typical firing rates of the neurons that we recorded as well as approximate the typical position width observed in the experimental choice-selective sequence plots (Fig. 1).

The parameters above produce saturation in the evidence tuning at the extreme evidence levels (Fig. 3E). As demonstrated in Extended Data Fig. 2J, for different parameters, this model will instead show linear tuning curves over the observed range of evidence (same parameters as above, but *X(t)* = 1000 Hz/s). We note that a similar effect of linear evidence tuning throughout the observed evidence range could have been achieved by decreasing the weight *f* on the cues. Both manipulations prevent the non-dominant chain from reaching 0 Hz, which results in saturation of the dominant chain.

In Extended Data Fig. 2G, we changed the form of the position gate to be a smooth gaussian, rather than a square pulse. To do so, we used the original parameters but took T = 1100 Hz/s and

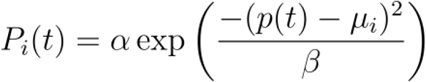

where *α* = 60 Hz/s, *β* = 1250 cm^2^, and *μ*_*i*_ = 20*i* - 10 cm.

In Extended Data Fig. 2H, we change the form of the position gate to have heterogeneous widths. We use the original parameters but only 12 neurons in each chain and rather than a fixed width *P*_*0*_ for each position, we draw a set of transition points *ρ*_*i*_ uniformly between -30 cm and 300 cm. If (*ρ*_*i+1*_ - *ρ*_*i*_) *<* 5 cm, we take *ρ*_*i+1*_ = *ρ*_*i+1*_ + 5 cm, to enforce a minimum position width of 5 cm. We then define

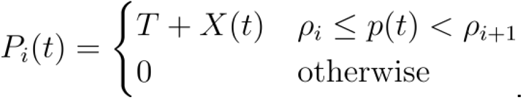

### The Mutually Inhibiting Chains Model for the Transition to Choice Representations

In Figure 7, we propose two models that could explain the transition from more graded evidence representations early in the maze to more choice-like representations later in the maze. The first model (Fig. 7A-C) incorporates a population of choice-readout cells. In this model, the evidence accumulator cells of the chains are modeled exactly as described in the previous section, but we introduce a population of choice readout cells at each position. The firing rate of a left choice readout cell at position i, *r*_*i,L,readout*_, is given by

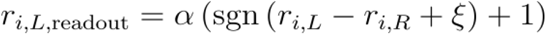

where *α* = 5 Hz, sgn is the signum function which returns the sign of its input, and *ξ ∼* 𝒩 (*0, 2*) Hz with the analogous equation for a right choice readout cell. We assume that there are *i* choice readout cells for each choice at position *i*.

In the second model (Fig. 7D-F), we set the parameters of the mutually inhibiting chain model such that the integrator becomes increasingly more unstable with advancing position down the chain. Specifically, we incrementally increase the weight of self-excitation *b* in Eq. 1 with progression down the chain by making it a function of the position *i*:

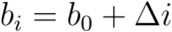

where *Δ* = 0.5 s^−1^, and *b*_*0*_ = 10 s^−1^ is the value of self-excitation that gives a perfect integrator.

### Parameterization of the Position-Gated Bump Attractor

In Eq. (2), we defined the evolution of the firing rate for a neuron at position *i* and evidence *j* as

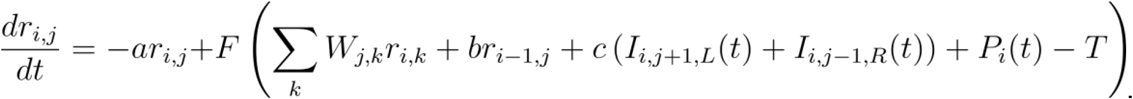

A neuron’s active position is defined by a square pulse over a range of positions, where *p(t)* is the position of the animal at time *t*,

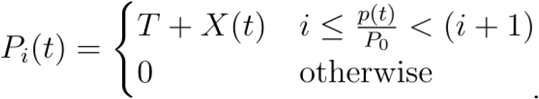

The neuronal nonlinearity is

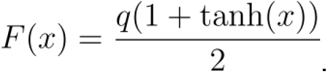

The strengths of the synaptic connections between neurons were chosen to be symmetric with excitatory connections between neurons nearby in their evidence tuning and inhibitory connections between neurons farther apart in their evidence tuning. This was done by assigning each evidence level an angle, evenly spaced between 0 and 2*π*, and setting

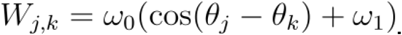

We note that, since evidence is not periodic, a sufficiently large number of evidence levels were used such that substantial interactions do not occur between the nominally periodically connected endpoints.

When an input is present, asymmetric connections cause the active evidence neurons to shift towards the input. This is modeled through shifter neurons (Fig. 4A, purple and orange small circles) that are activated by the coincidence of external cue signals and neural activity from the current location of the bump by

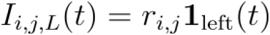

and

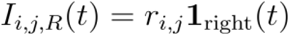

where

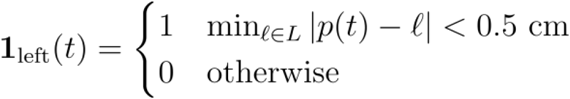

and

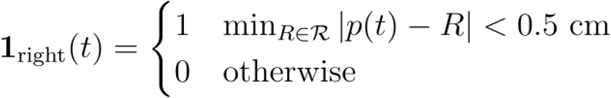

where *p*(*t*) is the position of the animal at time *t, L* is the set of positions of left cues and ℛ is the set of positions of right cues.

As in the competing chains models, the position-gating of each layer of neurons is activated over a length 20 cm (*P*_*0*_ = 20), again giving 17 different active positions. For the neurons to span the range of evidence of the experiments, we used 35 neurons in the evidence dimension, corresponding to evidence levels of -17 to 17. Thus we simulated a total of 595 neurons.

The parameters used in the simulation are: *a* = 55 s^−1^, *b* = 0.088 s^−1^, *T =* 300 Hz/s, *X*(*t*)=0.04 Hz/s, *q* = 1250 Hz/s, *ω*_*0*_ = 0.12 s^−1^, *ω*_*1*_ = -1.0, and *c* = 0.2 s^−1^. Neurons *r*_*0,-1*_, *r*_*0,0*_, and *r*_*0,1*_ are initialized to 15 Hz, 17.5 Hz, and 15 Hz activity, respectively, and all other neurons are initialized to 0 Hz activity.

### Simulation of Differential Equations

Differential equations for each model (Eqs. (1), (2), and (S2)) were simulated in Python using scipy.odeint, which uses the lsoda algorithm, with a maximum stepsize of 0.01 s (or equivalently 0.5 cm when assuming a constant velocity of 50 cm/s) to find the activity levels of the neurons in each model for points between -30 and 300 cm in 0.1 cm increments. In all simulations, we assume the animal travels at a constant velocity of 50 cm/s through the maze so that time and position are interchangeable. We simulated the neural activity for 1000 trials with cue positions derived from 1000 experimental trials from the behavioral and neural recording data. For the competing chains models, we say that the animal makes a left choice if the activity of the last neuron in the left chain is greater than the activity of the last neuron in the right chain. For the bump model, we say that the animal makes a left choice if the index *k* of the neuron with maximal firing in the last layer is less than 0.5*J*, where *J* is the total number of neurons in the layer. If the maximal firing occurs exactly at 0.5*J*, we determine the animal’s decision by randomly choosing between the two alternatives with equal probability.

#### Simulations with Input Noise

For each model, we simulated 25 sessions with input noise (Figs. 3C, 4C, Extended Data Fig. 2B). For each session, we selected 150 trials drawn randomly without replacement from the 1000 experimental trials that we used for simulation of the model. Let *L* and *R* be the set of left cues and right cues, respectively, in the original trial. For each cue *x* in *L*, we drew *z*, a random uniform number between 0 and 1, and if *z*<0.33, we added *x* to the set *L*’. We repeated the same process for each cue in *R*, yielding two reduced sets of cues when input noise is present, *L*’ and *R*’, corresponding to the animal ignoring or not perceiving 67% of the cues. We then simulated the model with the input terms ***1***_*left*_(*t*) and ***1***_*right*_(*t*) determined according to the reduced cue sets. We then calculated the psychometric curves (as described in Psychometric Curves below) with the difference in cues based on the original cue sets, but whether the animal turns left or right based on the simulation of the trial with the reduced cue sets.

#### Evidence Tuning Curves for Simulated Model Data

In order to find the evidence tuning curves (Figs. 3E, 4E, Extended Data Fig. 2D,J,K), we averaged the activity of each neuron binned by position and accumulated evidence for each of the 1000 simulated trials. Position bins were of size 0.1 cm, ranging from -30 to 300 cm. Evidence bins were of size 1 tower, ranging from -15 to 15 towers. Bins that had fewer than 2 samples were not included in further analysis and not plotted. For a given neuron, we averaged across evidence levels to find the position with maximum firing. We then plotted the average firing rate versus evidence level at this position.

#### Simulated Single-Neuron Perturbations

Single-neuron perturbations were simulated in the same manner as no-perturbation trials but in the absence of any cues. A subthreshold excitatory term *O*_*i*_(*t*) was added within the thresholded dynamics of each model (i.e. within the square brackets of Eq. (1) or within the function *F* of Eq. (2)) to simulate the effects of stimulating an individual neuron, where *O*_*i*_(*t*) = 25 Hz/s for the mutually inhibiting competing chains model, *O*_*i*_(*t*) *=* 12.5 Hz/s for the uncoupled competing changes model, and *O*_*i*_(*t*) = 0.2 Hz/s for the position-gated bump attractor, if neuron *i* is the neuron being stimulated, and 0 otherwise.

### Psychometric Curves

For each session, we sorted trials based on the difference in the final number of cues at the end of the maze. We binned these trial differences with bin size of 3 towers, ranging from - 14 to 14 towers. Within each bin, we then found the fraction of trials in each bin for which a left choice was made (Fig. 3C, Fig. 4C, Extended Data Fig. 2B).

### Data Analysis

#### Determining Position and Evidence Selectivity

In our neural recording data, we sought to identify the set of neurons whose firing encoded accumulated evidence. As described in the main text, due to the nonuniform sampling of position and evidence and fact that neurons are only active at some positions, we found that robustly identifying evidence selectivity required that we jointly identify the position range over which a neuron is active and the evidence selectivity within this set of positions. To do so, we assumed the firing rate of each neuron had the form,

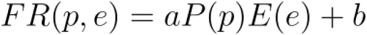

where *FR(p,e)* is the firing rate of the neuron at position *p* and evidence *e*, and *P* and *E* were described by Gaussians,

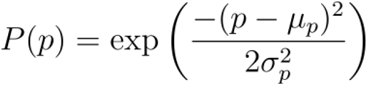

and

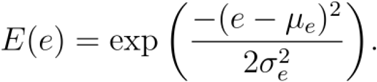

The above functional form, which assumes evidence and position tuning can be approximated by independent Gaussians, was not used as a detailed descriptor of neuronal firing (for less constrained fits around a given position, see Neural Evidence Tuning Curves Around the Most Active Position). Rather, we found that it was effective for the purpose of identifying evidence-modulated neurons and determining if the mean evidence selectivity favors left or right evidence (as used to sort neurons by evidence-selectivity in Fig. 1B-E). Here, the Gaussian was allowed to extend beyond the range of observed evidence values, so that it can accommodate both monotonic and non-monotonic tuning curves (i.e., if the mean evidence occurs at the extremes of the evidence range, the evidence tuning will be monotonic throughout the range, Extended Data Fig. 4D).

For each neuron, we normalized the firing rate observations (binned as in Position Binning of Neural and Evidence Data) to the range [0, 1]. We then fit the parameters of the above model using an iterative fitting procedure, alternating fitting *μ*_*p*_, *σ*_*p*_, *a*, and *b* with fitting *μ*_*e*_, *σ*_*e*_, *a*, and *b* with a nonlinear fitting method (scipy curvefit, which employs the trust region reflective algorithm, with bounds -50 ≤ *μ*_*p*_ ≤ 350, 0 ≤ *σ*_*p*_ ≤ 200, *e*_*low*_ - 1 ≤ *μ*_*e*_ ≤ *e*_*high*_ + 1, 0 ≤ *σ*_*e*_ ≤ 30, 0 ≤ *a* ≤ 10, and 0 ≤ *b* ≤ 1). The bounds on *μ*_*e*_ are defined by *e*_*low*_ = min(*E’*) and *e*_*high*_ = max(*E’*), where *E’* is the set of all evidence levels observed by the neuron within 5 cm of *μ*_*p*_. These bounds are updated on each iteration based on the current fit of *μ*_*p*_. Due to the iterative nature of our fitting procedure, after the final position update, *μ*_*e*_ may no longer satisfy these bounds, and we do not include such neurons when analyzing the distributions of the normalized evidence parameters.

We then tested for significance of the evidence tuning fit using the pseudosession method^147^. For each session of firing rate and position data, we generated new evidence for each trial in the session by resampling trials from actual sessions, and applied the iterative fitting procedure to the new pseudosession evidence and position data. We repeated this process for 50 pseudosessions and calculated the mean squared error between the true firing rates and the model predictions for each session. We then used a one sample t-test to compare the fit of the model, as measured by the mean squared error, in the true session to the fit of the null distribution from the pseudosession, and only considered neurons for which the fit was significantly better (*p<0*.*05*) than the null distribution for further analysis.

In order to compare the parameterization of the evidence tuning across positions (Fig. 5D,H,L, Extended Data Fig. 5D), we normalized *μ*_*e*_ and *σ*_*e*_ to account for the different observed evidence ranges at different positions. Let *ε* be the set of all evidence values observed by the neuron at positions within 5 cm (one position bin) of *μ*_*p*_. Define *ε*^*+*^ = {*e* ∈ *ε* | e > 0}, i.e. the set of positive observed evidence levels, and *ε*^**-**^ = {*e* ∈ *ε* | e < 0}. For *μ*_*e*_ < 0, the normalized mean is given by

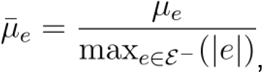

and for *μ*_*e*_ > 0, the normalized mean is given by

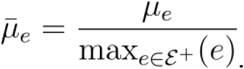

The normalized evidence standard deviation is given by

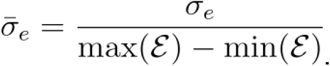

For visualization purposes (Fig. 5, Extended Data Fig. 5), we plot only the 80% of significantly evidence-selective neurons in each region with the best fit. For robustness to single large observations making a neuron appear narrowly tuned, for neurons with *σ*_*e*_ < 3, we identified the nearest position-evidence bin to (*μ*_*p*,_ *μ*_*e*_). If there were fewer than 4 observations in this bin or if there was a single outlier in this bin (defined by one observation greater than 1.5 times the interquartile range), the neuron was excluded from plotting. For visualization of all significantly evidence-selective neurons without any further criteria applied, see Extended Data Fig. 12. In all density plots, kernel density estimates are generated using the python package seaborn, which uses scipy.stats.gaussian_kde to estimate the density with Gaussian kernels (bandwidth determined by Scott’s Rule).

#### Neural Evidence Tuning Curves Around the Most Active Position

To generate the 1-D evidence tuning curves around the most active position (Fig. 5B,F,J, Extended Data Fig. 5B) for each neuron with significant evidence tuning, we identified the subset of data within 0.5*σ*_*p*_ of *μ*_*p*_. (Note that if this range extended to less than 0 cm or greater than 300 cm, only the data within 0-300 cm were used.) Using the data from within this range, the tuning curves were computed by computing the average of the firing rates at each evidence level. We then fit the averages to two functional forms, a Gaussian and a logistic function, and plotted the curve that better fit the average tuning curve. By fitting to the averages, rather than the individual data points, we found we could better account for the uneven distribution of sampling across evidence levels at different positions (Extended Data Fig. 4A). The Gaussian evidence tuning curve fit took the form

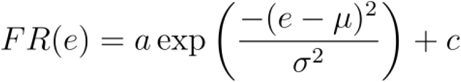

where *FR(e)* denotes the firing rate as a function of the evidence *e*. The fit was performed with a nonlinear fitting method (scipy curvefit, which employs the trust region reflective algorithm, with bounds *m* < *μ* < *M*, 0 < *σ* < 30, 0 < *a* < 10, 0 < *c* < 1, where *m* is the minimum and *M* is the maximum observed evidence within the active position). The logistic evidence tuning curve took the form

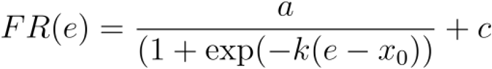

and was fit with a nonlinear fitting method (scipy curvefit, which employs the trust region reflective algorithm, with bounds -1 < *k* < 1, -15 < *x*_*0*_ < 15, 0 < *a* < 10, 0 < *c* < 1). To determine the curve with the better fit, we calculated the mean squared error between the predicted firing rates from each fit curve and the true firing rates. We note that each form of the tuning curve has the same number of fit parameters, so there was no need to correct for the numbers of parameters when comparing model predictions.

For comparison to our fit tuning curves, we also plotted the raw evidence tuning curves around the most active position (Extended Data Fig. 6). We note that such raw evidence tuning curves around a given position require a reasonable assessment of the peak position tuning, as was obtained in our joint position-evidence fits described above. However, once the most active positions were obtained, the fits no longer depended on requiring the evidence to have the same functional form at every position or, in the case of the raw fits, on a particular functional form. Thus, we found that the fits could provide a more quantitatively precise visualization of the evidence tuning around the most active position and, for the neurons best fit by logistic functions, a better quantitative assessment of the steepness of the sigmoid. However, beyond this small quantitative difference, the fits typically did not differ substantially from the joint Gaussian fits, and required fitting to two separate functional forms, so we used the joint Gaussian fits for the population-wide assessment of tuning in Figure 5C,D,G,H,K,L.

#### Cue Encoding Model

To find the average response to a cue (Fig. 6A-E), we fit a linear cue encoding model to the difference between the average right-preferring activity and average left-preferring activity on each session, using the subset of neurons that are significantly evidence tuned:

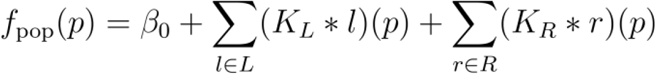

where *f*_*pop*_(*p)* is the difference between the average activity of the right evidence-preferring cells and the average activity of the left evidence-preferring cells at position *p, L* is the set of left cues, R is the set of right cues, and *K*_*L*_ and *K*_*R*_ are the cue response kernels to left and right cues respectively. As above, neurons that were significantly evidence tuned were categorized as right preferring if they had *μ*_*e*_ > 0 and left preferring if *μ*_*e*_ < 0.

For this analysis, we filtered the inferred spikes from the Ca^++^ imaging data using a causal half-Gaussian (window length = 1 s, standard deviation = 0.25 s), so that responses to cues would be purely causal. In order to fit the shapes of the cue response kernels of width 300 cm, each cue was convolved with a cubic spline basis set with 7 degrees of freedom^148^. We fit a ridge regression model with regularization strength 1 to predict the difference in population activity. The regression used the convolution of the cues with each of these splines as regressors, so that the cue encoding model was given by

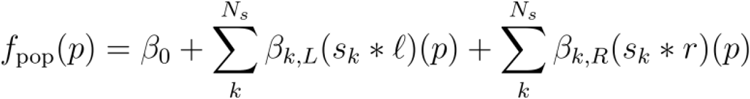

where *s*_*k*_ is the *k*^th^ basis function, *N*_*s*_ = 7 is the number of basis functions, and *ℓ* and *r* are binary vectors with value 1 at the onset (defined as the time at which the cue becomes visible, 10 cm ahead of its position) of each left and right cue respectively.

Left cues and right cues were convolved separately to derive separate cue response kernels for each cue type. In Extended Data Figure 13A,B, separate cue kernels were fit for left and right cues when the magnitude of the current evidence prior to cue appearance, *e*, was low (|*e*| ≤ 1), medium (2 ≤ |*e*| ≤ 4), or high (|*e*| ≥ 5). For the medium and high evidence cues, the width of the cue kernel was 200 cm. In Extended Data Figure 13C,D, separate cue kernels were fit for left and right cues when the cue occurred in the early (before 70 cm), middle (between 70 and 140 cm), or late (after 140 cm) part of the cue period. For early cues, the width of the cue kernel was 300 cm, for middle cues, the width of the cue kernel was 230 cm, and for late cues, the width of the cue kernel was 160 cm.

#### Single-Neuron Evidence vs. Choice Encoding Analysis

For each neuron at each position bin *p*, we fit a single-neuron evidence vs. choice encoding model (Fig. 6F-H) to describe the firing rate of the *i*^th^ neuron

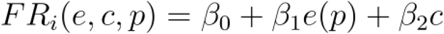

as a function of the current evidence *e*(*p*) at that position and the upcoming behavioral choice on that trial (*c*), using ordinary least squares regression. For this analysis, we fit to data from both correct and incorrect trials. For each coefficient, we calculated the F-statistic, and used the F-test (one-sided) to determine whether a coefficient was significant at a 0.01 significance level. At each position, for each session, we found the fraction of active neurons at that position that had a significant coefficient. For this analysis, a neuron was defined as active at position *p* if |*p - μ*_*p*_| < *σ*_*p*_, based on the results of the joint position-evidence fitting procedure described above.

#### Average Evidence Population Responses for Trials with Different Evidence Levels

In Extended Data Figure 13E,F, we examined the average population responses across positions for correct trials with different final evidence levels. We binned trials by final evidence for left and right trials for low (1 ≤ *e* ≤ 3), medium (4 ≤ *e* ≤ 6), and high evidence levels (*e* ≥ 7). At each position bin, we only analyzed neurons that met two criteria: (1) they were active at this position bin, where active at position *p* is defined as above as having |*p - μ*_*p*_| < *σ*_*p*_, and (2) had a significant evidence coefficient *β*_*1*_ (at a 0.01 significance level) at any position bin, based on the results of the single-neuron encoding model described above. For these neurons, we calculated the difference between the average firing rates (extracted from the Ca^++^ imaging using the causal half Gaussian filter, as described above) of the right-preferring and left-preferring populations. As above, a neuron is defined as right preferring if *μ*_*e*_ > 0 and left preferring if *μ*_*e*_ < 0.

#### Linear Evidence Decoding Model

To evaluate the ability to linearly decode evidence from the population (Fig. 6I-K), we fit two decoders at each position bin, one for trials in which the evidence was greater than or equal to 0 and one for trials for which the evidence was less than or equal to zero. Separate decoders were calculated so that positive correlations between actual and decoded evidence do not result just from correctly decoding the animal’s choice on correct trials. To further eliminate correlations resulting from predicting just the sign-of-evidence, we also applied the same analysis but with one decoder for evidence strictly greater than zero and one for evidence strictly less than zero (Extended Data Fig. 13G,H). For each decoder, we used nested five-fold cross-validation^149^ to fit a ridge regression model to predict evidence from the activity of the active neurons at that position. As above, neurons were defined as active at position *p* if |*p - μ*_*p*_| < *σ*_*p*_. In the inner loop of cross-validation, the regularization strength was chosen from [.0001, .001, .01, .1, 1, 10, 100, 1000], with the best performing regularization strength used for evaluation on the test set in the outer loop. Decoders were evaluated by the average correlation between the predicted and actual evidence in the test set. To compare the performance to chance, we repeated the same fitting procedure but randomly shuffled the evidence values across trials of the same sign-of-evidence for each position. We repeated this analysis for 5 shuffles.

#### Nonlinear Evidence Decoding in HPC

In Extended Data Figure 11, we use a nonlinear manifold method^103^ to decode evidence from the neural population. On each fold of a five-fold cross validation, trials were assigned to training or test sets. To maximize the amount of data used to build the embedding, both correct and incorrect trials were used. We then fit a CEBRA manifold^103^ (https://cebra.ai/) embedding of dimension *d* to the training set for values of *d* from 2 to 7, using the following parameter settings: model_architecture = ‘offset10-model’, batch_size = 512, learning_rate = 3e-4, temperature_mode = ‘auto’, output_dimension = d, max_iterations = 10000, distance = ‘cosine’, conditional = ‘time_delta’, device = ‘cuda_if_available’, time_offsets = 10. At each point, we provided the current evidence value as the label for the neural data. We trained a *k* nearest neighbors decoder (k = 3) to predict the evidence level from the embedded training set. We then embedded the test set into the CEBRA manifold and used the decoder to predict evidence from the embedded test set. We then found the Pearson correlation between the predicted and actual evidence levels on the test set. To compare performance to chance, we repeated this process for 5 shuffles of the data in which the neural data was preserved but the evidence labels were permuted across trials within trials of the same behavioral choice.

#### Population-Averaged Tuning Curves

In Figure 6L,M, we calculated the population-averaged response across all significantly evidence-selective neurons as follows. For each neuron, we calculated the average firing rate in each position-evidence bin. If a bin had fewer than 3 observations for a particular neuron, that neuron was not included in the population average for that bin. To combine data for neurons that preferred positive evidence with data from neurons that preferred negative evidence, for each neuron, we defined its preferred-evidence as the sign of the neuron’s *μ*_*e*_ value and remapped the evidence axis to be in terms of preferred evidence rather than absolute evidence (by multiplying the evidence values by -1 if the preferred tuning was for leftward values of evidence instead of rightward). We then took the average across all neurons in each bin of position by preferred-evidence. Bins which were not sampled three or more times on at least 10% of the sessions for a particular brain region were not plotted.

## Data Availability

Data from HPC is previously published and available^49^. Data from RSC is previously published and available^66^. Data from ACC and DMS will be made available upon publication.

## Code Availability

Code for the simulation of all models and the data analysis is available at https://github.com/lindseysbrown/evidence_accumulation_through_sequences.

## Acknowledgements

We thank Sue Ann Koay for her useful discussions about the retrosplenial cortex dataset. We thank Shizhe Chen, Tim Hanks, Ben Lankow, and Avinash Baidya for useful discussions about this project. Funding was provided by NIH T32MH065214 (LSB), F32MH132179 (LSB), NIH-NINDS BRAIN Initiative 5U19NS104648 (LSB, DWT, CBD, IBW, MSG) and 1U19NS132720 (LSB, DWT, CBD, IBW, MSG), K99DA053388 (EHN), Sloan Swartz Foundation (LSB), C.V. Starr Fellowships (JRC, MS), and Burroughs Wellcome Fund CASI awards (LSB, MS).

This manuscript is the result of funding in whole or in part by the National Institutes of Health (NIH). It is subject to the NIH Public Access Policy. Through acceptance of this federal funding, NIH has been given a right to make this manuscript publicly available in PubMed Central upon the Official Date of Publication, as defined by NIH.

---

**Extended Data Figure 1.**
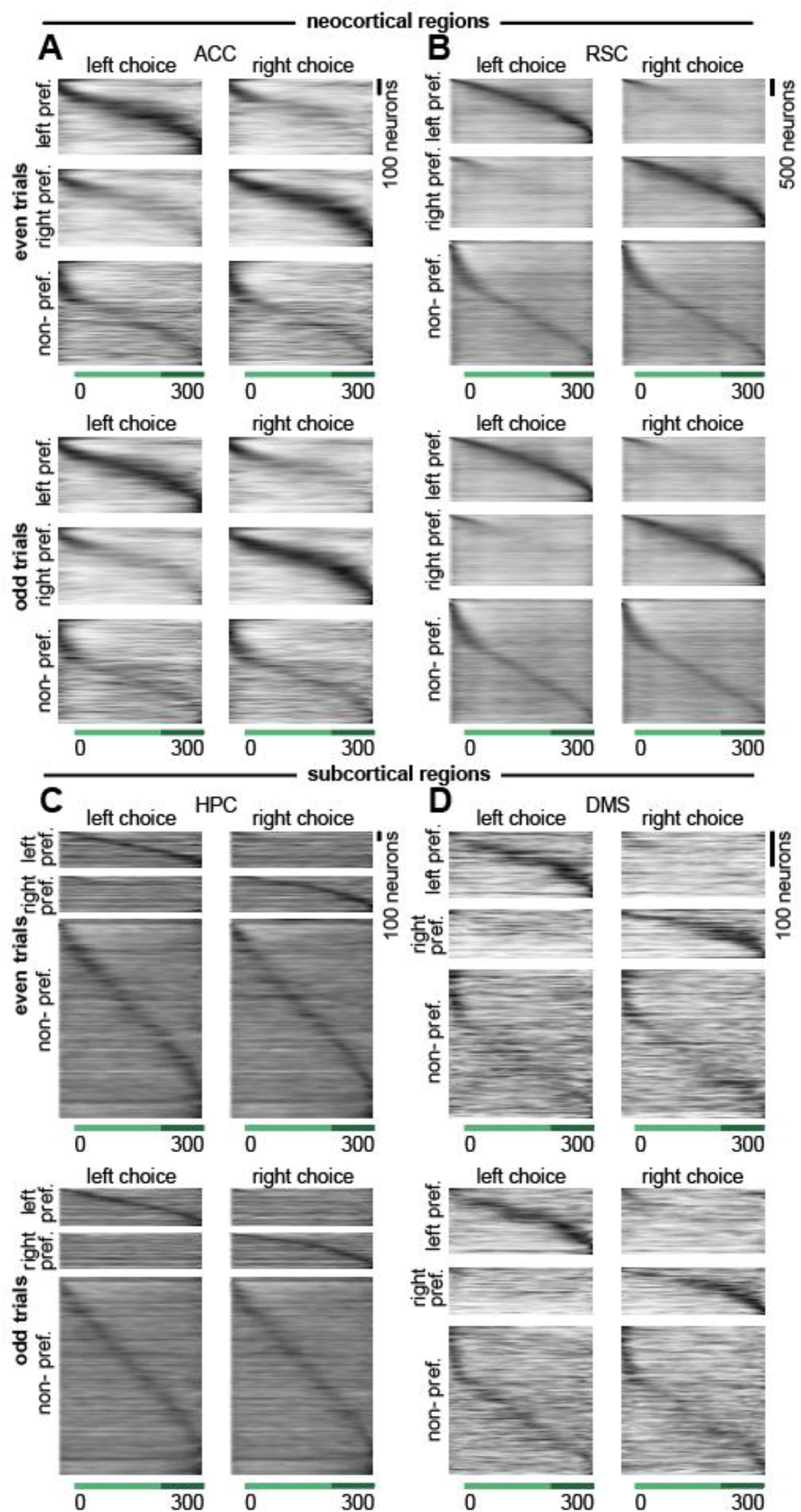
Choice-selective sequences across brain regions during the accumulating towers task appear similar on even and odd trials. **(A)** Each row shows the peak-normalized averaged firing rate (normalization based on the odd trials, with this odd-trial-based normalization then applied for cross-validation purposes to the neuronal responses on even trials) at each position in the maze of a neuron with significant evidence tuning (see Methods) recorded during the accumulating towers task from ACC (n = 1720 neurons), averaged across even trials (top) or odd trials (bottom) when the animal turned left (left choice, left column) or right (right choice, right column). Neurons were divided based on their choice-selectivity (see Methods) and ordered based on the position of peak activity. **(B-D)** Same as (A) but for RSC (n = 8579 neurons; B), HPC (n = 3144 neurons; C), and DMS (n = 804 neurons; D).

**Extended Data Figure 2.**
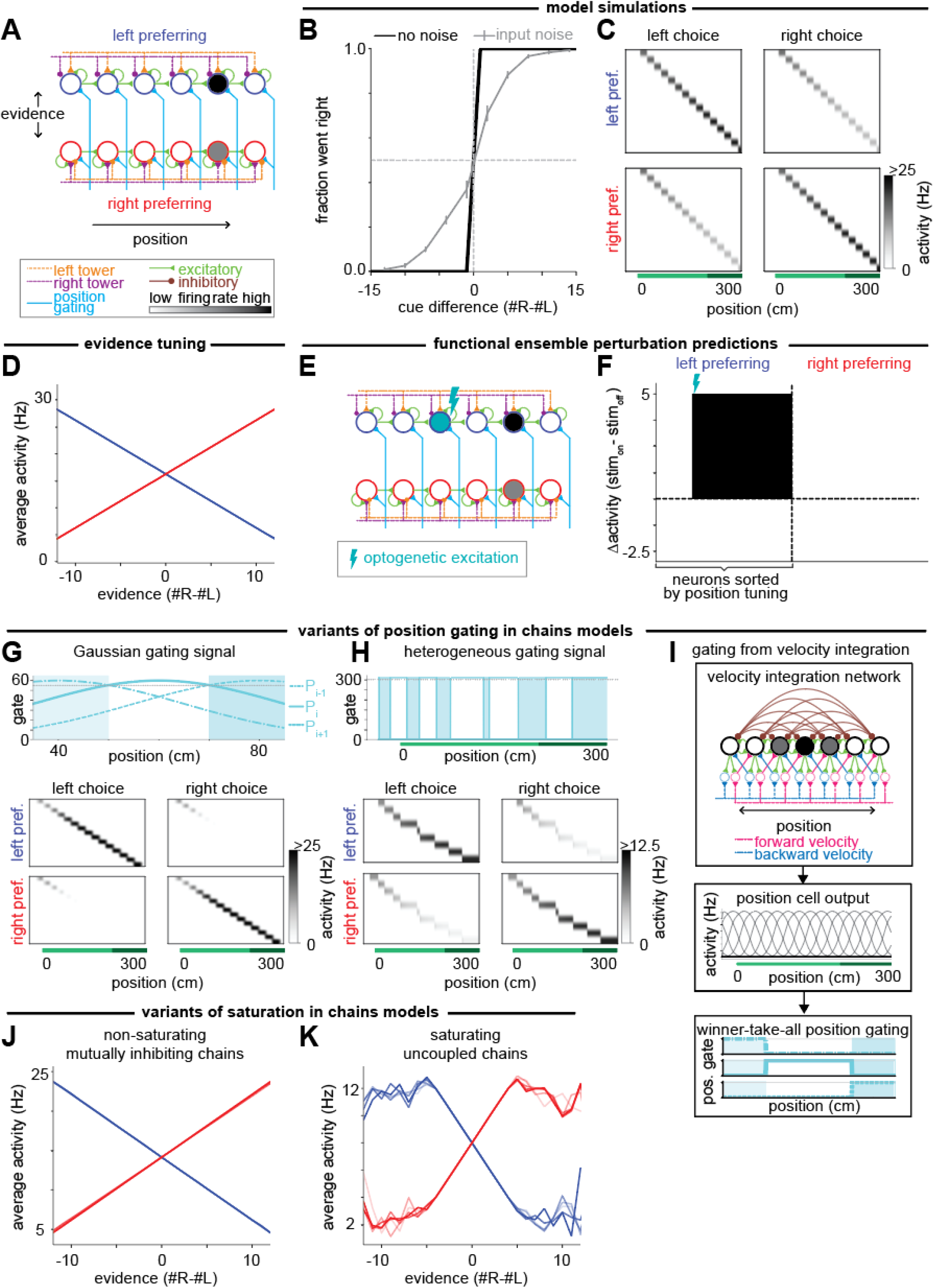
Variants of competing chains models of evidence accumulation through sequences. **(A)** Schematic of neural circuit architecture for the uncoupled competing chains model, showing excitatory (green) connections between neurons (circles) as well as inputs to the circuit from the external left (orange) and right (purple) towers and a position gating signal (cyan). **(B)** Psychometric data for model simulated trials describing how often the amplitude of the final neuron in the left chain was greater than that of the final neuron in the right chain for cases in which the model was simulated with (gray) and without (black) noise in the input. Error bars show s.e.m. **(C)** Each row shows the non-normalized amplitude of a model neuron at each position in the maze, averaged across simulated trials without input noise when the greater final amplitude was in the left (left choice, left column) or right (right choice, right column) chain. Neurons were divided based on their choice-selectivity (see Methods) and ordered based on the position of peak activity. **(D)** Tuning curves of individual neurons to evidence, defined by the average activity for different evidence levels at the neuron’s peak position, for left- preferring (blue) and right-preferring (red) neurons. **(E)** Schematic of a single neuron perturbation experiment in which optogenetic excitation is applied to a single neuron in the left chain. **(F)** Simulated changes of the firing rates of all neurons in the absence of cues when a single cell (denoted with the laser as in E) is optogenetically stimulated. **(G)** Variant of the mutually inhibiting chains model with a Gaussian position gating signal (top). Neuronal responses (bottom) are as in panel C. **(H)** Same as G but for a square position gating signal of heterogeneous width. **(I)** Schematic of a velocity integration method for generating square position-gating signals, where velocity is integrated through a bump attractor to produce cells representing different preferred positions. A winner-take-all network takes these position cells as input to produce a square position gate. **(J)** As in D, but for a parameterization of the mutually inhibiting chains model which does not saturate within the observed evidence range. **(K)** As in D, but for a variant of the uncoupled competing chains model where an upper bound for saturation is enforced.

**Extended Data Figure 3.**
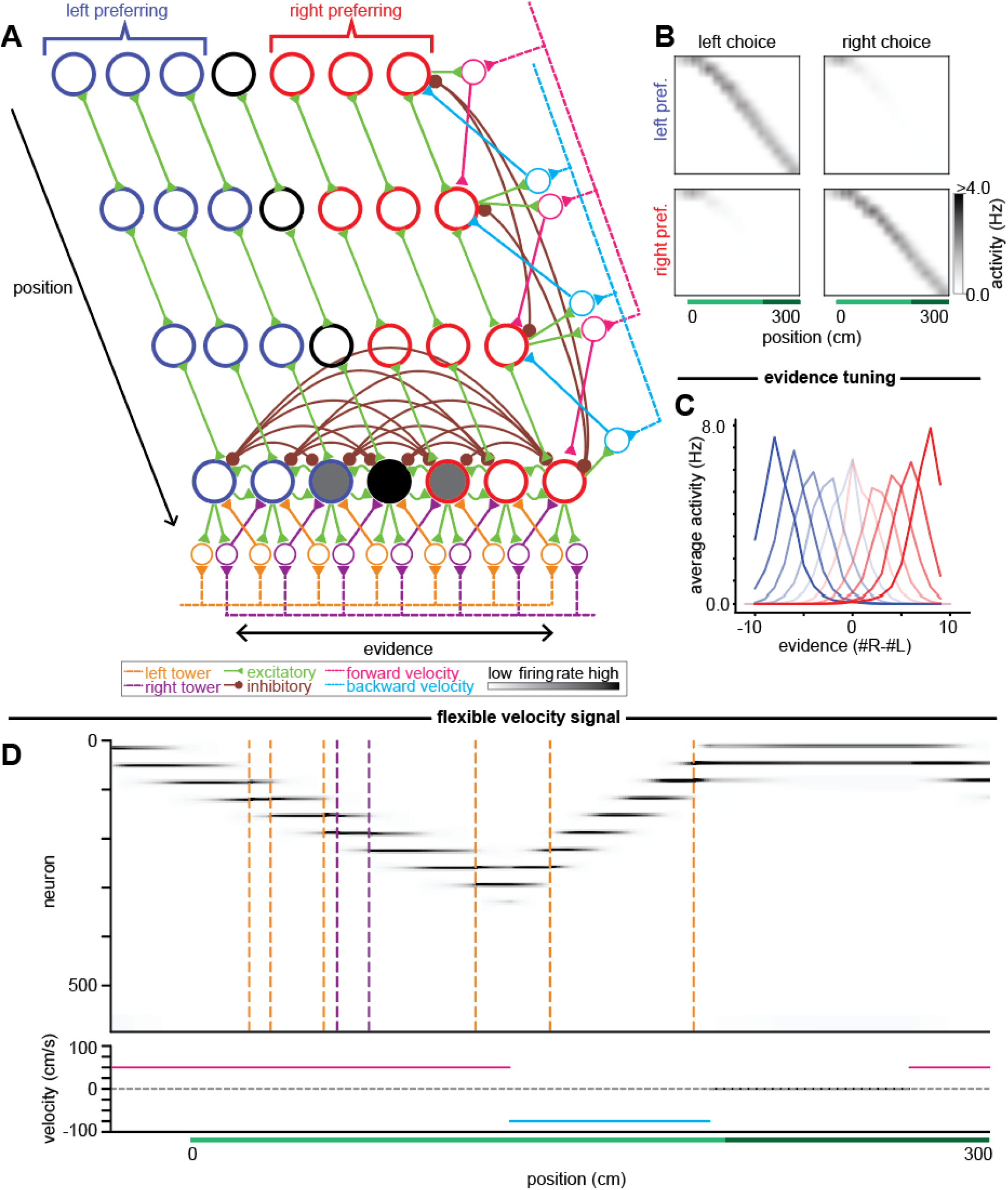
Planar bump attractor that jointly accumulates evidence and position through sequences. **(A)** Schematic of the neural circuit architecture for the planar bump attractor. Each row represents neurons (circles) that respond at a given position. Within a row, each neuron represents a different evidence level, ranging from left-most to right-most. Blue: left-preferring neurons; red: right-preferring neurons. As in the position-gated bump attractor, the bottom row of neurons illustrates the cue-related inputs and connectivity within any given row: local excitatory (green) and broader inhibitory (brown) connections between neurons as well as external inputs to the circuit from the left (orange lines) and right (purple lines) towers via the corresponding shifter neurons (orange and purple circles). The rightmost column illustrates the velocity-related inputs and connectivity within any given column: local excitatory and broader inhibitory connections between neurons at different positions as well as either forward-direction (pink) or backward-direction (cyan) external velocity inputs via corresponding position-shifter neurons (pink and cyan circles). **(B)** Each row shows the non-normalized firing rate of a model neuron at each position in the maze, averaged across simulated trials without input noise when the greater final amplitude was in the left (left choice, left column) or right (right choice, right column) chain. Neurons were divided based on their choice-selectivity (see Methods) and ordered based on the position of peak activity. **(C)** Tuning curves of a subset of individual neurons to evidence, calculated when the neurons have peak position activity, for left-preferring (blue) and right-preferring (red) neurons. **(D)** Example simulation of the planar bump attractor for a theoretical trial where the animal can move both forward and backward. Top: Raster of activity of individual model neurons across different positions in the trial. Incoming towers are indicated by the dashed orange (left cue) and purple (right cue) lines. Neurons are ordered first by position and then by evidence within a given position. Bottom: The velocity of the animal at different points of the trial, including forward (pink) and backward (cyan) movement.

**Extended Data Figure 4.**
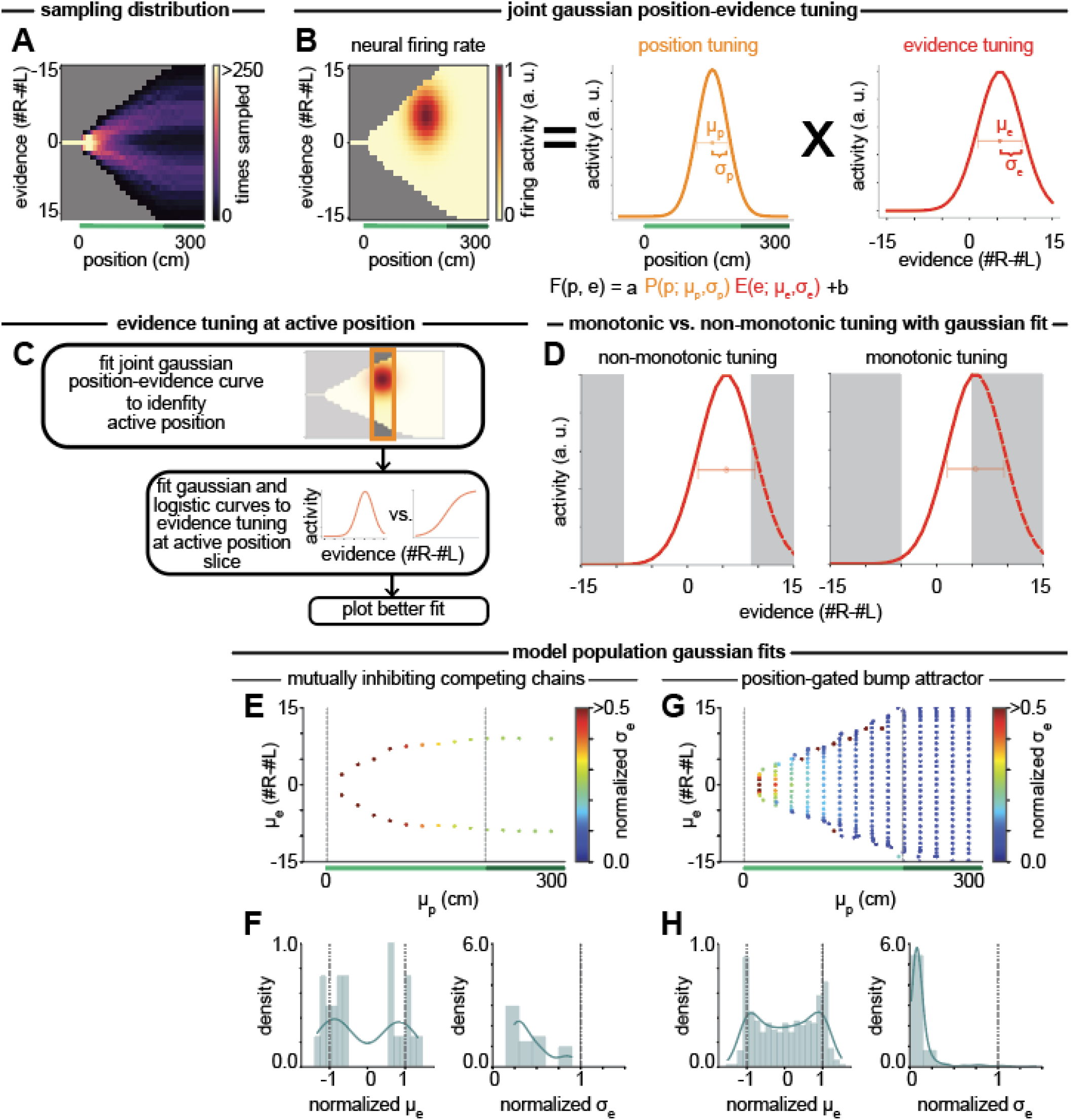
Joint position-evidence tuning curve fitting procedure and Gaussian fits to the different models. **(A)** Number of times each position-evidence bin is sampled within the 1000 empirical trials used to simulate the models. The statistics of cue counts, distribution, and position used to generate trials make it such that at different positions, different evidence levels are not sampled uniformly. For example, neurons tuned to the beginning of the maze only see a small range of small evidence levels at the positions where they are most active; such a cell may not appear to respond to high levels of evidence because its position tuning is never active at these extreme evidence levels. **(B)** Fitting firing rates to a joint position-evidence Gaussian to identify *μ*_*p*_, *σ*_*p*_, *μ*_*e*_, and *σ*_*e*_. **(C)** Schematic of procedure for plotting evidence tuning curves by first identifying the active position region and then comparing the fit between a Gaussian and logistic curve. **(D)** Examples of how Gaussian fits can capture non-monotonic (left) and monotonic (right) tuning to evidence, with the monotonicity of the fit curve dependent on the observed evidence range (gray regions indicate non-observed evidence levels). **(E)** Scatter plot showing the location of the fit mean position (*μ*_*p*_) and fit mean evidence (*μ*_*e*_) with color indicating the normalized fit evidence standard deviation (*σ*_*e*_) (see Methods) of simulated neurons in the mutually inhibiting competing chains model. Dashed lines indicate the boundaries of the cue period. **(F)** Density plots of the normalized fit mean evidence (left) and the normalized fit evidence standard deviation (right) for the neurons shown in (E). **(G-H)** Same as for (E-F) but for simulated neurons from the position-gated bump attractor.

**Extended Data Figure 5.**
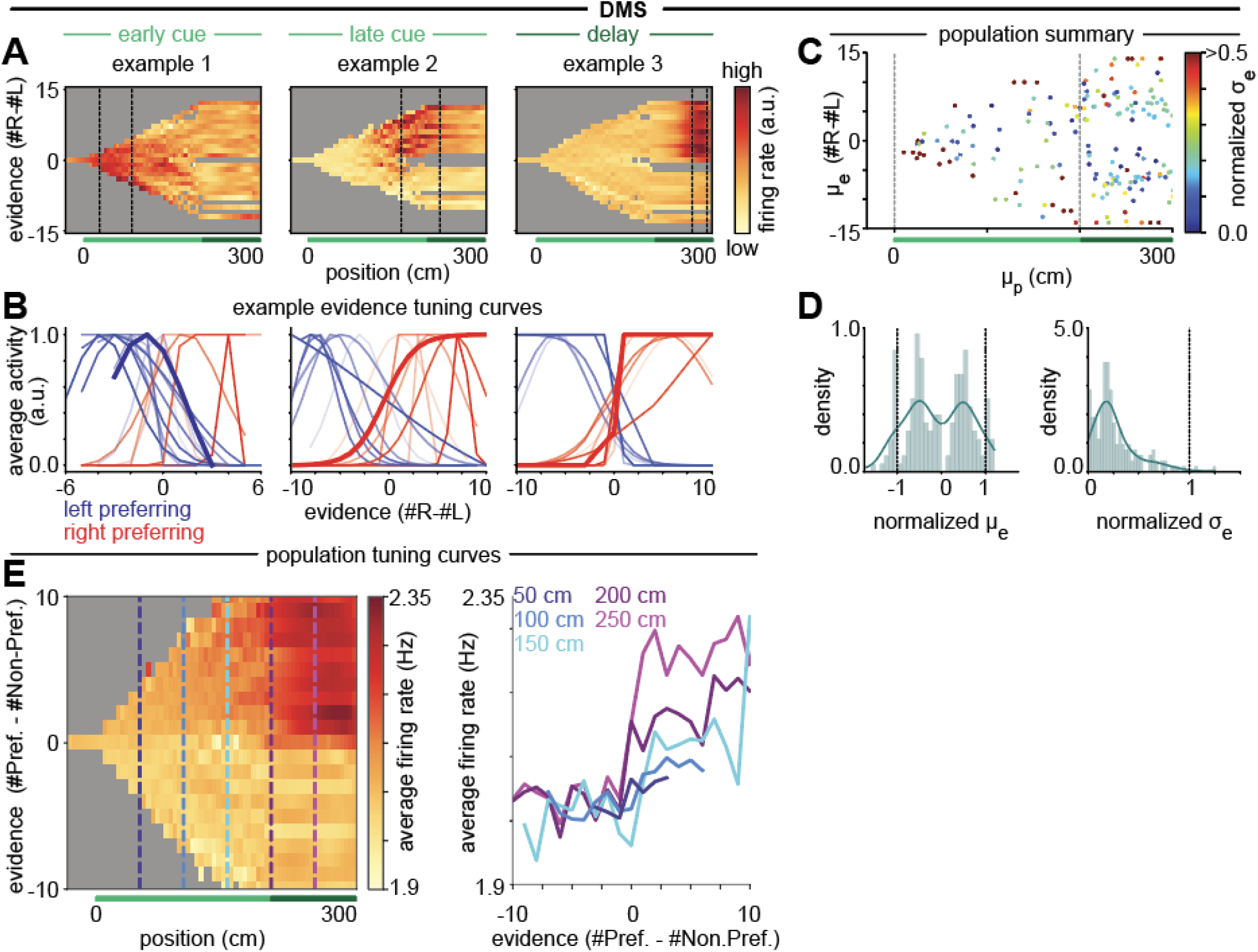
DMS exhibits mainly monotonic evidence tuning dominated by neurons with position tuning in the delay region. **(A)** Heatmaps showing the average firing in position by evidence bins for example individual neurons in DMS with mean position in the early cue (left), late cue (middle), or delay (right) region of the maze. Gray bins denote regions for which there were fewer than 2 samples during the session. **(B)** Example DMS evidence tuning curves fit to the region of the neuron’s peak activity (see Methods) for a collection of neurons with mean position tuning in the early cue (left), late cue (middle), or delay (right) region of the maze. Red coloring indicates neurons classified as right-preferring, and blue indicates left-preferring. Bold lines correspond to the examples in (A), for which the neuron’s region of peak activity is the region between the dashed vertical lines. **(C)** Scatter plot showing the location of the fit mean position (*μ*_*p*_) and fit mean evidence (*μ*_*e*_) with color indicating the normalized fit evidence standard deviation (*σ*_*e*_) (see Methods) of the 80% of neurons recorded in DMS with the best fit between the neural data and the model predictions. Dashed lines indicate the boundaries of the cue period. **(D)** Density plots showing the normalized fit mean evidence (left) and the normalized fit evidence standard deviation (right) for the neurons shown in (C). **(E)** Left: Heatmap showing average firing rates across all evidence-tuned neurons in bins of position by preferred-evidence. Gray indicates bins that were not sampled more than twice on at least 10% of sessions. Right: Cross-sections of the heatmap at points in the cue period (50 cm, dark blue; 100 cm, medium blue; 150 cm, cyan) and delay period (200 cm, purple; 250 cm, magenta), showing average firing rate across evidence-tuned neurons as a function of preferred evidence.

**Extended Data Figure 6.**
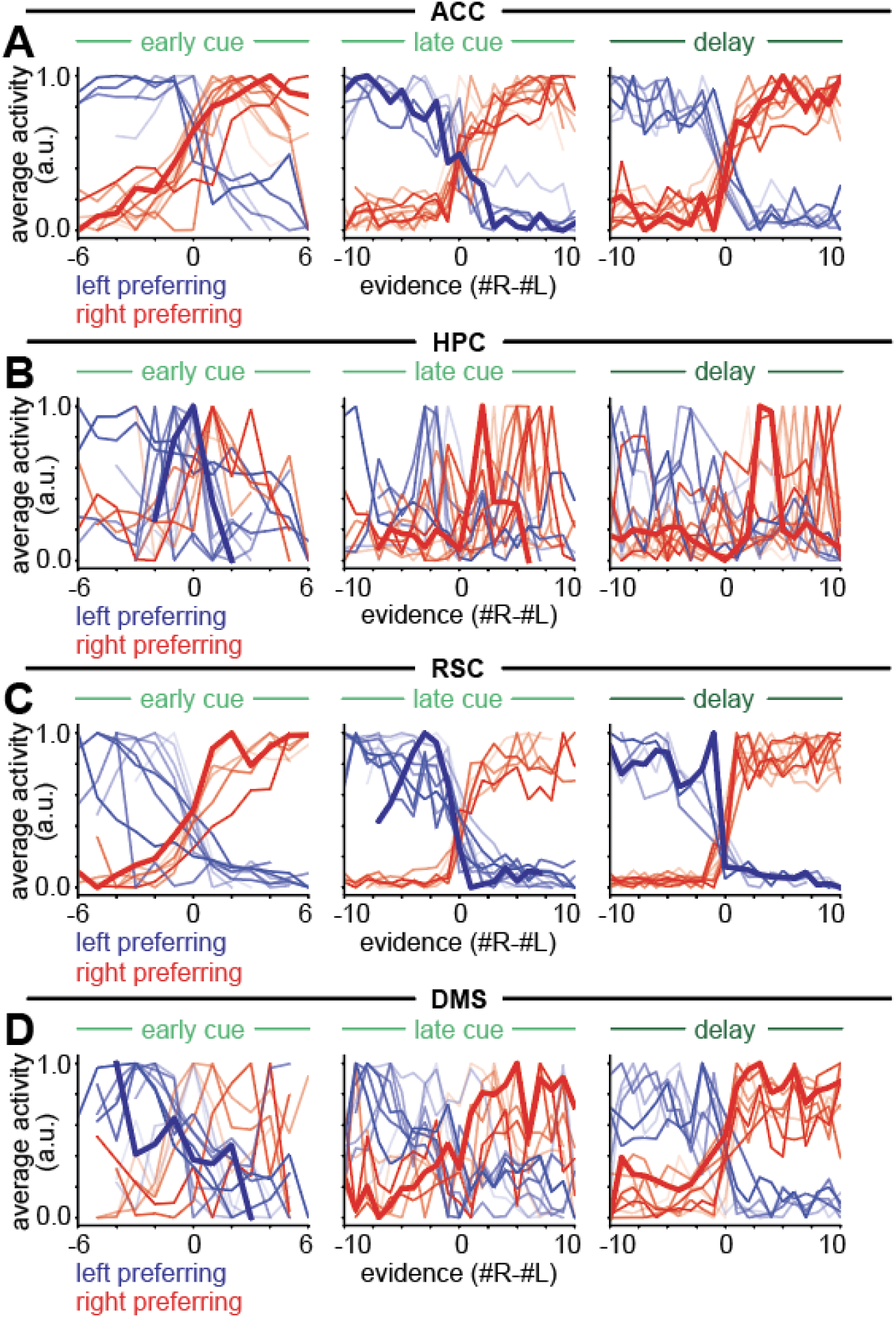
Raw tuning curves for the four recorded brain regions. **(A)** Average activity of example ACC neurons at different evidence levels in the region of the neuron’s peak activity (see Methods) for a collection of neurons with mean position tuning in the early cue (left), late cue (middle), or delay (right) region of the maze. Red coloring indicates neurons classified as right preferring, and blue indicates left preferring. Bold lines correspond to the examples plotted in Figure 5. **(B)** Same as (A) but for HPC. **(C)** Same as (A) but for RSC. **(D)** Same as (A) but for DMS, with the examples from Extended Data Fig. 5.

**Extended Data Figure 7.**
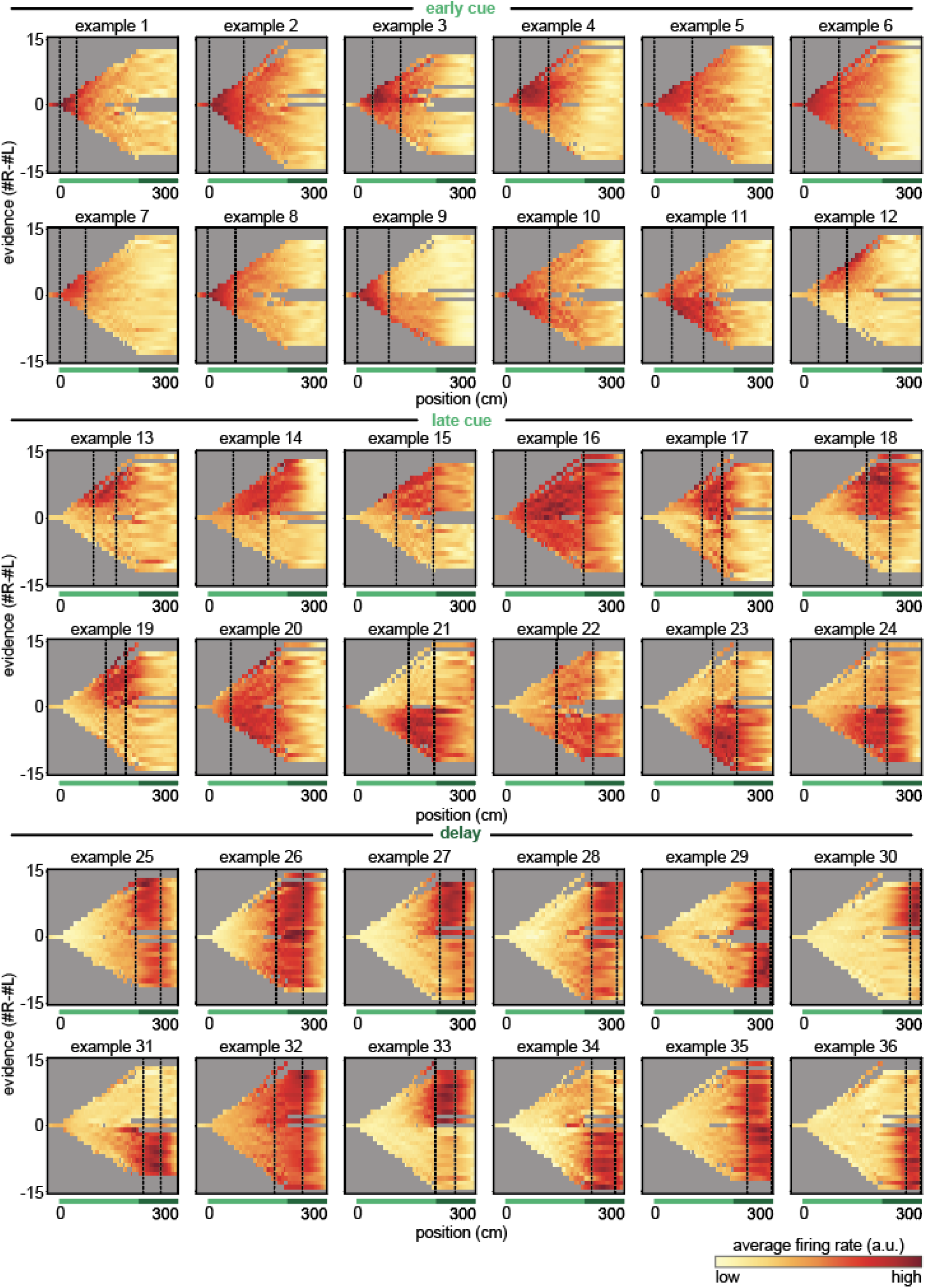
Example evidence-tuned neurons in ACC. Heatmaps of average normalized firing activity within position-by-evidence bins of 36 example individual neurons from ACC with significant evidence tuning (see Methods). Gray denotes position-evidence bins where there were fewer than 2 observations. A representative selection of neurons was made from neurons with mean position tuning in the early cue (0-100 cm), late cue (100-200 cm), and delay (200-300 cm) regions of the maze.

**Extended Data Figure 8.**
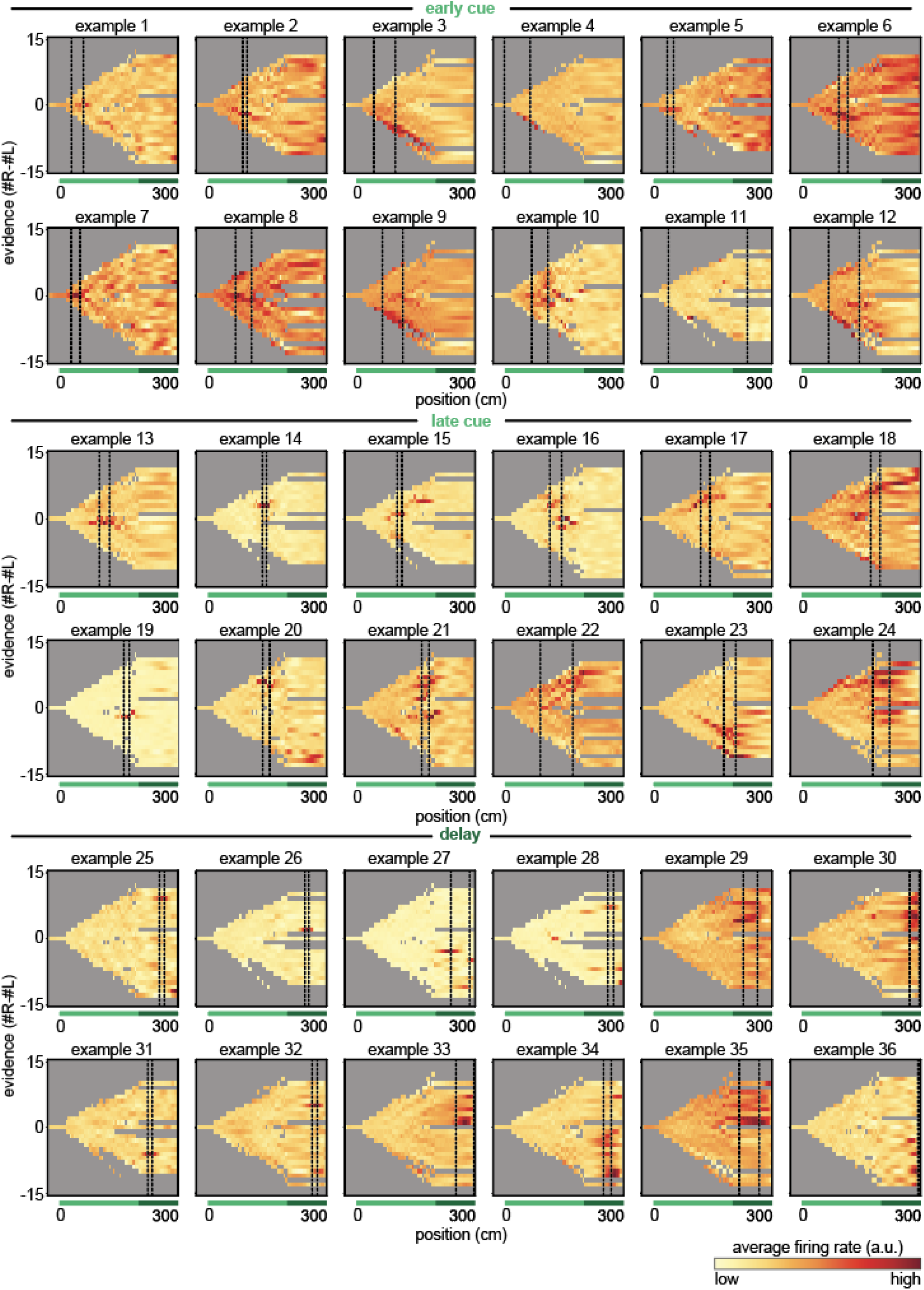
Example evidence-tuned neurons in HPC. Same as in Extended Data Fig. 7 but for neurons from HPC.

**Extended Data Figure 9.**
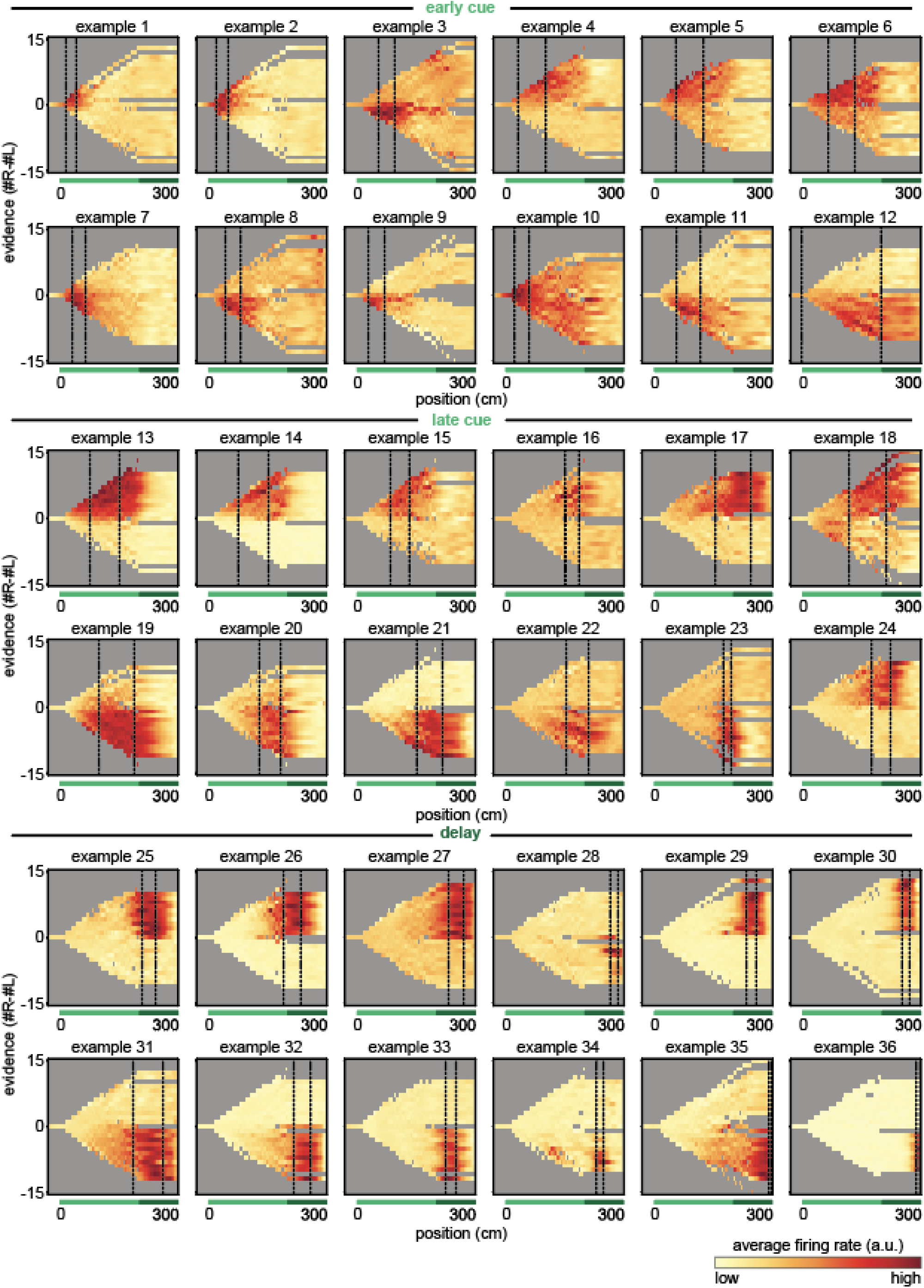
Example evidence-tuned neurons in RSC. Same as in Extended Data Fig. 7 but for neurons from RSC.

**Extended Data Figure 10.**
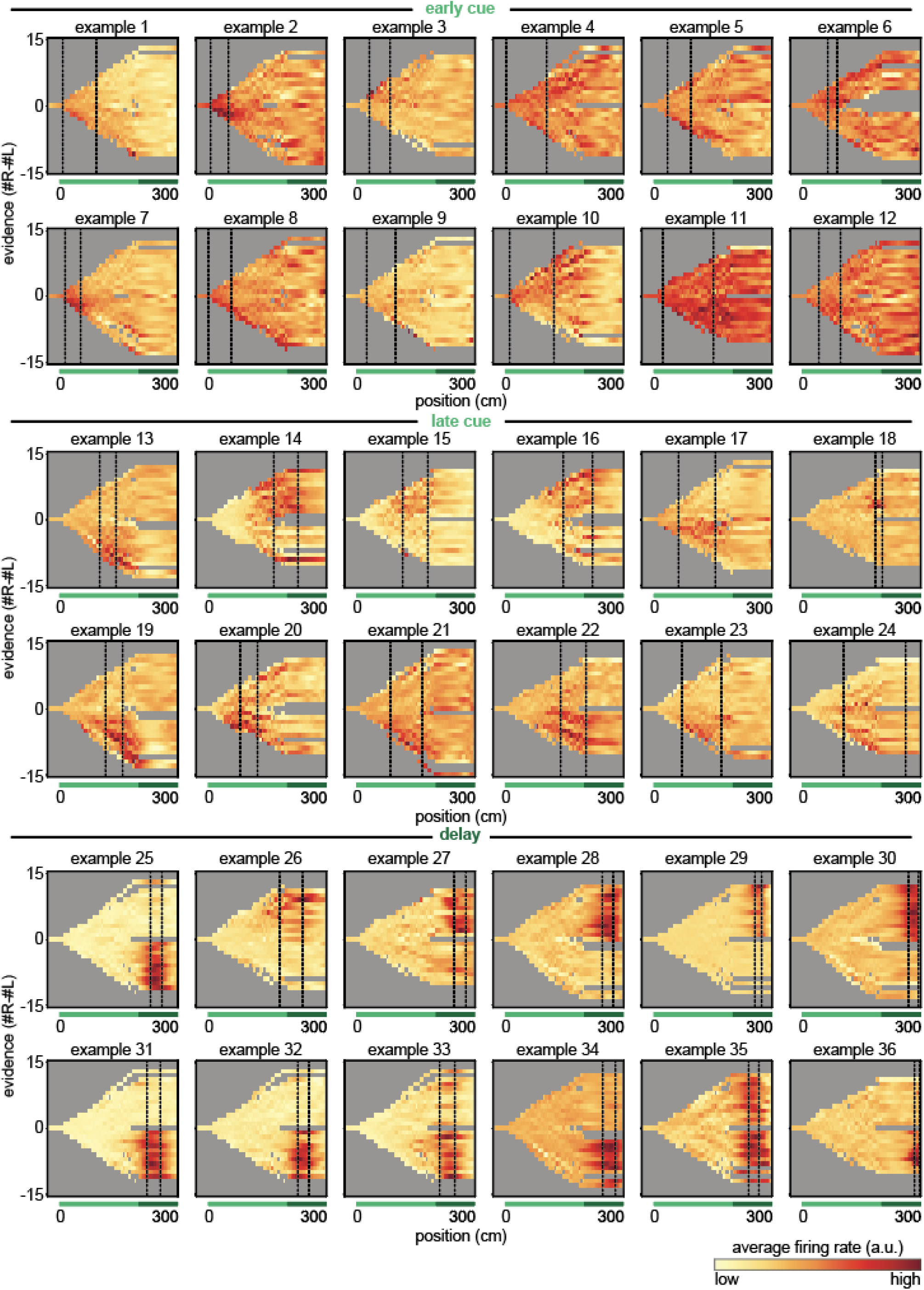
Example evidence-tuned neurons in DMS. Same as in Extended Data Fig. 7 but for neurons from DMS.

**Extended Data Figure 11.**
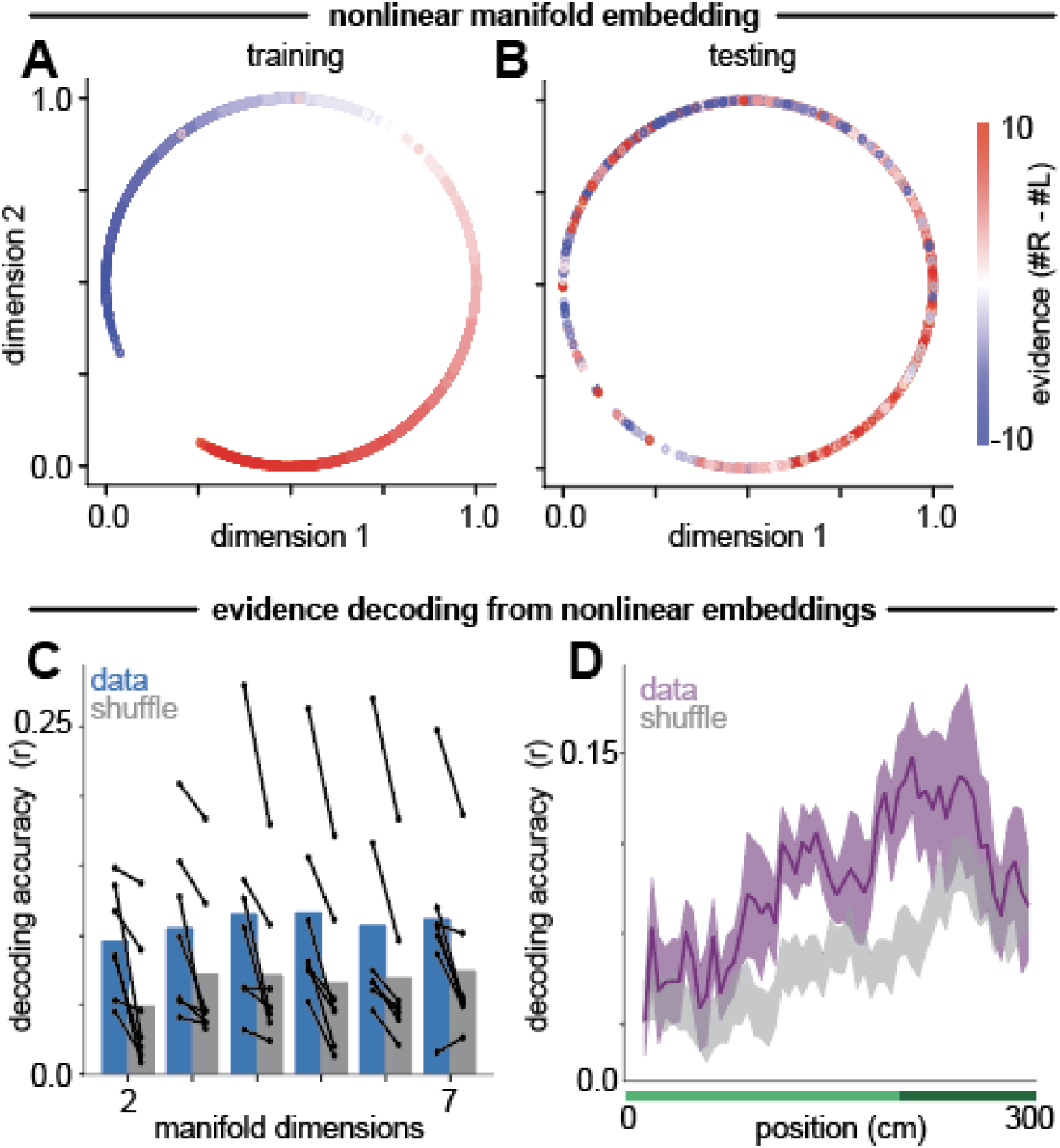
Decoding evidence from HPC populations. **(A)** CEBRA embeddings uncover a nonlinear mapping that smoothly captures evidence levels (indicated by the color of the points) in the training set. **(B)** Same as in (A) but for the test set. **(C)** A k-nearest neighbors decoder can decode evidence from the CEBRA embedding (average performance measured by Pearson’s correlation coefficient (r) between the actual and decoded evidence, blue) above a sign-of-evidence matched shuffle (gray). Lines indicate performance on individual sessions, compared to the corresponding shuffle. **(D)** Average decoding performance across different positions in the maze compared to a sign-of-evidence matched shuffle (gray) for 2-dimensional CEBRA embeddings.

**Extended Data Figure 12.**
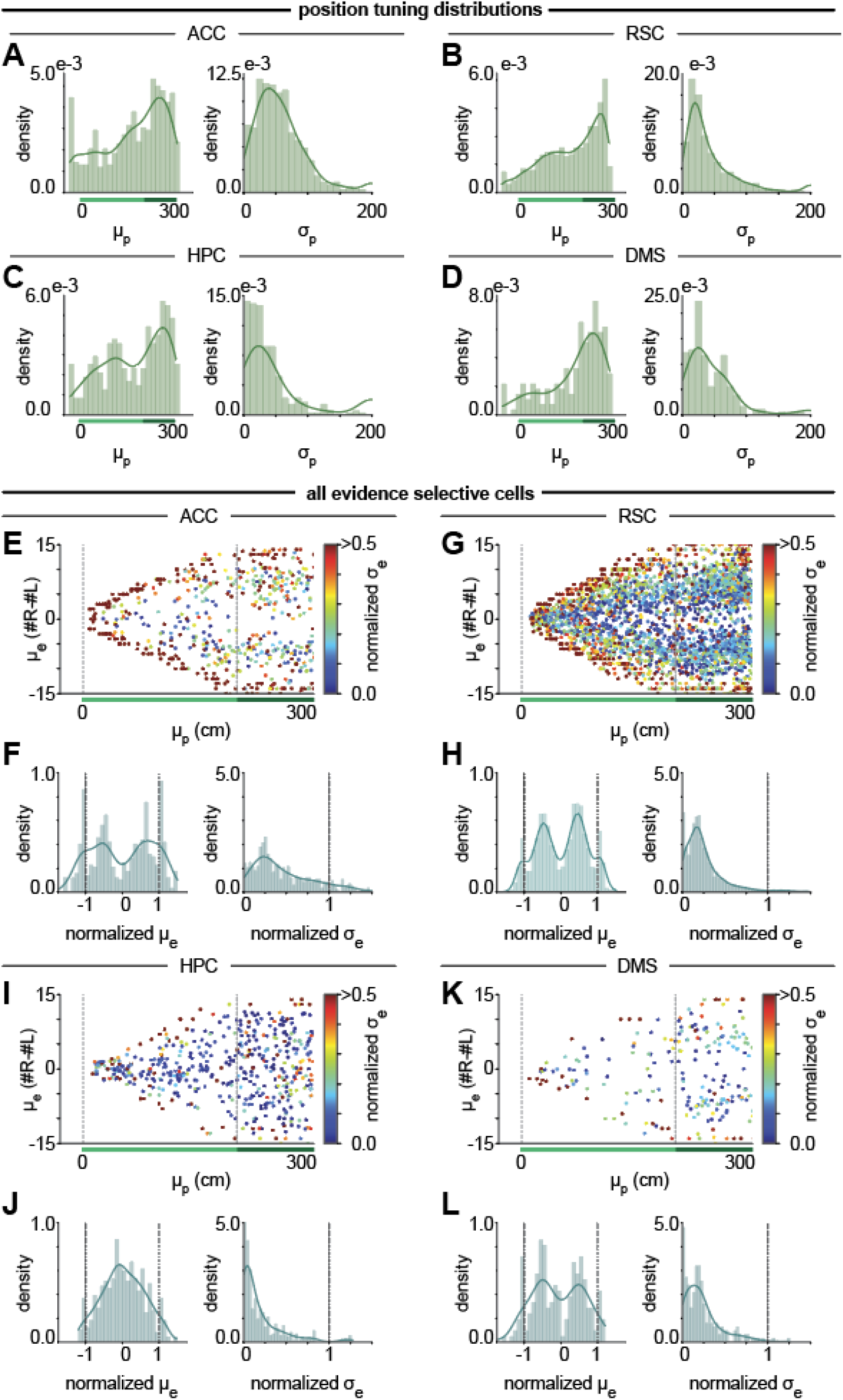
Supplementary distribution plots. **(A)** Left: Distribution of *μ*_*p*_ for the neurons in ACC plotted in Figure 5. Right: Distribution of *σ*_*p*_ for the same neurons. **(B-D)** Same as (A) but for neurons in RSC (B), HPC (C), and DMS (D), where DMS neurons are plotted in Extended Data Fig. 5. **(E)** Scatter of fit position mean vs. fit evidence mean colored by normalized *σ*_*e*_ for all significantly evidence-tuned neurons in ACC. **(F)** Left: Distribution of normalized *μ*_*e*_ for the neurons in E. Right: Distribution of normalized *σ*_*e*_ for the neurons in E. **(G-H)** Same as E-F but for RSC. Note that, by comparison to Figure 5, the narrowly tuned neurons in RSC deep blue tend to have poorer fits. **(I-J)** Same as E-F but for HPC. **(K-L)** Same as E-F but for DMS.

**Extended Data Figure 13.**
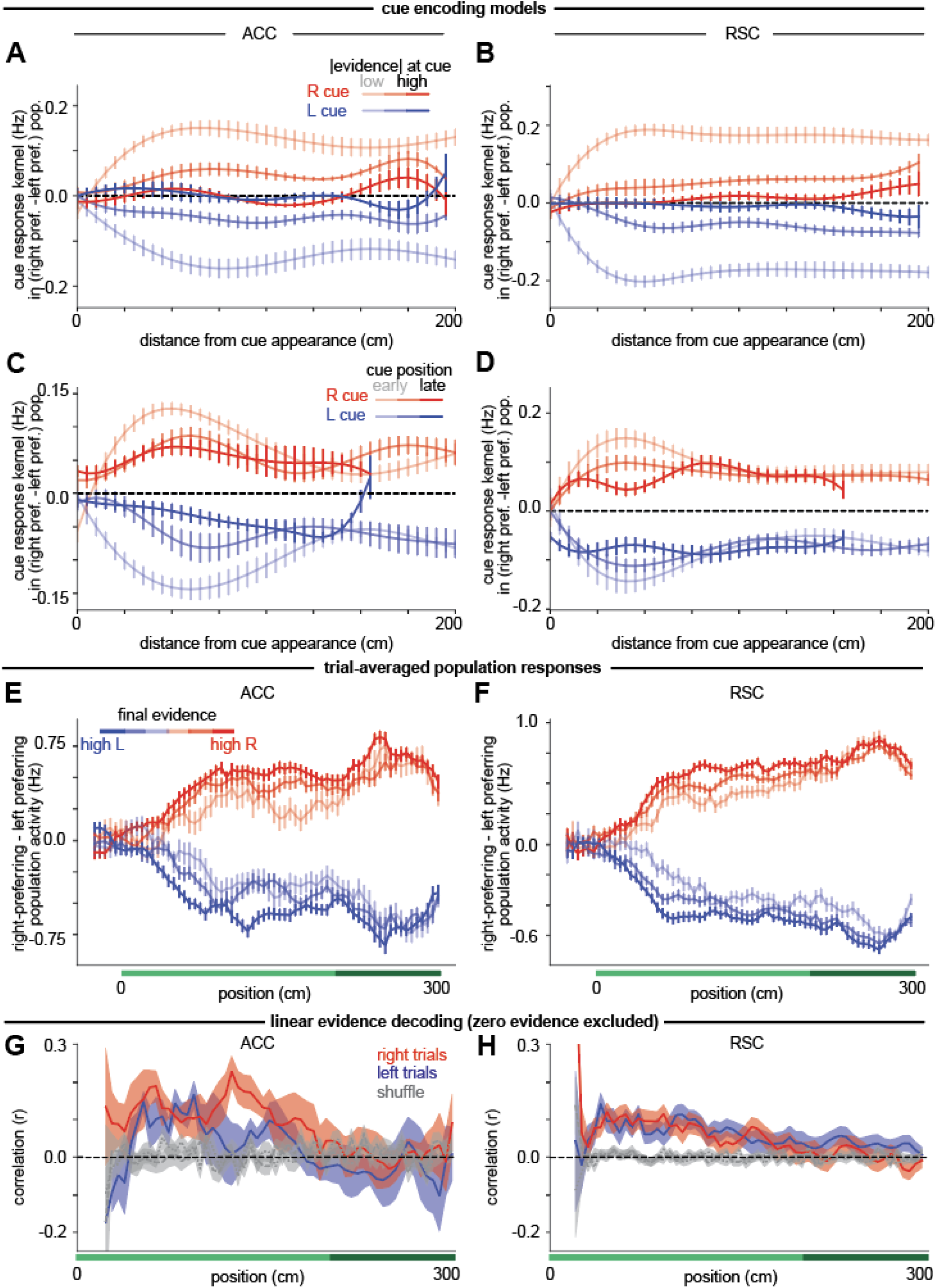
Single trial signatures of graded evidence accumulation. **(A)** Change in the difference in activity between the right and left populations in ACC following a left cue (blue) or right cue (red) when the current absolute value of evidence is low (light colors) to high (dark colors), where low is defined by |*e*| ≤ 1, medium by 2 ≤ |*e*| ≤ 4, or high by |*e*| ≥ 5. (Average r^2^ = 0.29 across sessions.) **(B)** Same as (A) but for RSC. (Average r^2^ = 0.21 across sessions.) **(C)** Change in the difference in activity between the right and left populations in ACC following a left cue (blue) or right cue (red) when the cue appears in the early (light colors) to late cue region (dark colors). (Average r^2^ = 0.27 across sessions.) **(D)** Same as (C) but for RSC. (Average r^2^ = 0.20 across sessions.) **(E)** For active cells in ACC with significant evidence coefficient in the single cell encoding analysis (see Methods), trial-averages of the difference between the mean activities of the left and right populations of for trials with different final evidence levels (indicated by the color bar). **(F)** Same as E but for RSC. **(G)** At each position, the cross-validated correlation between actual evidence and predicted evidence from a linear population decoder in ACC on correct right evidence (*e* > 0) trials (red) and left evidence (*e* < 0) trials (blue), compared to shuffle (gray). Error bars indicate s.e.m. across sessions. **(H)** Same as (G) but for RSC.

## SUPPLEMENTARY TEXT

In the following, we provide the mathematical analysis of our two classes of models. We show explicitly that, for the parameter conditions derived below, the models achieve the two fundamental operations of (i) evidence accumulation within a position and (ii) transfer of information between positions. We begin with an analysis of the competing chains models, for both the uncoupled competing chains models (Extended Data Fig. 2) and the mutually inhibiting competing chains models (Fig. 3), and conclude with the analysis of the position-gated bump attractor (Fig. 4) and planar bump attractor (Extended Data Fig. 3).

### COMPETING CHAINS MODELS

The competing chains models represent evidence in the difference of activity of two chains of neurons. In principle, these chains could be completely independent, with each chain integrating the cues only to its respective side, but this would result in chains that only increase in amplitude across the trial, inconsistent with observed data. Instead, we consider architectures with ipsilateral excitation and contralateral inhibition, where the contralateral inhibition comes either from external inputs or from the other chain. Specifically, we consider: (i) a model in which the chains are uncoupled but towers appearing to the ipsilateral side are excitatory and towers appearing to the contralateral side are inhibitory (the uncoupled competing chains model, Extended Data Fig. 2A) and (ii) a model in which each chain receives input only from the ipsilateral side and competes with the other chain through mutual inhibition within the same position (the mutually inhibiting competing chains model, Fig. 3A).

### Basic Features of Competing Chains Models

The neurons in each model can be represented by the same basic set of differential equations, where the firing rate of the *i*^th^ neuron in the left chain, *r*_*i,L*_, evolves according to

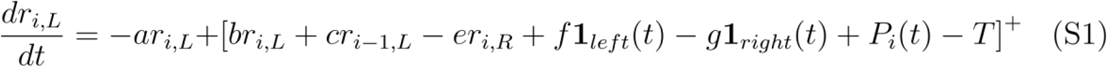

and similarly for the firing rate of neurons in the right chain. The various terms on the right side of the equation are described in the main text (see text below Eq. (1)), although here we add a term *g****1***_*right*_(*t*) that allows for inhibitory external inputs from cues on the contralateral side. The external cue inputs are given by ***1***_*left*_(*t*) for the left inputs and ***1***_*right*_(*t*) for the right inputs, which take the form

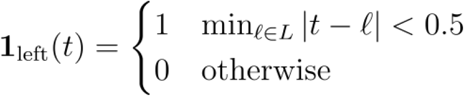

and

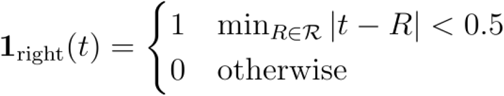

where *L* is the set of times of left cues and ℛ is the set of times of right cues.

The position signal is given by

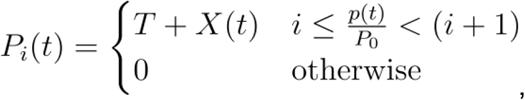

where *X*(*t*) ≥ 0 and *p*(*t*) is the position of the animal along the maze, so that each neuron receives an active position signal for length *P*_*0*_, and only one neuron in each chain receives an active signal at any time.

For both models, we require the chains to exhibit a number of properties in order to accurately accumulate evidence, so that at the end of a trial, the chain corresponding to the side with more cues will have greater amplitude. In particular, we look to achieve the following properties: (i) Each neuron is only active around a specific position, tiling space, with activity decaying away when the simulated animal leaves this position. (ii) External cue inputs are integrated linearly. (iii) In the absence of external cue inputs, when at a fixed position, the difference in neural activity of the two chains remains constant, without the sum growing unboundedly. (iv) In the absence of external cue inputs, the difference in amplitude of the two chains is preserved across positions. We analyze the necessary conditions on the model parameters to achieve each of these properties.

### Uncoupled Competing Chains Model

For the case of the uncoupled competing chains model (Extended Data Fig. 2), we take *e* = 0, giving us

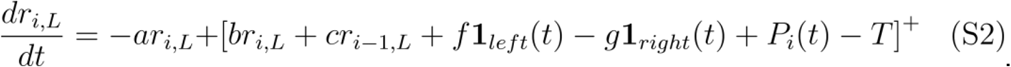

We consider below the conditions needed for this model to achieve the desired properties.

#### Neurons Active Only Around a Specific Position

Activity only occurring around a specific position when the simulated animal is navigating down the maze is achieved through the position-gating mechanism. We set the threshold *T* of the neuron sufficiently high that the term in square brackets will only affect the firing rate of the neuron in the presence of the position gating input *P*_*i*_(*t*). Otherwise, neural activity decays exponentially. This leads to the transient, sequential responses.

#### External Cues Integrated Linearly

In order for each cue to be integrated linearly, first, each cue should be weighted equally in its contribution to the final activity of the chain. This is achieved by giving each cue the same form of input in ***1***_*left*_(*t*) and ***1***_*right*_(*t*).

Second, this signal must be fully integrated by the chain. Since the external cue input term resides in the square brackets, only the neurons with the active position gating signal can integrate this signal. Without loss of generality, we consider the left-side chain. For the left-side neuron with an active position gating signal,

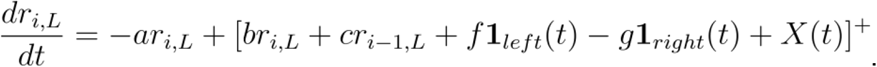

For this neuron to integrate its inputs, we need the term inside square brackets to be positive. We can see that this is guaranteed if *br*_*i,L*_ + *cr*_*i-1,L*_ is greater than *-g*. We show below that the difference in activity of the chains is preserved across positions in the absence of external cues when *b=c=a*, so that *r*_*i,L*_+*r*_*i-1,L*_ = *C* for some constant *C*. Thus, by initializing the activity at the start of the trial, *r*_*0,L*_, to be sufficiently large that the activity of the lower firing rate chain does not drop below *g/a*, these inputs will always be integrated.

We note that the argument above does not require the cues to have the square pulse specified here. Rather, it only requires that each cue has the same form.

#### Difference in Activity Between Chains Remains Constant at a Fixed Position

When the animal maintains a fixed position, the activity of the neurons at the preceding position decay away, so that in the absence of external cue inputs, we have

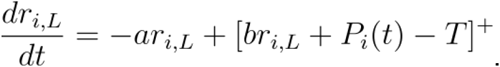

If we take *X*(*t*)=0, we have *P*_*i*_(*t*) *= T* at the active position, and thus

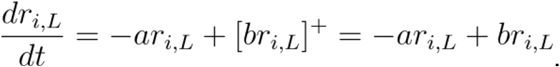

Thus, when *a = b*, each chain maintains constant activity, both preserving the difference between the chains and keeping the sum constant.

If instead *X*(*t*) *>* 0, this signal will be integrated, causing both chains to increase in amplitude. Such a signal could be designed to act as an urgency signal, but must be carefully crafted to prevent the activity in the chains from growing too rapidly.

#### Difference in Amplitude Preserved Across Positions

Preserving the amplitude of the chain across positions is a special case of preserving the difference across positions. To preserve amplitude across positions in the absence of cues, the next neuron in the chain must integrate the activity of the previous neuron at the same rate at which it decays. At the active position in the absence of external cue inputs, and assuming *X*(*t*)=0, we have

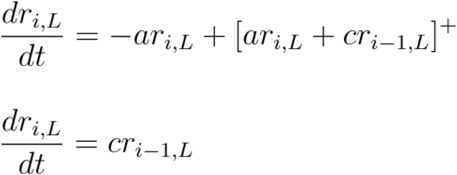

while we also have

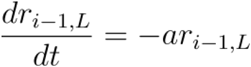

To preserve the total amplitude of activity,

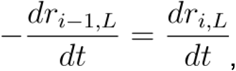

giving us the condition *a = c*.

We note that neuron *i* will only fully integrate the activity of the previous position in the infinite time limit. Due to the transient nature of the position-gating signal, which is active for the length of the position signal *P*_*0*_ or equivalently for time *P*_*0*_*/v* for an animal traveling at constant velocity *v*, we have that the final amplitude of the *i*^*th*^ neuron after the animal has traversed the length of the position gating signal will be

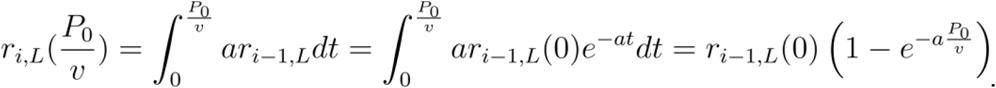

For *a* and *P*_*0*_ sufficiently large or *v* sufficiently small, the difference in amplitude between positions will be exponentially small.

Previous analysis of mouse running speed versus performance has shown little correlation within a session and a positive correlation on average across all sessions (see Supplementary Figure 7 in Pinto et al. (2018)^1^), suggesting that mice do not run at such a fast speed that information loss is problematic. Alternatively, additional connections could support the transfer of information from longer distances in the chain, increasing robustness to information loss.

#### Parameterization of the Uncoupled Competing Chains Model

In our simulations, in addition to taking *X*(*t*) = 0 Hz/s, we set *a* = 50 s^−1^, *b* = 50 s^−1^, *c* = 50 s^−1^, *f* = 50 s^−1^, *g* = 50 s^−1^, *P*_*0*_ = 20 cm, and *T* = 15000 Hz/s. Neurons *r*_*0,L*_ and *r*_*0,R*_ are initialized to 16.25 Hz activity level. We note that the neurons must be initialized to a sufficiently high non-zero activity level such that an external input to the contralateral side can be negatively integrated into the activity.

#### The Uncoupled Competing Chains Model with Saturation

In Extended Data Figure 2K, we present a version of the uncoupled chains model that has saturation in its firing. This was enforced by placing an upper bound on its firing rate. Note that because the dynamics are governed by the sum of an exponential decay and a non-negatively thresholded term, the firing rate of the neuron will be lower bounded at zero. To enforce the upper bound, we take

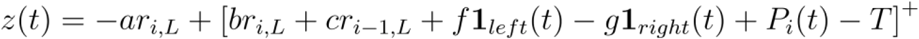

and

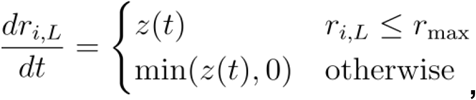

where *r*_*max*_ is the upper bound on the firing rate, which we set to 13 Hz in our simulations.to 13 Hz in our simulations.

### Mutually Inhibiting Competing Chains Model

In this model, the chains compete through mutual inhibition, rather than receiving opposing inputs. For the mutually inhibiting competing chains model, we take *g* = 0 in Eq. (S1), giving

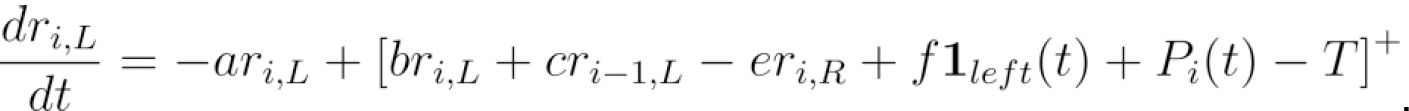

We next present to the necessary conditions for this model to achieve the desired properties.

#### Neurons Active Only Around a Specific Position

Activity only occurring around a specific position when the simulated animal is navigating down the maze is achieved through the same position gating mechanism as before, with *T* much larger than the other terms in square brackets so that the neuron will only be active when the position gate *P*_*i*_(*t*) is active.

#### External Cues Integrated Linearly

Each cue should be weighted equally in its contribution to the final activity of the chain. This is achieved by giving each cue the same form of input in ***1***_*left*_(*t*) and ***1***_*right*_(*t*). Moreover, the weights of mutual inhibition and self-excitation must be tuned such that the system of equations is a perfect integrator when the term in square brackets is above threshold. The position gating ensures that this can be true for at most two neurons, both at the same position. Specifically, when the term in square brackets is above threshold, we have

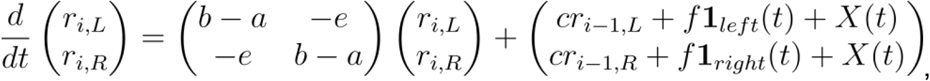

which has eigenvalues (*b-a-e*) for the common mode eigenvector, (1, 1), and (*b-a+e*) for the difference mode eigenvector, (1, -1). For a perfect integrator that accumulates evidence in the difference of firing between the two chains, we require that the eigenvalue corresponding to the difference mode eigenvector is zero, giving *e* = *a* - *b*. Due to the mutual inhibition architecture, the eigenvalue corresponding to the common mode (*b-a-e*) is negative (corresponding to decay). We note that, for the assumption that both neurons are above threshold to be true, we require *X*(*t*) *+ br*_*i,L*_ *+ cr*_*i-1,L*_ *- er*_*i,R*_ *>* 0 and *X*(*t*) *+ br*_*i,R*_ *+ cr*_*i-1,R*_ *- er*_*i,L*_ *>* 0. This can be achieved by setting *X*(*t*) *= I*_*ext*_ for some sufficiently large constant *I*_*ext*_, and integration will only be perfect in the range over which this is true. This common external input to both chains serves as a background about which integration occurs at each position in the chain.

#### Difference in Activity Between Chains Remains Constant at a Fixed Position

Holding the difference in activity between the neurons in the chain constant at a constant position is a natural consequence of having a perfect integrator for the difference mode eigenvector. Furthermore, because the common mode eigenvector is associated with a negative (decay-associated) eigenvalue, the sum of activity will not grow uncontrollably.

#### Difference in Amplitude Preserved Across Positions

To appropriately represent the difference in accumulated evidence between the chains, we require the difference *Δ*_*i*_ *= r*_*i,L*_ *- r*_*i,R*_ between the chains to be preserved between positions.

This difference evolves over time according to

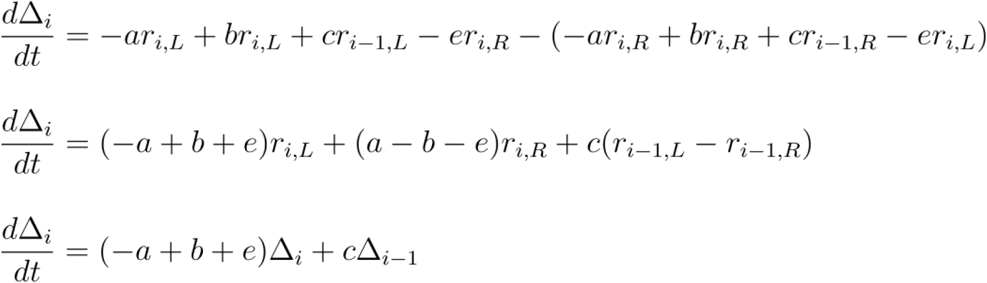

From our conditions on perfect integration, we have (*-a + b + e*) *=* 0, giving us

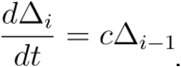

We also have that

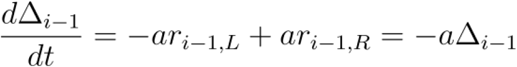

For the total difference in activity of the chains to be preserved, we require

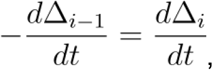

giving the condition *a = c*.

As in the case of the uncoupled chains model, due to the finite time of integration of the signal from the previous neuron in the chain, we have that

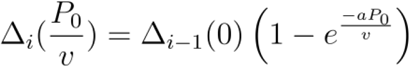

so that the difference between positions will be exponentially small for *a* and *P*_*0*_ sufficiently large or *v* sufficiently small.

### POSITION-GATED BUMP ATTRACTOR

Bump attractor based models have historically been used to model heading direction^2–13^ and path integration^14–16^. In this work, we modified the bump attractor model to be position-gated, so that within each position, there is a bump attractor that integrates visual cues. To accurately accumulate evidence, we require this model to exhibit several properties. Specifically, we show below that our model has the following properties: (i) Each neuron is only active around a specific position, tiling space, with the exact neurons that are active determined by the value of accumulated evidence. When the simulated animal leaves this position, activity decays away. (ii) In the absence of external inputs, when at a fixed position, the bump remains fixed. (iii) External cue inputs cause the bump to shift in the direction of the input, with the magnitude of the shift independent of the current bump location. (iv) In the absence of external cue inputs, the location of the bump along the evidence axis is maintained across positions.

Recall from Eq. (2) that our position-gated bump attractor dynamics for a neuron at position *i* and evidence *j* are governed by

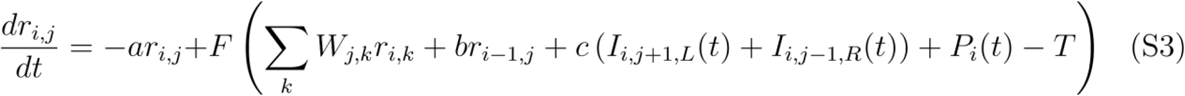

where

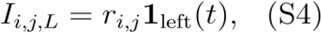

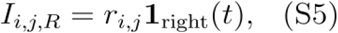

and where

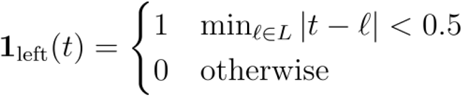

and

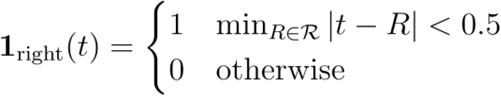

where *L* is the set of times of left cues and ℛ is the set of times of right cues. The various terms on the right side of the equation are described in the main text (see text below Eq. (2)). We use a sigmoidal nonlinearity

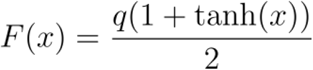

with *q > 0* and synaptic connection strengths governed by

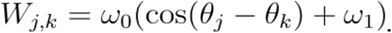

Although formally this synaptic connection matrix is circularly symmetric, we assume (and appropriately define parameters) such that we are working in a regime where the number of neurons is much larger than the space of observed evidence levels (see Methods). Thus, we do not encounter any effects from the circular boundary conditions. Alternatively, the tuning can be accomplished by adjusting the weights appropriately near the extremes of evidence^17,18^. Here, we use circular boundary conditions because this mathematical simplification facilitates the mathematical demonstration of the tuning conditions for the network provided below.

As in the competing chains models, the position signal is given by

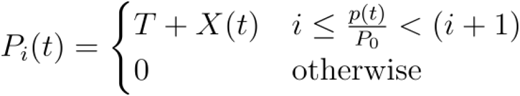

for *p*(*t*) the position of the animal along the maze at time *t*.

We consider below the conditions needed for this model to achieve the desired properties.

#### Neurons Active Only Around a Specific Position

Activity only occurring around a specific position when the simulated animal is navigating down the maze is achieved by the nonlinearity *F* saturating at 0 for large negative input. When the position gate *P*_*i*_(*t*) *≥ T*, the bump attractor becomes active in layer *i*. For all other layers, for *T* sufficiently large, the activities of all neurons decay temporally with time constant *1/a*, leading to sequential, transient responses.

#### Bump is Stable Within a Position in the Absence of Input

At the active position, *P*_*i*_(*t*) *≥ T* and we assume that the position-gating signal is constant, with *X*(*t*) = *X*_0_. Assume that the activity at position *i-*1 has decayed to be sufficiently small, and that there are no external cues. A stationary solution will satisfy

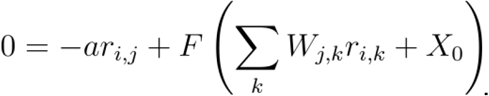

Since the entries of *W* depend only on | *j - k* |, any rigid translation of this stationary solution will also be stationary. Furthermore, these weights also guarantee that a stationary solution will be symmetric. A stationary solution centered at evidence *j* will correspond to a solution of the equations

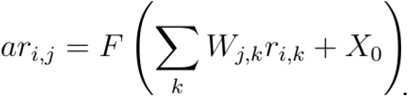

For ease of analysis, we approximate the discrete sum above by an integral. This is done to facilitate the analysis below of moving the bump along the direction of the attractor and strictly would correspond to the limit of having sufficiently close together evidence levels that they form a continuum, parameterized by *θ* at position *i*, with corresponding firing rates at each evidence level given by *r*(*θ*). In this continuum limit, at any given position, we have

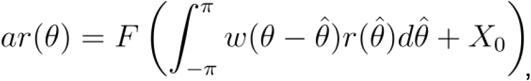

where

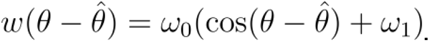

By the symmetry arguments above, we have that if *r*(*θ*) is a stationary solution, *r*(*θ+δ*) is also a stationary solution.

To examine the stability of this solution at the active position, we return to the original differential equation, in the continuum limit,

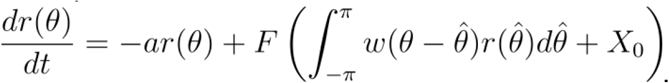

Taking the weighted integral of each side of the equation,

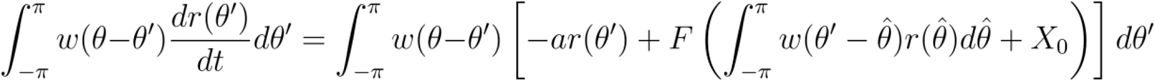

and defining

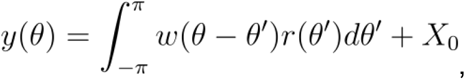

which can be interpreted as the total input current to the neuron whose tuning is centered at position *θ*, we have

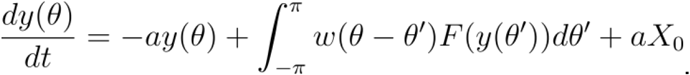

With this transformation, our equation is now in the same form as in the standard proofs that this stationary solution is a neutrally stable attractor, rather than an unstable solution. We do not repeat the proof here, and instead refer the reader to Kishimoto and Amari (1979)^19^.

We note that *X*_0_ along with the synaptic connectivity determines the width of the bump. For a given level of synaptic connectivity, increasing *X*_0_ will lift up the bump, making the super-threshold portion wider, and decreasing *X*_0_ will lower the bump and make it narrower. Kishimoto and Amari (1979)^19^*X*_0_ ^19^

#### External Cues Cause a Shift in the Bump

We next show that inputs to the model cause the bump to shift and that the shift is linear in the number of cues. Inputs to the model in Eq. (S3) are given by Eqs. (S4) and (S5).

In the continuous framework, we have input to *r*(*θ*) of the form

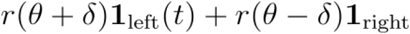

where *δ* is the magnitude of the asymmetric shift in the connections from the shifter neurons (orange and purple circles in Fig. 4A) to the evidence neurons.

We noted above that this model supports a bump of activity that is neutrally stable since all translations of this bump are also stationary. Neutral stability makes it such that inputs along the direction of the attractor cause a shift in the location of the bump, while inputs perpendicular to the attractor direction decay away without shifting the bump.

In the previous section, we showed that in the continuum limit, the attractor satisfies

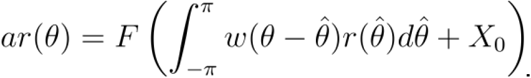

Define **r***\**(*θ*) to be the solution (i.e., vector of steady state neuronal firing rates) that is centered at *θ =* 0. We note that this should not be confused with the non-boldface *r*\*(*θ*) that occurs below, which we use to denote the firing rate of the single neuron that has preferred evidence level *θ* when the network is at its stationary solution. Without loss of generality, we consider inputs to the network when it is at the stationary solution **r***\**(*θ*), as inputs when the network is at other stationary solutions are equivalent, up to translation, by symmetry. The direction along the attractor is given by

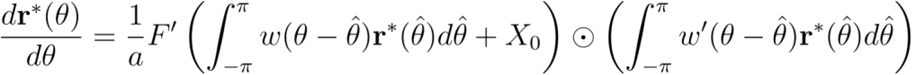

where ⊙ denotes the Hadamard product (i.e., element-wise multiplication), *F’* is the derivative of *F* with respect to its argument, and

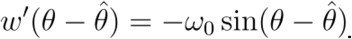

An input will cause a shift along the attractor if the projection of the input along this direction is nonzero. Since *F* is monotonically increasing, *F’(z) > 0* for all *z*, and *1/a* > 0, so for the projection to be nonzero, we require only that the projection of the input onto

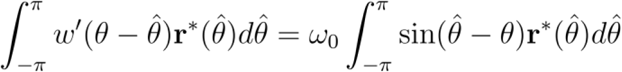

is nonzero. Since the inputs **r**\*(*θ - δ*) and **r**\*(*θ + δ*) do not arrive at the same time, we can consider each input component individually to show the projection is non-zero. For the **r**\*(*θ - δ*) component, we have

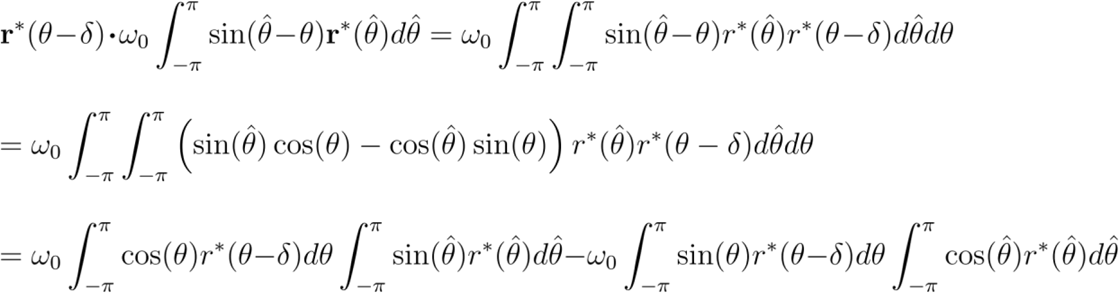

Since 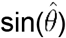 is an odd function and 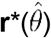 is an even function, the integral of their product will be zero, so the first term above disappears and we are left with:

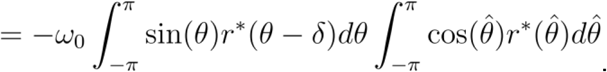

The second integral 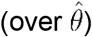 is nonzero and positive, since 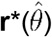 is even and, from the shape of the bump, has a nonzero fundamental frequency in its Fourier expansion. The first integral (over *θ*) is also nonzero for *δ* nonzero because sin(*θ*) is an odd function and, from the shape of the bump and since 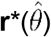 is even, the shifted bump **r**\*(*θ - δ*) has a nonzero odd fundamental frequency component. Together, this gives a nonzero projection of the input onto the direction of the attractor, so that the input causes a shift in the bump.

In the discrete case, the input additionally must be active for sufficiently long so as to move the location of the bump to the location of the next evidence level.

The above argument also shows that the shift in the bump will be linear in the number of cues. This is because the stationary solutions are invariant under circular shifts. This means that the input will have the same form for any stationary solution, and hence the same shift along the attractor, provided that there is sufficient time between inputs for the solution to converge to the attractor.

#### Bump Maintains the Evidence Location Across Positions

When the position gate moves to position *i*, neuron *r*_*i,j*_ of the active position layer receives input from the corresponding neuron *r*_*i-1,j*_ of the previously active layer. Denote the evidence level of the peak of the bump by *B*. Since the bump at position *i-*1 has its greatest activity at *B*, neuron *r*_*i,B*_ will have the greatest input. Since the bump of activity is a symmetrically shaped neutrally stable attractor, and the connectivity between layers *i-*1 and *i* is also symmetric, the bump at position *i* will also form a peak at *B*.

### PLANAR BUMP ATTRACTOR

In Extended Data Figure 3, we present a model that accumulates velocity signals along the position axis to create the shifts in the active position while evidence continues to be integrated along the evidence axis, giving a planar bump attractor. The activity *r*_*i,j*_ of a neuron at position *i* and evidence level *j* evolves according to

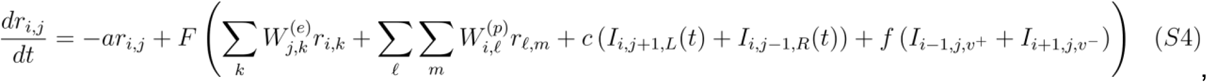

where as in the position-gated bump attractor, *a* is the exponential decay rate in the absence of input and *F* is the output nonlinearity. In this case, *W*^*(e)*^ is the matrix of synaptic connections between neurons at the same position but different evidence levels and *W*^*(p)*^ is the matrix of synaptic connections between neurons at different positions regardless of evidence level. *I*_*i,j+1,L*_ and *I*_*i,j-1,R*_ are the firing rates of the “shifter neurons” in evidence, which have synaptic connection strength *c. I*_*i-1,j,v*_ and *I*_*i+1,j,v*_ are the firing rates of the “shifter neurons” in position, which have synaptic connection strength *f*.

#### Parameterization of the Planar Bump Attractor

We took neurons at 17 different positions and 35 different evidence levels, for a total of 595 neurons. Each neuron was defined by an evidence angle, *θ*, and a position angle, *ϕ*. These angles were determined by assigning each evidence level an angle evenly spaced between 0 and 2*π*, and similarly assigning each position bin an angle evenly spaced between 0 and 2*π*. Along each axis, the strengths of synaptic connections were symmetric with excitatory connections between neurons with similar tuning on that axis and inhibitory between neurons farther apart in their tuning. Specifically, the entries of the matrix of synaptic connections along the evidence axis W^(e)^ for a neuron tuned to evidence *j* and a neuron tuned to evidence *k* are given by

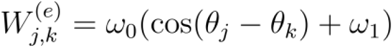

and the entries of the matrix of synaptic connections along the position axis W^(p)^ for a neuron tuned to position bin *i* and a neuron tuned to position bin *h* are given by

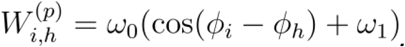

When an input is present, asymmetric connections result in a shift of the bump along the evidence axis in the direction of the input via a set of evidence shifter neurons,

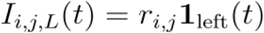

and

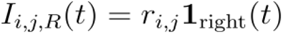

where

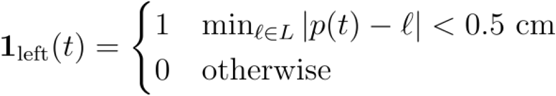

and

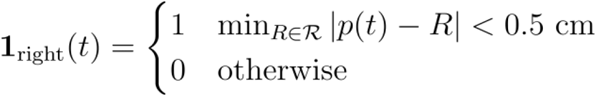

Similarly, asymmetric input connections with a set of velocity shifter neurons result in shifts along the position axis, based on the animal’s current velocity,

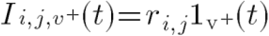

and

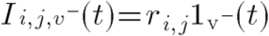

where

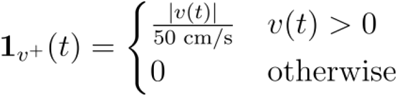

and

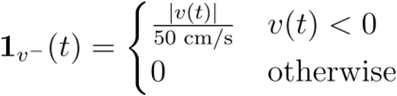

where *v*(*t*) is the animal’s velocity at time *t*.

The neuronal nonlinearity is defined by

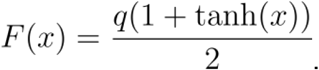

The parameters used in the simulations are: *a* = 55 s^−1^, *T* = 300 Hz/s, *q* = 1250 Hz/s, *ω*_*0*_ = 0.12, *ω*_*1*_ = -1.0, *c* = 0.2 s^−1^, *f* = 0.045 s^−1^. In Extended Data Fig. 3B,C, we used *v*(*t*) = 50 cm/s for the entire maze. Neurons *r*_*0,-1*_, *r*_*0,0*_, and *r*_*0,1*_ are initialized to 15 Hz, 17.5 Hz, and 15 Hz respectively, and all other neurons are initialized to 0 Hz activity. In Extended Data Fig. 3D, we used the same parameters but varied the velocity during different positions of the maze, taking

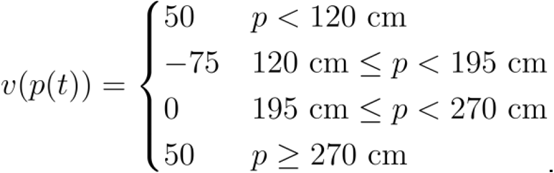

#### Properties of the Planar Bump Attractor

The same arguments as for the position-gated bump attractor show that the bump will be stable along the evidence axis and shift linearly with evidence inputs. Similar arguments show that the bump will be stable along the position axis and that velocity will enact shifts in the bump location due to the connectivity along the position axis being analogous to that along the evidence axis. As in the case of the position-gated bump attractor, the location along the evidence axis will be maintained across positions since the neuron at the same evidence level will have the greatest input. Analogous arguments show that position will be maintained when evidence levels shift.

